# FOSSIL CALIBRATIONS FOR THE ARTHROPOD TREE OF LIFE

**DOI:** 10.1101/044859

**Authors:** Joanna M. Wolfe, Allison C. Daley, David A. Legg, Gregory D. Edgecombe

## Abstract

Fossil age data and molecular sequences are increasingly combined to establish a timescale for the Tree of Life. Arthropods, as the most species-rich and morphologically disparate animal phylum, have received substantial attention, particularly with regard to questions such as the timing of habitat shifts (e.g. terrestrialisation), genome evolution (e.g. gene family duplication and functional evolution), origins of novel characters and behaviours (e.g. wings and flight, venom, silk), biogeography, rate of diversification (e.g. Cambrian explosion, insect coevolution with angiosperms, evolution of crab body plans), and the evolution of arthropod microbiomes. We present herein a series of rigorously vetted calibration fossils for arthropod evolutionary history, taking into account recently published guidelines for best practice in fossil calibration. These are restricted to Palaeozoic and Mesozoic fossils, no deeper than ordinal taxonomic level, nonetheless resulting in 80 fossil calibrations for 102 clades. This work is especially timely owing to the rapid growth of molecular sequence data and the fact that many included fossils have been described within the last five years. This contribution provides a resource for systematists and other biologists interested in deep-time questions in arthropod evolution.

**ABBREVIATIONS:**

AMNH: American Museum of Natural History
AMS: Australian Museum, Sydney
AUGD: University of Aberdeen
BGR: Bundesanstalt fur Geowissenschaften und Rohstoffe, Berlin
BMNH: The Natural History Museum, London
CNU: Key Laboratory of Insect Evolutionary & Environmental Change, Capital Normal University, Beijing
DE: Ulster Museum, Belfast
ED: Ibaraki University, Mito, Japan
FMNH: Field Museum of Natural History
GMCB: Geological Museum of China, Beijing
GSC: Geological Survey of Canada
IRNSB: Institut Royal des Sciences Naturelles de Belgique, Brussels
KSU: Kent State University
Ld: Musee Fleury, Lodeve, France
LWL: Landschaftsverband Westfalen-Lippe-Museum fur Naturkunde, Munster
MACN: Museo Argentino de Ciencias Naturales, Buenos Aires
MBA: Museum fur Naturkunde, Berlin
MCNA: Museo de Ciencias Naturales de Alava, Vitoria-Gasteiz, Alava, Spain
MCZ: Museum of Comparative Zoology, Harvard University
MGSB: Museo Geologico del Seminario de Barcelona
MN: Museu Nacional, Rio de Janeiro
MNHN: Museum national d'Histoire naturelle, Paris
NHMUK: The Natural History Museum, London
NIGP: Nanjing Institute of Geology and Palaeontology
NMS: National Museum of Scotland
OUM: Oxford University Museum of Natural History
PBM: Palaobotanik Munster
PIN: Paleontological Institute, Moscow
PRI: Paleontological Research Institution, Ithaca
ROM: Royal Ontario Museum
SAM: South Australian Museum, Adelaide
SM: Sedgwick Museum, University of Cambridge
SMNK: Staatliches Museum fur Naturkunde, Karlsruhe
SMNS: Staatliches Museum fur Naturkunde, Stuttgart
TsGM: F.N. Chernyshev Central Geologic Prospecting Research Museum, St. Petersburg
UB: University of Bonn
USNM: US National Museum of Natural History, Smithsonian Institution
UWGM: University of Wisconsin Geology Museum
YKLP: Yunnan Key Laboratory for Palaeobiology, Yunnan University
YPM: Yale Peabody Museum
ZPAL: Institute of Paleobiology, Polish Academy of Sciences, Warsaw.

## 1. Introduction

Accurate and precise systematic placement and dating of fossils underpins most efforts to infer a chronology for the Tree of Life. Arthropods, as a whole or in part, have received considerable focus owing to their incredible morphological disparity, species richness, and (relative to much of the Tree of Life) excellent fossil record. A growing number of recent studies have constructed timetrees for arthropods as a whole or for major groups therein (e.g. Bellec and Rabet, 2016; Bond et al., 2014; Bracken-Grissom et al., 2014, 2013; Djernæs et al., 2015; Fernandez et al., 2016, Fernandez et al., 2014a; Fernandez and Giribet, 2015; Garrison et al., 2016; Garwood et al., 2014; Giribet and Edgecombe, 2013; Herrera et al., 2015; Klopfstein et al., 2015; Legendre et al., 2015; Malm et al., 2013; McKenna et al., 2015; Misof et al., 2014; Oakley et al., 2013; Rehm et al., 2011; Schwentner et al., 2013; Song et al., 2015; Sun et al., 2015; Thomas et al., 2013; Tsang et al., 2014; Wahlberg et al., 2013; Wiegmann et al., 2011; Wood et al., 2013; Xu et al., 2015; Zhu et al., 2015). These studies vary in how well they have adhered to best practices for selecting calibration fossils, as many previous calibrations assume that fossil taxonomy accurately reflects phylogeny. Compounding the issue is the expansion of divergence time studies for a variety of comparative questions far beyond systematics and biogeography, including habitat shifts (Letsch et al., 2016; Lins et al., 2012; Rota-Stabelli et al., 2013a; Yang et al., 2013), genome evolution (Cao et al., 2013; Schwarz et al., 2014; Starrett et al., 2013; Wissler et al., 2013; Yuan et al., 2016), origins of novel characters and behaviours (Rainford et al., 2014; Sanggaard et al., 2014; Wheat and Wahlberg, 2013), evolution of parasites and disease (Ibarra-Cerdeña et al., 2014; Palopoli et al., 2014; Rees et al., 2014; Zhou et al., 2014), rate of diversification and its relationship to morphology and ecology (Lee et al., 2013; Wiens et al., 2015), coevolution (Kaltenpoth et al., 2014; Shelomi et al., 2016; Wilson et al., 2013), conservation (Owen et al., 2015), and the use of arthropods as a model for methodological development (O’Reilly et al., 2015; Ronquist et al., 2012; Warnock et al., 2012; Zhang et al., 2016).

Recent consensus on best practices for calibration fossil selection requires reference to specific fossil specimen(s), phylogenetic or morphological evidence justifying placement of the fossil, and stratigraphic and/or absolute dating information for the fossil (Parham et al., 2012). The importance of accurate phylogenetic knowledge of calibration fossils is underscored by recent controversies in dating the evolution of insects, where arguments hinge on the classification of particular ‘roachoid’ fossils on the stem lineage of Dictyoptera, with resulting differences on the order of 100 Myr (Kjer et al., 2015; Tong et al., 2015). With the explosion of taxonomic sampling in molecular phylogenies due to improvements in sequencing technology, improving the coverage of fossil calibrations is equally important. Recommendations include, for example, including as many as one fossil per ten extant OTUs for precise ages, with a varied distribution across lineages and clade depth (Bracken-Grissom et al., 2014). As a response, we have compiled an atlas of 80 rigorously scrutinized calibration fossils for 102 key nodes in arthropod phylogeny. These represent four basal ecdysozoan and arthropod clades, 17 chelicerates, 12 myriapods, 30 non-hexapod pancrustaceans, and 39 hexapod clades.

Where possible, we favour clade topologies resulting from a phylogenetic analysis of the largest total dataset. If phylogenomic analysis of genomes or transcriptomes has been performed but conflicts with morphology, a strongly supported molecular result is presented (e.g. putative clades such as Oligostraca that do not yet have identified morphological autapomorphies). If, however, molecular phylogenies have been constructed with few genes (e.g. clades such as Peracarida) or with highly conflicting results (e.g. Arachnida), morphological results are given greater weight. Where relevant, we discuss clade names with respect to NCBI’;s GenBank taxonomy (as recommended by the Fossil Calibrations Database: Polly et al., 2015), as this review is intended to be used by molecular biologists who are interested in dating the evolution of arthropod groups.

As there are >1.2 million species of arthropods, our calibrations are limited to fossils from the Palaeozoic and Mesozoic. Many extant clades have their oldest fossils in Cenozoic ambers such as the Eocene Baltic amber but are predicted to be vastly older based on fossils of allied taxa (e.g. Symphyla and Pauropoda relative to Chilopoda and Diplopoda).

While acknowledging the complexity of estimating an age prior for a fossil species known from multiple deposits, we use the oldest (e.g. section 28.3) and/or best known (e.g. section 51.3) horizons to provide minimum age constraints with the narrowest and most accurate age interval. Where there is substantial variation in age estimates for a fossil species, this issue is noted in the text. To accommodate the possibility of older fossils not yet discovered, we provide generous soft maxima (Ho and Phillips, 2009). Accordingly, when conducting divergence time analyses, prior distributions accounting for the large probability tail (e.g. gamma or lognormal) of an older undiscovered fossil may be appropriate. All fossil calibrations described herein are listed with their age information in Table A.1, formatted for adding age constraints in BEAST or PhyloBayes.

## 2. Crown Ecdysozoa

This clade comprises Euarthropoda, Onychophora (velvet worms), Tardigrada (water bears), Nematoida (itself comprising Nematoda and Nematomorpha), and Scalidophora (itself comprising Kinorhyncha, Loricifera, and Priapulida), their last common ancestor and all of its descendants (Fig. 1). Monophyly has been demonstrated on the basis of coding and non-coding molecular data (Campbell et al., 2011).

**Fig. 1.**
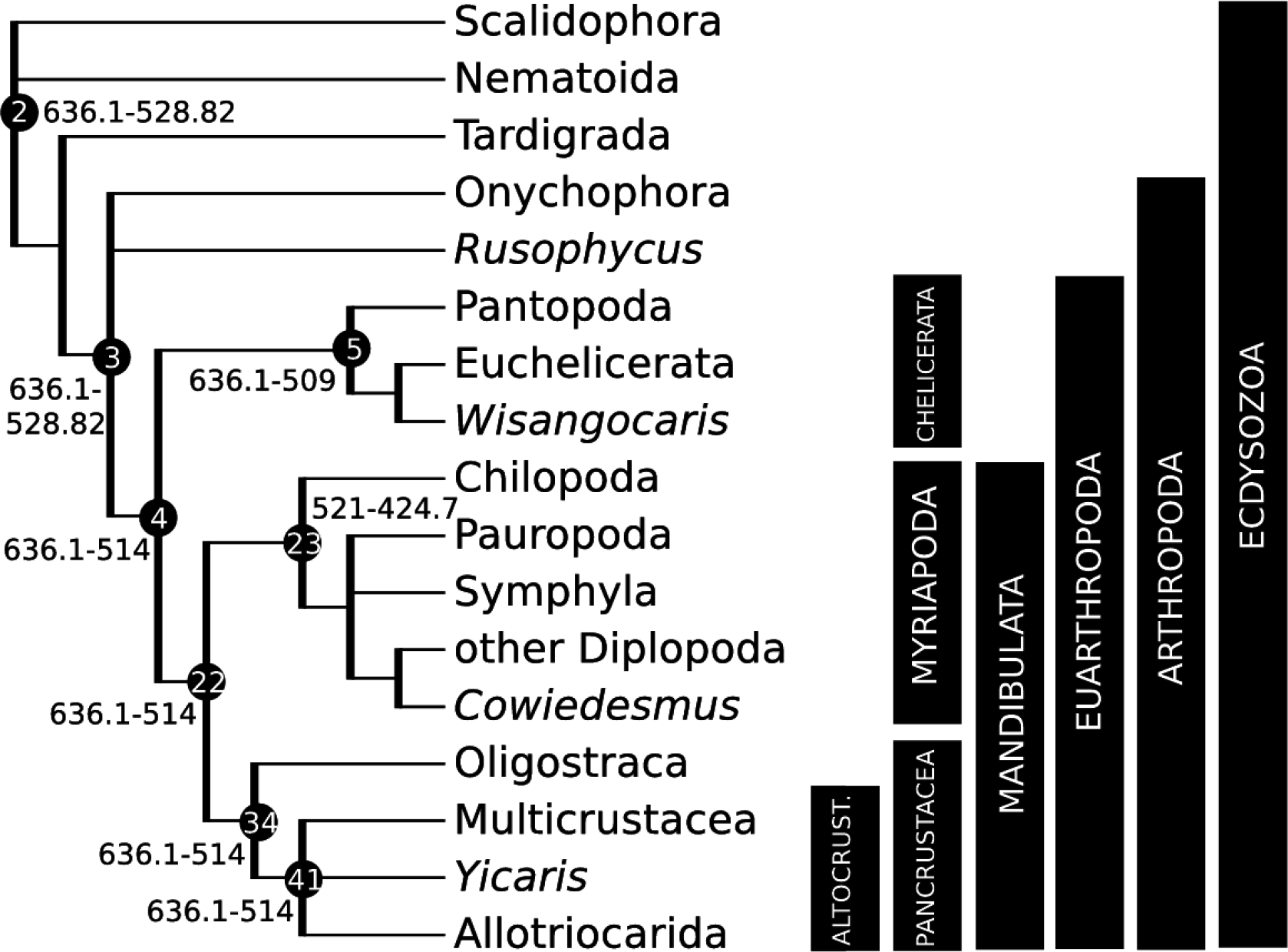
Calibration diagram for Ecdysozoa (nodes 2–5, 22–23, 34, 41). Altocrust. = Altocrustacea.

### 2.1. *Fossil specimens*

*Rusophycus* trace fossils in Member 2 of the Chapel Island Formation of the southwestern Burin Peninsula of southeastern Newfoundland, defining the base of the *R. avalonensis* Biozone. Arbitrarily, we fix this calibration on a specimen (Fig. 2a) from this unit figured by Narbonne et al. (1987: Fig. 6I; GSC 85983), as in Benton et al. (2015).

**Fig. 2.**
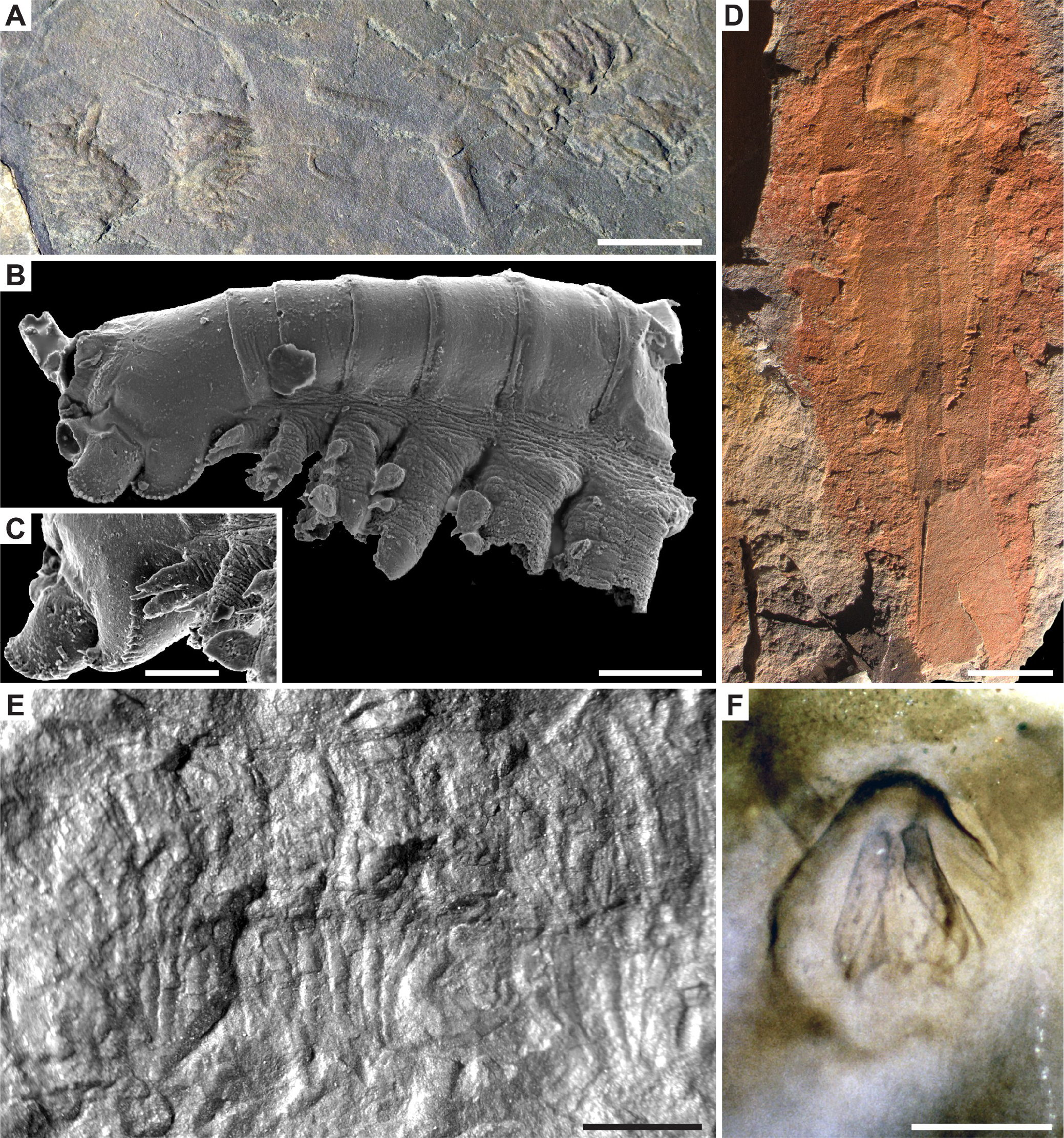
Major fossil calibrations for (A) nodes 2–3: *Rusophycus* trace fossils, GSC 85983, scale bar 20mm, image credit M. Coyne; (B-C) nodes 4, 22, 34, 41: *Yicaris dianensis,* YKLP 10840, scale bars 100μm, images credit X. Zhang; (B) whole specimen; (C) detail of epipodites; (D) node 5: *Wisangocaris barbarahardyae,* SAM P43679a, scale bar 5mm; (E) nodes 23–25, 27: *Cowiedesmus eroticopodus,* AMS F.64845, scale bar 2mm, image credit Y. Zhen; (F) nodes 64–65: *Rhyniella praecursor,* BMNH In.38228, scale bar 200μm, image credit NHMUK.

### 2.2. *Phylogenetic justification*

*Rusophycus* trace fossils are widely accepted to have been produced by arthropod-grade organisms, showing bilateral symmetry and evidence of segmented limbs used in their construction, the latter an apomorphy of Euarthropoda (Budd and Jensen, 2000).

### 2.3. *Age justification*

*Rusophycus* occurs well below the first animal body fossils in Cambrian sections around the world (Crimes, 1987; Crimes and Jiang, 1986; Goldring and Jensen, 1996; MacNaughton and Narbonne, 1999; Weber and Zhu, 2003). In many of these regions, records of *Rusophycus* begin with proximity to the base of the Cambrian. However, their ages are only well constrained in sections in Newfoundland, Canada, and Yunnan, China. Of these, records of *Rusophycus* begin low in Member 2 of the Chapel Island Formation of the southwestern Burin Peninsula of southeastern Newfoundland, defining the base of the *R. avalonensis* Biozone (Narbonne et al., 1987). The Biozone is itself dated through correlations to a section in New Brunswick where the younger ash bed has been dated by U-Pb series to 530.02 Ma ± 1.2 Myr (Isachsen et al., 1994; Peng et al., 2012), thus providing for a minimum constraint of 528.82 Ma.

A soft maximum constraint is based on that used by Benton et al. (2015), the maximum age interpretation of the Lantian Biota (Yuan et al., 2011). This, together with the Doushantuo Biota (Yuan et al., 2002), provides a series of Konservat-Lagerstätten preserving the biota in Orsten-and Burgess Shale-like modes of fossilization. None of these Lagerstätten, least of all the Lantian, preserves anything that could possibly be interpreted as even a total group eumetazoan and on this basis we define our soft maximum constraint at 635.5 Ma ± 0.6 Myr (Condon et al., 2005) and, thus, 636.1 Ma.

## 3. Crown Arthropoda

This clade comprises Euarthropoda and Onychophora *(sensu* Ortega-Hernandez, 2016), their last common ancestor and all of its descendants (Fig. 1). This clade has traditionally been called Panarthropoda (Nielsen, 1995), however, this latter name is most often used to refer to a group encompassing Arthropoda and Tardigrada, but we exclude tardigrades from our current grouping. Monophyly of this clade has been established through phylogenetic analysis of both non-coding and protein-coding gene datasets (Campbell et al., 2011), and morphological data sets (Legg et al., 2013), although it has been challenged by other recent morphological analyses that endorsed a rival sister group relationship between Euarthropoda and Tardigrada (e.g. Smith and Ortega-Hernandez, 2014). Note the name Arthropoda in GenBank refers to what we consider Euarthropoda; there is no GenBank taxonomy ID for the clade comprising Euarthropoda and Onychophora.

### 3.1. *Fossil specimens*

As for 2.1.

### 3.2. *Phylogenetic justification*

As for 2.2.

### 3.3. *Age justification*

As for 2.3.

## 4. Crown Euarthropoda

This clade comprises Chelicerata, Myriapoda and Pancrustacea, their last common ancestor and all of its descendants (Fig. 1). The monophyly of Euarthropoda, comprising the sister clades Chelicerata and Mandibulata (itself comprising Myriapoda and Pancrustacea) has been established on the basis of protein-coding and noncoding molecular data, as well as morphological data (Rota-Stabelli et al., 2011). Note that in Benton et al. (2015) this node was named Arthropoda (likely with reference to GenBank taxonomy). Here we follow the naming conventions outlined by Ortega-Hernandez (2016).

### 4.1. *Fossil specimens*

*Yicaris dianensis* Zhang et al., 2007. YKLP 10840, holotype (Fig. 2b,c), consisting of an almost complete articulated specimen (Zhang et al., 2007).

### 4.2. *Phylogenetic justification*

Several phylogenetic analyses of morphology (Edgecombe, 2010; Legg et al., 2013; Wolfe and Hegna, 2014) and morphology plus molecules (Oakley et al., 2013) place *Y. dianensis* within the crown group of Pancrustacea. Key characters include the presence of epipodites on the thoracic limbs; paddle-shaped mandibular and maxillary exopods; and the protopodite of post-mandibular limbs elongated as soft, setiferous endites. Any position supported within the crown group of Pancrustacea is within crown Euarthropoda.

### 4.3. *Age justification*

*Y. dianensis* was recovered from the Yu’anshan Formation at Xiaotan section, Yongshan, Yunnan Province, attributed to the *Eoredlichia-Wutingaspis* Biozone (Zhang et al., 2007). Chinese Cambrian stratigraphy has been revised substantially and the *Eoredlichia-Wutingaspis* Biozone is no longer recognized (Peng, 2009, 2003). However, *Eoredlichia* is known to co-occur with *Hupeidiscus,* which is diagnostic of the *Hupeidiscus-Sinodiscus* Biozone, which is formally recognised as the second biozone of the Nangaoan Stage of the Qiandongian Series of the Cambrian of China (Peng and Babcock, 2008). The Nangaoan is the proposed third stage of the Cambrian System for the International Geologic Timescale (Peng et al., 2012). Thus, a minimum constraint can be established on the age of the top of the Nangaoan, which has been dated to 514 Ma (Peng et al., 2012; Peng and Babcock, 2008).

Soft maximum as for 2.3.

### 4.4. *Discussion*

There are older records of euarthropods than *Y. dianensis*, notably trilobites, but their phylogenetic position within Arthropoda is contested (it is unclear whether trilobites are stem-Euarthropoda, stem-Chelicerata or stem-Mandibulata). *Wujicaris muelleri* Zhang et al., 2010 has an equal claim to being the earliest record of Arthropoda, but it is of equal age to the holotype of *Y. dianensis*.

## 5. Crown Chelicerata

This clade comprises Pantopoda (sea spiders) and Euchelicerata, their last common ancestor and all of its descendants (Figs. 1, 3 and 4). Monophyly of this clade has been established by phylogenetic analysis of nuclear protein-coding genes (Regier et al., 2010), transcriptomes (Meusemann et al., 2010; Rota-Stabelli et al., 2011), and morphology (Legg et al., 2013).

**Fig. 3.**
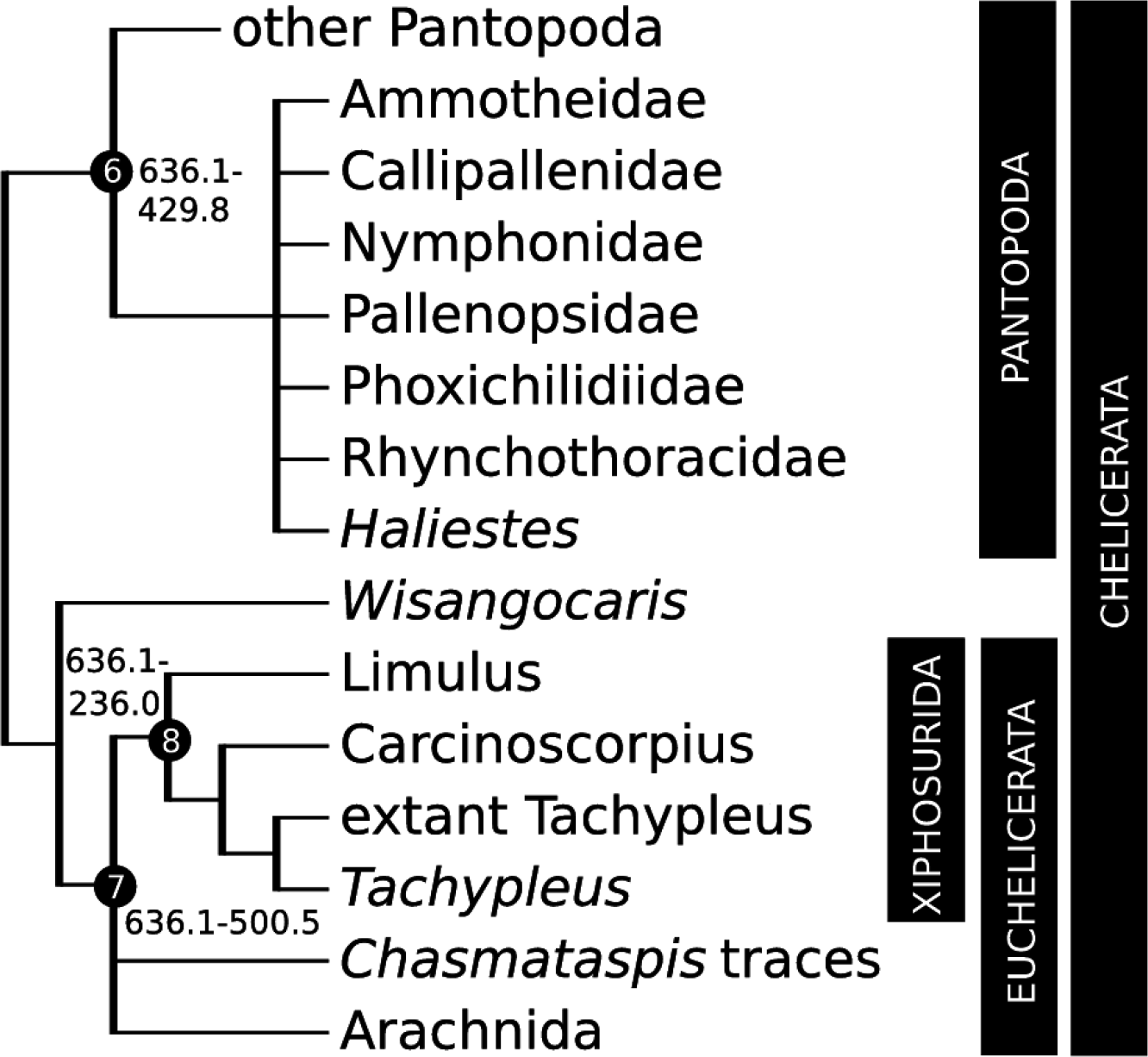
Calibration diagram for non-arachnid chelicerata (nodes 6–8).

**Fig. 4.**
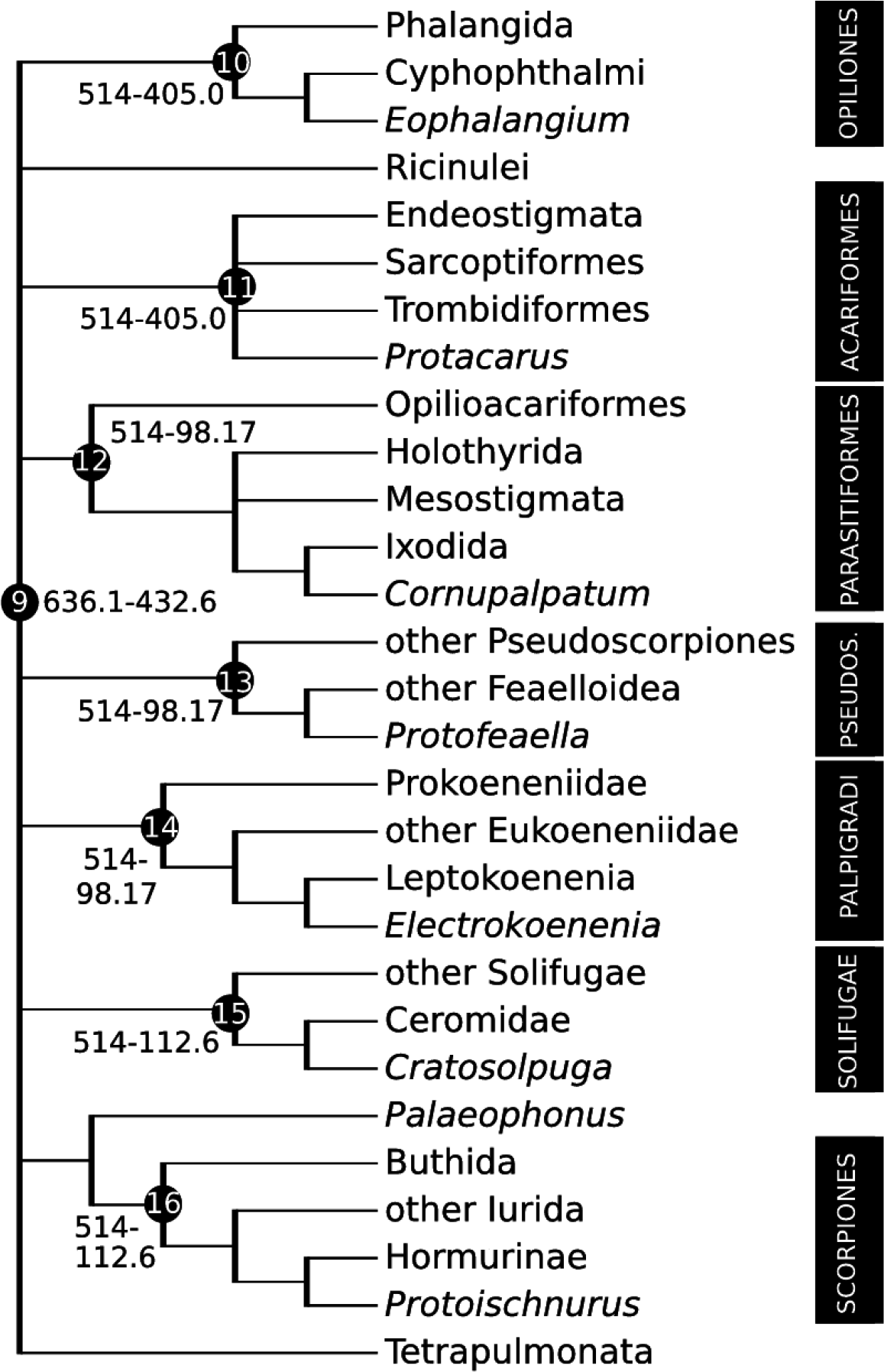
Box initially at rest on sled sliding across ice.Calibration diagram for non-tetrapulmonate Arachnida (nodes 9–16). Pseudos. = Pseudoscorpiones.

### 5.1. *Fossil specimens*

*Wisangocaris barbarahardyae* Jago, Garria-Bellido and Gehling, 2016. SAM P45427, holotype, almost complete specimen (Fig. 2d).

### 5.2. *Phylogenetic justification*

Few recent phylogenetic studies have addressed the stem-lineage of Euchelicerata (notable exceptions being Lamsdell, 2013; Legg, 2014; Legg et al., 2013). Including *W. barbarahardyae* in the dataset of Legg (2014), this species was resolved in most shortest cladograms as sister taxon to the middle Cambrian *Sanctacaris* and *Sidneyia,* and in all shortest cladograms as more closely related to Euchelicerata than Pantopoda, i.e. as crown group Chelicerata (Jago et al., 2016). This relationship is supported by the shared presence of pediform cephalic exites, multi-partite trunk exites, and a trunk composed of a posterior limb-less abdomen in both crown euchelicerates and the Cambrian taxa.

### 5.3. *Age justification*

*W. barbarhardyae* was collected from the Emu Bay Shale on Kangaroo Island, South Australia. Trilobite biostratigraphy correlates this unit with the upper part of the *Pararaia janeae* Zone in mainland South Australia (Jell in Bengtson et al., 1990; Fig. 2 in Jago et al., 2012), equivalent to the Canglangpuan Stage in South China and the late Botoman in Siberia (Gehling et al., 2011, Fig. 9). This dates the Emu Bay Shale to Cambrian Series 2, Stage 4, providing a minimum constraint of 509 Ma.

Soft maximum as for 2.3.

### 5.4. *Discussion*

Until recently the oldest evidence of chelicerates in the fossil record were thought to be represented by *Chasmataspis-like* trace fossils from the Furongian of Texas (Dunlop et al., 2004), and a putative pycnogonid larva from the Furongian of Sweden (Waloszek and Dunlop, 2002). However, in a number of recent phylogenetic analyses (e.g. Legg, 2014; Legg et al., 2013), a number of taxa from the middle Cambrian Burgess Shale Formation, namely *Sanctacaris uncata* Briggs and Collins, 1988, *Sarotrocercus oblita* Whittington, 1981, and *Sidneyia inexpectans* Walcott, 1911, have been resolved as stem-lineage representatives of Euchelicerata. These relationships are preserved with the addition of the older *W. barbarahardyae* to the dataset (Jago et al., 2016). Although another purported species of *Sidneyia (S. sinica* Zhang et al., 2002) is known from the older Chengjiang biota, it lacks many diagnostic features of this genus, and could therefore not be reliably used for calibration purposes. It should also be noted that *Sidneyia* only resolved as a stem representative of Euchelicerata under some iterations of the analysed data set of Legg (2014), specifically only when all characters were weighted equally, and therefore its exact phylogenetic position is equivocal.

## 6. Crown Pantopoda

This clade comprises Ammotheidae, Austrodecidae, Callipallenidae, Colossendeidae, Endeididae, Nymphonidae, Pallenopsidae, Phoxichilidiidae, Pycnogonidae and Rhynchothoracidae, their last common ancestor and all of its descendants (Fig. 3). Phylogenetic analyses of protein-coding genes (Arabi et al., 2010) or protein-coding genes combined with morphology (Arango and Wheeler, 2007) indicate monophyly.

### 6.1. *Fossil specimens*

*Haliestes dasos* Siveter et al., 2004. OUM C.29571, holotype (Fig. 5a). As the reconstruction of Herefordshire fossils requires serial grinding and photography of 20 μm sections (Sutton et al., 2002), the holotype figured in Siveter et al. (2004) and herein was thus destroyed in preparation. Morphological data for Herefordshire fossils are published as 3D models of the thin section photographs.

**Fig. 5.**
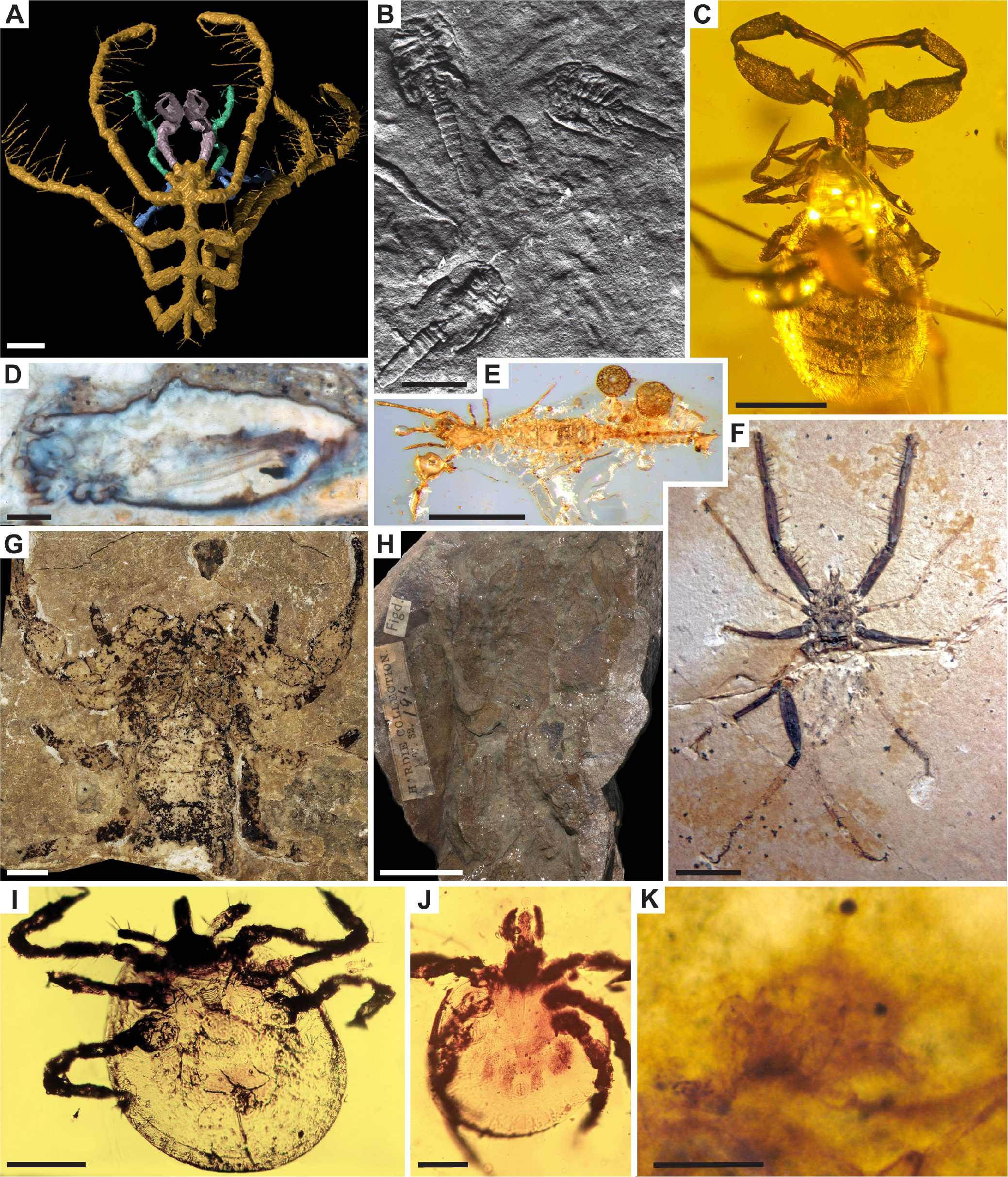
Chelicerate fossil calibrations for (A) node 6: *Haliestes dasos,* OUM C.29571, scale bar 500pm, image credit D. Siveter, M. Sutton, D. Briggs & D. Siveter; (B) node 7: *Chasmataspis-like* resting traces, MBA 1084, scale bar 20mm, image credit J. Dunlop; (C) node 13: *Protofeaella peetersae,* NHMII3115, scale bar 500pm, image credit H. Henderickx; (D) node 10: *Eophalangium sheari,* PBM 3503, scale bar 1mm, image credit J. Dunlop; (E) node 14: *Electrokoenenia yaksha,* NIGP 163263, scale bar 500pm, image credit M. Engel; (F) node 15: *Cratosolpuga wunderlichi,* SMNK 1268PAL, scale bar 5mm, image credit P. Selden; (G) node 9: *Eramoscorpius brucensis,* holotype ROM 5324, scale bar 10mm, image credit D. Rudkin & J. Waddington; (H) node 9: *Palaeophonus loudonensis,* NMS 1897.122.196, scale bar 20mm, image credit: J. Lamsdell; (I-J) node 12: *Cornupalpatum burmanicum,* scale bars 100 pm, image credit G. Poinar; (I) Holotype A-10–160; (J) Paratype A-10– 261; (K) node 11: *Protacarus crani,* BMNH In.24665, scale bar 100pm, image credit NHMUK.

### 6.2. *Phylogenetic justification*

Arango and Wheeler (2007) resolved *H. dasos* as sister to part of Ammotheidae (nested within clade of Ammotheidae, Callipallenidae, Nymphonidae, Pallenopsidae, Phoxichilidiidae, Rhynchothoracidae), i.e. as crown-group Pantopoda. *H. dasos* was classified separately from extant pycnogonids (Pantopoda) as an Order Nectopantopoda by Bamber (2007), although without explicit phylogenetic justification. It should be noted that *H. dasos* was included in the phylogenetic analysis of Legg et al. (2013), and resolved as sister-taxon to *Palaeopantopus,* which in turn resolved as sister-taxon to extant pycnogonids, however, just three extant exemplars were included in this study and as this was not extensive enough to determine the exact position of these fossil taxa with respect to crown-group exemplars, we continue to follow Arango and Wheeler (2007) in their placement.

### 6.3. *Age justification*

This fossil is preserved as a carbonate concretion from the volcaniclastic Herefordshire Lagerstatte of Herefordshire, England, at the Sheinwoodian-Homerian stage boundary, within the Early Silurian Wenlock Series (Siveter, 2008). As the Homerian begins at 430.5 Ma ± 0.7 Myr, a minimum age constraint for the Herefordshire can thus be placed at 429.8 Ma.

Soft maximum as for 2.3.

### 6.4. *Discussion*

Although *H. dasos* is the oldest assignable crown group pycnogonid, there is an older fossil, *Cambropycnogon klausmuelleri* Waloszek and Dunlop, 2002, from the Cambrian Orsten biota (minimally 497 Ma). *C. klausmuelleri,* however, is known only from larval stages, and does not share specific apomorphies with any extant larva. Without such characters, it is not possible to adequately confirm crown group affinity. Another fossil species, *Palaeomarachne granulata* Rudkin et al., 2013 from the Upper Ordovician of Manitoba, is specifically noted as a stem pantopod due to its likely plesiomorphic head tagmosis.

## 7. Crown Euchelicerata

This clade comprises Xiphosurida (horseshoe crabs) and Arachnida, their last common ancestor and all of its descendants (Fig. 3). Monophyly is established on the basis of phylogenetic analysis of transcriptomes (Rota-Stabelli et al., 2011; Sharma et al., 2014) and morphology (Legg et al., 2013). Note that monophyly of Euchelicerata is challenged by a recent morphological phylogeny, a result attributed to outgroup sampling (Garwood and Dunlop, 2014). Euchelicerata is not recognized in GenBank taxonomy.

### 7.1. *Fossil specimens*

*Chasmataspis-like* resting trace fossils (MBA 1084), Fig 5b. Described and illustrated in Dunlop et al. (2004, Figs. 9 and 10).

### 7.2. *Phylogenetic justification*

The assignment of the traces to Chasmataspida is based on impressions of plate-like opisthosomal opercula, one of the characters used to define Euchelicerata (Dunlop et al., 2004) Furthermore, recent phylogenetic analyses of morphology place chasmataspid body fossil species within Euchelicerata, as sister group to eurypterids (Garwood and Dunlop, 2014; Legg et al., 2013) or sister group to a clade composed of eurypterids and arachnids (Lamsdell, 2013; Lamsdell et al., 2015; Selden et al., 2015).

### 7.3. *Age justification*

The *Chasmataspis-like* resting traces were found in the Cambrian Hickory Sandstone Member of the Riley Formation, Texas (Dunlop et al., 2004). The top of the Hickory Sandstone preserves trilobite representatives of the *Bolaspidella* Zone and the *Cedarina* Zone (Miller et al., 2012; Palmer, 1954). These trilobite biozones are assigned to the lowermost Marjumiid Biomere in the Marjuman Stage of the Lincolnian Series (Miller et al., 2012). The lower Marjuman itself is correlated to the Drumian Stage of Cambrian Series 3 (Taylor et al., 2012). The end of the Drumian is dated to 500.5 Ma, providing a minimum age for *Chasmataspis-like* trace fossils.

Soft maximum as for 2.3.

## 8. Crown Xiphosurida

This clade comprises four extant species, all members of the family Limulidae: *Carcinoscorpius rotundicauda, Limulus polyphemus, Tachypleus gigas* and *Tachypleus tridentatus,* their last common ancestor and all of its descendants (Fig. 3). Monophyly is established by phylogenetic analyses of housekeeping genes (Obst et al., 2012) and morphology (Lamsdell and McKenzie, 2015).

### 8.1. *Fossil specimens*

*Tachypleus gadeai* Vía Boada and de Villalta, 1966. MGSB 19195, holotype.

### 8.2. *Phylogenetic justification*

*Heterolimulus gadeai* Vía Boada and de Villalta, 1966 was reassigned to the extant genus *Tachypleus* by Diedrich (2011), who cited the presence of lateral immobile opisthosomal spines as evidence. This was validated by the phylogenetic analysis of Lamsdell and McKenzie (2015), who resolved *T. gadeai* as sister-taxon to a clade composed of all other members of *Tachypleus.* This more inclusive clade in turn resolved as sister-taxon to the extant genus *Carcinoscorpius.*

### 8.3. *Age justification*

*T. gadeai* was discovered in the Alcover unit of the Montral site, Tarragona province, Catalonia, Spain (Vía Boada and de Villalta, 1966). Based on sequence stratigraphy, the Alcover dolomite unit is dated to the late Muschelkalk, a European regional stage of the Triassic (Calvet and Tucker, 1995; Vía Boada and de Villalta, 1966). The middle and late Muschelkalk correspond to the global Ladinian stage (Calvet and Tucker, 1995). The upper boundary of the Ladinian is 237.0 Ma ± 1 Myr (Ogg, 2012), thus, a minimum age of 236.0 Ma.

Soft maximum as for 2.3.

### 8.4. *Discussion*

We note that morphological phylogenetic analysis has suggested paraphyly of Xiphosura (crown Xiphosurida plus several fossil genera), and resolved synziphosurines as basal euchelicerates (Lamsdell, 2013). A subsequent morphological phylogeny resolved synziphosurines as polyphyletic (Garwood and Dunlop, 2014). Some other morphological phylogenies resolve the traditional monophyletic Xiphosura (Briggs et al., 2012; Legg et al., 2013).

Crown xiphosurid affinities of older fossils cannot be confirmed. For example, an undescribed Lower Ordovician fossil from Morocco (Van Roy et al., 2010) exhibits fused opisthosomal tergites, a synapomorphy of Xiphosurida *sensu* Lamsdell (2013), but its position with respect to the crown has not been tested in a phylogeny. The Upper Ordovician *Lunataspis aurora* Rudkin et al., 2008 and the Pennsylvanian genus *Euproops* are resolved on the stem group of Xiphosurida in a morphological phylogeny (Lamsdell, 2013).

Morphological conservatism in the evolution of Xiphosura (as illustrated by a Jurassic member assigned to a living genus) has led to use of the misnomer ‘living fossil’ for the clade, despite continued molecular evolution throughout its history (Avise et al., 1994; Obst et al., 2012). The more appropriate term of ’;stabilomorph’ was proposed with the discovery of *Limulus darwini* (Kin and Blazejowski, 2014); it refers to morphological stability over evolutionary history, at the genus level. However, long branches unbroken by unsampled extinct species may have significantly underestimated divergence times among crown Xiphosurida (Lamsdell and McKenzie, 2015), though this assertion has not yet been tested with a divergence time analysis.

## 9. Crown Arachnida

This clade comprises Acariformes (acariform mites), Opiliones (harvestmen), Palpigradi (microwhip scorpions), Parasitiformes (parasitiform mites, ticks), Pseudoscorpiones, Ricinulei (hooded tickspiders), Schizomida, Scorpiones, Solifugae (camel spiders), and Tetrapulmonata, their last common ancestor and all of its descendants (Fig. 4). Monophyly is established on the basis of phylogenetic analysis of transcriptomes (Rota-Stabelli et al., 2013a), nuclear protein-coding genes (Regier et al., 2010), strong support from morphology (Garwood and Dunlop, 2014; Legg et al., 2013; Rota-Stabelli et al., 2011; Shultz, 2007), and combined morphological and molecular data (Giribet et al., 2002; Lee et al., 2013). Some recent phylogenetic analyses of transcriptomes have failed to resolve a monophyletic Arachnida; instead Xiphosurida is variably placed within the traditional arachnids (Sharma et al., 2014; von Reumont et al., 2012).

### 9.1. *Fossil specimens*

*Palaeophonus loudonensis* Laurie, 1899. NMS 1897.122.196 (holotype), a dorsally preserved specimen lacking walking legs and telson (Fig. 5h). For additional anatomical detail, we refer to *Eramoscorpius brucensis* Waddington et al., 2015. ROM 53247, holotype (Fig. 5g).

### 9.2. *Phylogenetic justification*

The genus *Dolichophonus* Petrunkevitch, 1949, was erected for *P. loudonensis,* based on the increased length of the prosoma compared to other palaeophonids, namely *P. nuncius* Thorell and Lindstrom, 1885, and *P. caledonicus* Hunter, 1886. An examination (by D.A.L.) of the single known specimen of *P. loudonensis* could not confirm this character because the specimen is very poorly preserved, and the junction between the prosoma and mesosoma is not easily distinguished. We retain the holotype within *Palaeophonus.* It is even possible that this specimen may belong to one of the other currently recognised species of *Palaeophonus*, although more material would be required. *P. loudonensis* was included in a phylogenetic analysis by Legg et al. (2013), in which it resolved amongst total-group Scorpiones. Other Siluro-Devonian fossil scorpions such as *Proscorpius osborni* Whitfield, 1885 and *Palaeoscorpius devonicus* Lehmann, 1944 have been placed in the stem group of Orthosterni (crown-group Scorpiones) (e.g. Legg et al., 2013; Garwood and Dunlop, 2014), which are therefore crown group members of Arachnida.

### 9.3. *Age justification*

*P. loudonenesis* was recovered from the Gutterford Burn section of the “Eurypterid Bed” (Reservoir Formation) in the Pentland Hills, Midlothian, Scotland (Kjellesvig-Waering, 1986), which has been dated as Upper Llandovery. The associated graptolite fauna suggests a late Llandovery (Telychian) age for this formation, within the *Oktavites spiralis* Biozone (Bull and Loydell, 1995). A spline-fit age for the upper boundary of the *O. spiralis* Biozone provides a minimum age constraint of 435.15 Ma (Melchin et al., 2012).

Soft maximum as for 2.3.

## 10. Crown Opiliones

This clade comprises Cyphophthalmi and Phalangida (itself comprising Laniatores, Dyspnoi and Eupnoi), their last common ancestor and all of its descendants (Fig. 4). Monophyly has been demonstrated by phylogenetic analysis of transcriptomes (Sharma et al., 2014), morphology (Garwood et al., 2011), and combined morphology and molecules (Garwood et al., 2014; Giribet et al., 2002).

### 10.1. *Fossil specimens*

*Eophalangium sheari* Dunlop et al., 2003. PBM slide no. 3503, holotype (Fig. 5d), consisting of a lateral thin section outlining nearly the entire female body (Dunlop et al., 2003).

### 10.2. *Phylogenetic justification*

In a phylogenetic analysis of morphology, *E. sheari* was placed in a polytomy with members of Phalangida, to the exclusion of Cyphophthalmi (Garwood and Dunlop, 2014). In combined analysis of molecules and morphology, *E. sheari* was resolved as a stem group Cyphophthalmi (Garwood et al., 2014). Both positions, however, fall within the crown group of Opiliones.

### 10.3. *Age justification*

This fossil is known from the Lower Devonian (Pragian) Rhynie Chert of Aberdeenshire, Scotland. Spore assemblages of the Windyfield and stratigraphically underlying Rhynie Chert are dated to the early but not earliest Pragian to early (earliest?) Emsian (polygonalis-emsiensis Spore Assemblage Biozone) (Parry et al., 2011). Radiometric dating of the underlying Milton of Noth Andesite at ca. 411 Ma (Parry et al., 2013, 2011) has been subject to a dispute over its temporal relationship to hot spring activity associated with the cherts (Mark et al., 2013, 2011) and predates the biostratigraphic dating of the Rhynie Chert relative to the global dating of the base of the Pragian Stage. Therefore, a minimum age constraint may be defined at 405.0 Ma for the Rhynie Chert, using the Pragian-Emsian boundary (407.6 Ma ± 2.6 Myr) as a reference.

A soft maximum constraint comes from the oldest chelicerate *W. barbarhardyae* from the Emu Bay Shale on Kangaroo Island, South Australia, which has been correlated based on trilobite biostratigraphy to the upper part of the *Pararaia janeae* Zone in mainland South Australia (Jell in Bengtson et al., 1990; Fig. 2 in Jago et al., 2012). As this is equivalent to the Canglangpuan Stage in South China and the late Botoman in Siberia (Gehling et al., 2011, Fig. 9), the Emu Bay Shale can be dated to Cambrian Series 2, Stage 4, providing a maximum age of ~514 Ma.

## 11. Crown Acariformes

This clade comprises Sarcoptiformes, Trombidiformes and ‘Endeostigmata’, their last common ancestor and all of its descendants (Fig. 4). Monophyly is established by phylogenetic analysis of nuclear ribosomal genes (Pepato and Klimov, 2015), morphology (Garwood and Dunlop, 2014), and combined molecular and morphological data (Pepato et al., 2010).

### 11.1. *Fossil specimens*

*Protacarus crani* Hirst, 1923. BMNH In. 24665, holotype, preserving a nearly complete individual in chert (Fig. 5k).

### 11.2. *Phylogenetic justification*

Originally described as a single species (Hirst, 1923), *P. crani* from the Rhynie Chert was subsequently treated as five species belonging to five different genera (Dubinin, 1962). *P. crani* itself, as exemplified by the holotype, was described as a member of Eupodidae, within Trombidiformes (Hirst, 1923), or potentially more basal within the Acariformes (Bernini, 1986). While the specifics of its classification may be debatable due to the lack of preserved diagnostic characters, the fan-like setae observed dorsally in *P. crani* support a relationship with endeostigmatids within crown group Acariformes (Bernini, 1986; Dunlop and Selden, 2009).

### 11.3. *Age justification*

As for 10.3.

### 11.4. *Discussion*

Bernini et al. (2002) figured a putative oribatid mite from terrestrial sediments dated to the Early Ordovician of Sweden. However, its age and systematic placement were queried by Dunlop (2010, p. 134) and this species is not used for calibration herein.

Arguments that *P. crani* is a Recent contaminant (Crowson, 1985) have been countered by Kuhne and Schluter (1985) and Greenslade (1988). Other species of fossil Acariformes have also been described from the Rhynie Chert (all originally *P. crani*), including *Protospeleorchestes pseudoprotacarus, Pseudoprotacarus scoticus, Palaeotydeus devonicus,* and *Paraprotocarus hirsti* (all Dubinin, 1962).

## 12. Crown Parasitiformes

This clade comprises Opilioacariformes, Ixodida (hard and soft ticks), Holothyrida and Mesostigmata, their last common ancestor and all of its descendants (Fig. 4). Monophyly is established by phylogenetic analysis of nuclear ribosomal genes (Pepato and Klimov, 2015) and morphology (Garwood and Dunlop, 2014).

### 12.1. *Fossil specimens*

*Cornupalpatum burmanicum* Poinar and Brown, 2003. Holotype larva (accession number A-10-260; Fig. 5i) and paratype (accession number A-10–261; Fig. 5j) deposited in the Poinar amber collection maintained at Oregon State University, later to be deposited at the California Academy of Sciences (Poinar, 2015; Poinar and Brown, 2003).

### 12.2. *Phylogenetic justification*

Morphological characters such as the subcircular body with a marginal groove, free coxae, ventral anal opening, the presence of a capitulum and Haller’s organ, absence of an anal groove, and elongate four-segmented palpi are all suggestive of Parasitiformes affinity for *C. burmanicum* (Poinar and Brown, 2003). A particularly diagnostic character, suggesting placement within at least total-group Ixodida (and thus crown Parasitiformes), is the presence of claws on palpal segment 3 in the larva (Poinar and Brown, 2003). Putative morphologies similar to bacterial pathogens exclusive to modern Ixodida were recently described from the paratype (Poinar, 2015).

### 12.3. *Age justification*

This fossil is sourced from amber mines in the Hukawng Valley of Kachin State, northern Myanmar (formerly Burma). The depositional age of Burmese amber was estimated from U-Pb dating of zircons from the volcaniclastic matrix surrounding the amber (Shi et al., 2012). Shi et al. (2012) argue the amber is not older than its associated sediments, as burial and preservation would have to be rapid for survival of organic material, so the amber was probably formed at, but not earlier than the U-Pb date: at 98.79 Ma ± 0.62 Myr. Therefore, a minimum age for any fossil from Burmese amber deposits is 98.17 Ma.

Soft maximum as for 10.3.

## 13. Crown Pseudoscorpiones

This clade comprises Feaelloidea, Chthonioidea, Neobisiodea, Garypoidea, Sternophoroidea and Cheliferoidea, their last common ancestor and all of its descendants (Fig. 4). While relationships between superfamilies remain unclear, monophyly of Pseudoscorpiones was demonstrated with wide taxon sampling and three genes (Murienne et al., 2008). More limited taxon sampling supports monophyly with morphology (Garwood and Dunlop, 2014) and morphology combined with ribosomal genes (Pepato et al., 2010).

### 13.1. *Fossil specimens*

*Protofeaella peetersae* Henderickx in Henderickx and Boone, 2016. NHM II 3115, holotype, near complete specimen preserved in amber (Fig. 5c).

### 13.2. *Phylogenetic justification*

Although *P. peetersae* has not been included in a formal phylogenetic analysis, it was assigned to the extant family Feaellidae by Henderickx and Boone (2016:8), based on its narrow cephalothorax, granulated abdomen, and presence of small pedipalps with narrow coxa and small hands. Whilst these features are certainly found in both *P. peetersae* and feaellids, other features, such as slender pedipalp fingers, and the overall shape of the cephalic shield are more like those of pseudogarypids (Harvey, 1992). Both the feaellid and pseudogarypids belong to the superfamily Feaelloidea *(sensu* Harvey, 1992), and thus it is still likely *P. peetersae* belongs within the pseudoscorpion crown-group.

### 13.3. *Age justification*

As for 12.3.

### 13.4. *Discussion*

*Dracochela deprehendor* Schawaller et al., 1991 from the Middle Devonian of Gilboa, New York State, was originally interpreted as a crown-group pseudoscorpion but was reassigned to the pseudoscorpion stem group (Judson, 2012). Preliminary morphological phylogenetic analyses by one of us (D.A.L.), however, suggest that *D. deprehendor* may be within the crown group. If so, this would drastically extend the range of crown Pseudoscorpiones by over 280 Myr (to a minimum age of 382.7 Ma; Richardson et al., 1993).

Note that other Cretaceous pseudoscorpions have been preserved in amber (older from Lebanon and Spain; younger from France, New Jersey and Alberta), but these have yet to be fully described or examined for their systematic positions (Judson, 2009). If the Lebanese or Spanish fossils were found to be members of the crown group, they could extend the range of Pseudoscorpiones by up to 27 Myr.

## 14. Crown Palpigradi

This clade comprises the families Eukoeneniidae and Prokoeneniidae, their last common ancestor and all of its descendants (Fig. 4). Monophyly has been supported by a phylogenetic analysis of two nuclear ribosomal and one mitochondrial gene (Giribet et al., 2014).

### 14.1. *Fossil specimens*

*Electrokoenenia yaksha* Engel et al., 2016b. NIGP 163253, holotype in amber (Fig. 5e).

### 14.2. *Phylogenetic justification*

*E. yaksha* was classified within the extant family Eukoeneniidae (Engel et al., 2016b). This was justified with morphological characters, in particular the rounded lateral “arms” to the frontal organ of the propeltidium, as seen in the extant genus *Leptokoenenia* (Engel et al., 2016b). Thus *E. yaksha* is within the crown group of Palpigradi.

### 14.3. *Age justification*

As for 12.3.

## 15. Crown Solifugae

This clade comprises Ammotrechidae, Ceromidae, Daesiidae, Eremobatidae, Galeodidae, Gylippidae, Hexisopodidae, Karschiidae, Melanoblossidae, Mummuciidae, Rhagodidae and Solpugidae, their last common ancestor and all of its descendants (Fig. 4). No phylogenetic analysis has yet included all families, but limited taxon sampling has shown monophyly with morphology (Garwood and Dunlop, 2014; Shultz, 2007) and morphology combined with nuclear genes (Giribet et al., 2002; Pepato et al., 2010). Extensive morphological work on the homology of cheliceral characters was recently published by Bird et al. (2015).

### 15.1. *Fossil specimens*

*Cratosolpuga wunderlichi* Selden and Shear, 1996 (Sol. 1 in the private Wunderlich collection, Straubenhardt, Germany), holotype. An additional specimen (SMNK 1268 PAL; Fig. 5f), not a paratype, is also deposited.

### 15.2. *Phylogenetic justification*

*C. wunderlichi* was assigned to the extant family Ceromidae on the basis of shape of the cheliceral flagellum, shape of the propeltidium, eye tubercle, and leg spination (Selden and Shear, 1996). Only a single tarsal segment is present on the legs (Selden and Shear, 1996). A position in total-group Ceromidae would therefore be within crown-group Solifugae.

### 15.3. *Age justification*

This fossil was recovered from the Nova Olinda Member of the Crato Formation in the Araripe Basin, northeastern Brazil. This unit is generally agreed to be around the Aptian/Albian border (Martill et al., 2007). Batten (2007) suggests that if assemblages in the upper layers are consistent across the lower layers, a late Aptian date should be considered. The Crato formation has been dated using palynomorphs (Pons et al., 1990) to the Aptian, though an accurate date for the Nova Olinda Member is not available. The upper boundary of the Aptian, at 113.0 Ma ± 0.4 Myr, gives a minimum date of 112.6 Ma.

Soft maximum as for 10.3.

### 15.4. *Discussion*

The Pennsylvanian *Protosolpuga carbonaria* Petrunkevitch, 1913, the only older possible fossil solifuge, was discounted from the crown group of Solifugae in the same paper as described *C. wunderlichi* (Selden and Shear, 1996). It is too poorly preserved to assign to the crown group without additional phylogenetic justification.

## 16. Crown Scorpiones

This clade comprises Buthida and Iurida, their last common ancestor and all of its descendants (Fig. 4). The composition of Buthida and Iurida are as detailed in Sharma et al. (2015). Monophyly has been supported by phylogenetic analysis of transcriptomes (Sharma et al., 2015, 2014), morphology (Garwood and Dunlop, 2014; Shultz, 2007), and combined ribosomal sequences and morphology (Pepato et al., 2010).

### 16.1. *Fossil specimens*

*Protoischnurus axelrodurum* Carvalho and Lourenço, 2001. MN-7601-I, holotype, a male. We also refer to the specimen SMNS 65534, which preserves phylogenetically relevant details of the pedipalps (Fig. 3c in Menon, 2007).

### 16.2. *Phylogenetic justification*

Menon (2007) placed *P. axelrodurum* in the extant family Hemiscorpiidae, based on, amongst other things, an inverse Y-shape on sulcus on the cephalic shield, the placement of Est trichobothria on the pedipalp chela, and the placement of carinae V2 and V3 in the pedipalp chela, all of which are diagnostic of the hemiscorpidid subfamily Hormurinae (Soleglad et al., 2005). Hemiscorpiidae is classified within Iurida (Sharma et al., 2015), and is thus crown group Scorpiones.

### 16.3. *Age justification*

As for 15.3.

### 16.4. *Discussion*

A number of fossil taxa have been placed within crown group scorpion families, including *Protobuthus elegans* Lourenço and Gall, 2004, from the Early Triassic Buntsanstein of France. It was assigned to the superfamily Buthoidea, however, a subsequent study (Baptista et al., 2006), considered this taxon and *Archaeobuthus* from the Early Cretaceous of Lebanon, outside the crown-group based on trichobothrial arrangement.

*Araripescorpius ligabuei* Campos, 1986 is coeval to *P. axelrodurum,* and from the same locality. Menon (2007) placed *A. ligabuei* in the extant family Chactidae based on general habitus and trichobothrial pattern. Therefore it is also a member of the crown group of Scorpiones, albeit a less well-preserved species.

## 17. Crown Tetrapulmonata

This clade comprises Pedipalpi and Araneae (spiders), their last common ancestor and all of its descendants (Fig. 6). Monophyly is supported by phylogenetic analysis of transcriptomes (Sharma et al., 2014), nuclear protein-coding genes (Regier et al., 2010), and morphology (Garwood and Dunlop, 2014; Legg et al., 2013; Shultz, 2007). This clade is not recognized in GenBank taxonomy.

**Fig. 6.**
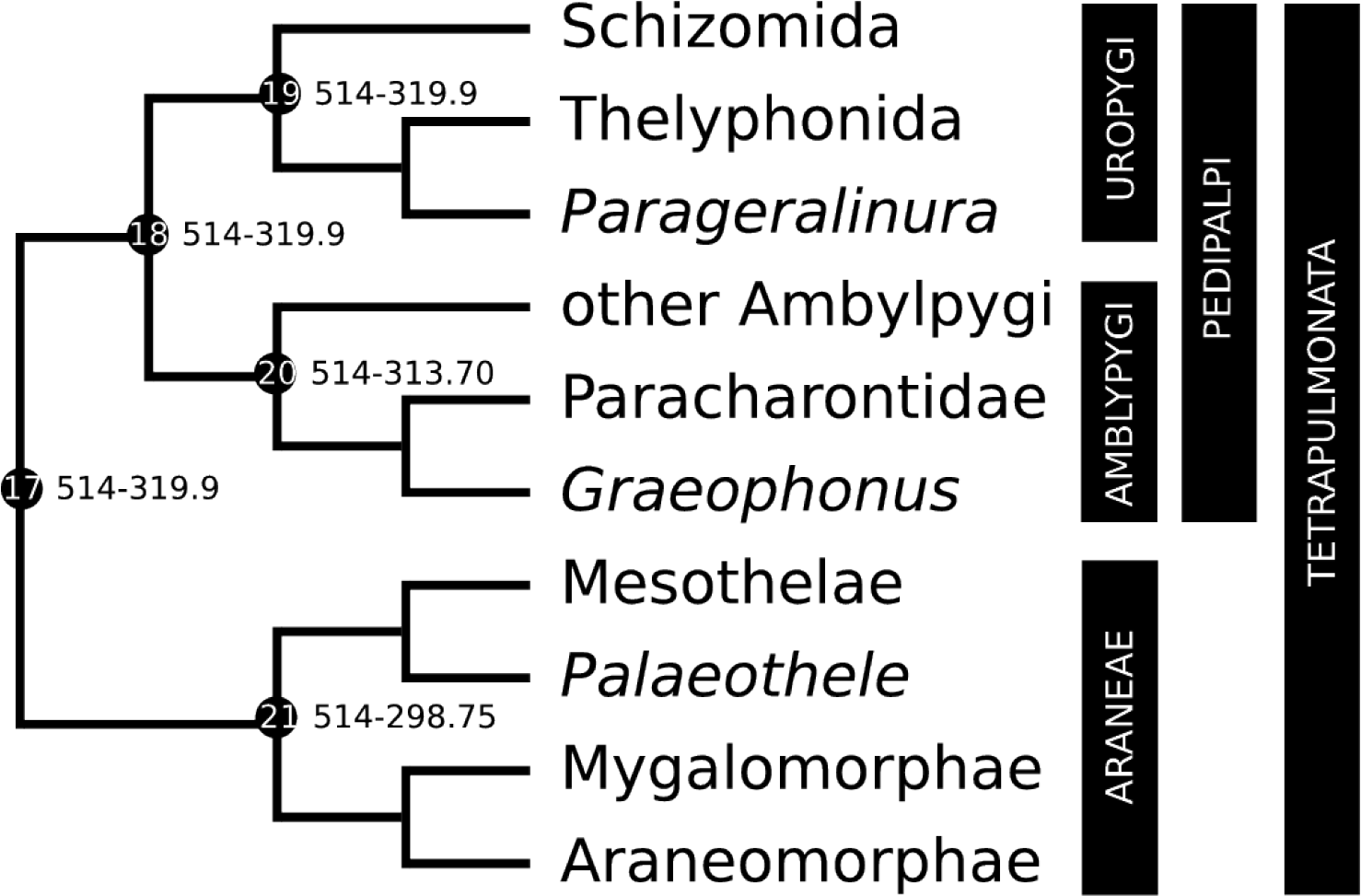
Calibration diagram for Tetrapulmonata (nodes 17–21).

### 17.1. *Fossil specimens*

*Parageralinura naufraga* (Tetlie and Dunlop, 2008), LWL Ar.K.1 (Fig. 7a). Counterpart in the private collection of W. Elze, Ennepetal, Germany.

**Fig. 7.**
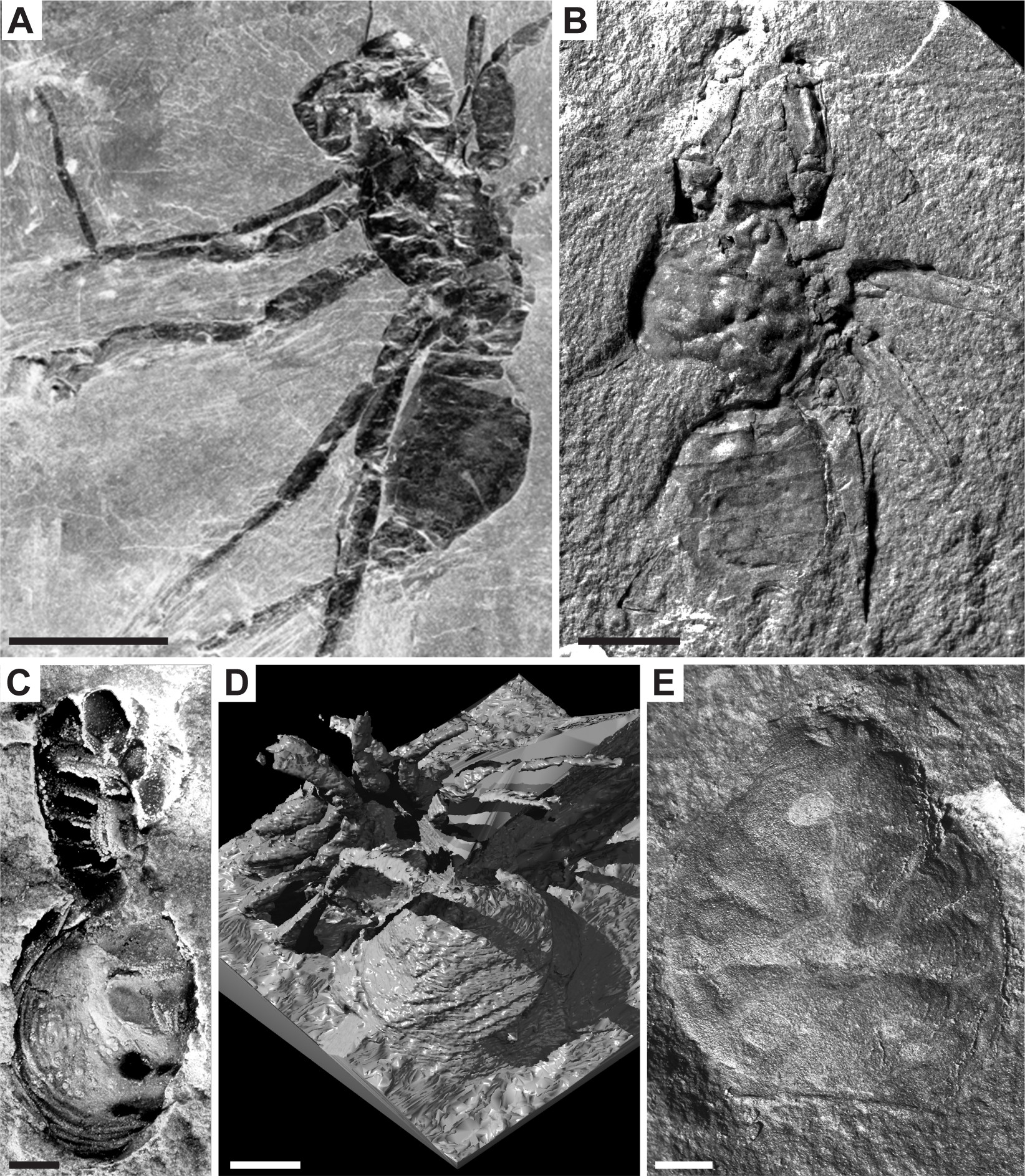
Tetrapulmonata fossil calibrations for (A) nodes 17–19: *Parageralinura naufraga,* LWL Ar.K.1, scale bar 5mm, image credit C. Brauckmann; (B) node 20: *Graeophonus anglicus,* BMNH In 31233, scale bar 5mm, image credit J. Dunlop; (C, D) node 21: *Palaeothele montceauensis,* images credit P. Selden; (C) Holotype MNHN 51961, scale bar 2mm; (D) Reconstructed specimen BMNH In.62050, scale bar 0.2mm; (E) node 21: *Arthrolycosa* sp., PIN 5431/9, scale bar 1mm, image credit P. Selden.

### 17.2. *Phylogenetic justification*

Tetlie and Dunlop (2008) interpreted Coal Measures uropygids to comprise a plesion on the stem of the extant Thelyphonidae, the sole extant family of Thelyphonida. A subchelate pedipalp is considered apomorphic of the crown group but is lacking in *Geralinura* and *P. naufraga.* This identifies them as crown Uropygi, and thus, Tetrapulmonata.

### 17.3. *Age justification*

Of the uropygid fossils, the oldest are *P. naufraga* (formerly *Prothelyphonus naufragus)* from deposits of “Ziegelei-Grube,” Hagen-Vorhalle, Nordrhein-Westphalia, Germany (Brauckmann and Koch, 1983; Tetlie and Dunlop, 2008). The fossil-bearing deposits are assigned to the Namurian B (Marsdenian) based on the *Bilinguites metabilinguis* R2c1 subzone of ammonoid stratigraphy (Brauckmann et al., 1994; Tetlie and Dunlop, 2008). The (upper) Namurian-(lower) Westphalian boundary is defined by the earliest occurrence of the goniatite *Gastrioceras subcrenatum* (Waters and Davies, 2006), but lacks a precise isotopic date. Pointon et al. (2012) estimated an age of c. 319.9 Ma for the base of the Westphalian (top of the Namurian, only slightly younger than the Marsdenian) based on Milankovitch cycles of sedimentation, giving a minimum age for *P. naufraga.*

Soft maximum as for 10.3.

## 18. Crown Pedipalpi

This clade comprises Amblypygi (tailless whip scorpions) and Uropygi, their last common ancestor and all of its descendants (Fig. 6). Monophyly is supported by phylogenetic analysis of transcriptomes (Sharma et al., 2014), nuclear protein-coding genes (Regier et al., 2010), and morphology (Garwood and Dunlop, 2014; Legg et al., 2013; Shultz, 2007). This clade is not recognized in GenBank taxonomy.

### 18.1. *Fossil specimens*

As for 17.1.

### 18.2. *Phylogenetic justification*

As for 17.2.

### 18.3. *Age justification*

As for 17.3.

## 19. Crown Uropygi

This clade comprises Thelyphonida (whip scorpions) and Schizomida, their last common ancestor and all of its descendants (Fig. 6). Monophyly is supported by phylogenetic analysis of nuclear protein-coding genes (Regier et al., 2010), and morphology (Garwood and Dunlop, 2014; Legg et al., 2013; Shultz, 2007).

### 19.1. *Fossil specimens*

As for 17.1.

### 19.2. *Phylogenetic justification*

As for 17.2.

### 19.3. *Age justification*

As for 17.3.

## 20. Crown Amblypygi

This clade comprises Paracharontidae, Charinidae, Charontidae, Phrynichidae and Phrynidae, their last common ancestor and all of its descendants (Fig. 6). Monophyly of Amblypygi has not been fully investigated with phylogenetic analysis; however, monophyly has been shown for at least some families with morphological data (Garwood and Dunlop, 2014; Shultz, 2007) and morphology combined with nuclear genes (Pepato et al., 2010).

### 20.1. *Fossil specimens*

*Graeophonus anglicus* Pocock, 2011. BMNH In. 31233, holotype (Fig. 7b). Figured in Dunlop et al. (2007: Fig. 1 a,b).

### 20.2. *Phylogenetic justification*

*G. anglicus* was redescribed by Dunlop et al. (2007) as a member of the Amblypygi crown group. This was based on several morphological character comparisons to living members, such as the pedipalp femur with dorsal spination similar to *Paracharon* (the monotypic extant species of the family Paracharontidae). *G. anglicus*, unlike *Paracharon*, has a pear-shaped ocular tubercle, suggesting it was not blind. *G. anglicus* is inferred to be on the stem lineage of Paracharontidae, and thus, crown group Amblypygi (Dunlop et al., 2007).

### 20.3. *Age justification*

The genus *Graeophonus* was originally described from the Sydney Basin, Cape Breton Carboniferous Coal Measures, Nova Scotia, Canada, which corresponds to Westphalian in age (Dunlop et al., 2007; Giles et al., 2002; Scudder, 1890a). Further studies are needed on the Canadian material, so the minimum age was taken from the oldest European specimen (which is roughly the same age as the Cape Breton specimen) from the British Middle Coal Measures (Coseley, Stafffordshire), which is Westphalian B (or Duckmantian) at the youngest (Waters et al., 1994; Waters and Davies, 2006). U-Pb dating of zircons constrains the upper boundary of the Duckmantian to 313.78 Ma ± 0.08 Myr (Pointon et al., 2012), so a minimum age for *G. anglicus* is 313.70 Ma.

Soft maximum as for 10.3.

## 21. Crown Araneae

This clade comprises Mesothelae, Mygalomorphae (tarantulas, funnel-web spiders) and Araneomorphae (most spiders), their last common ancestor and all of its descendants (Fig. 6). Monophyly is established by phylogenetic analysis of transcriptomes (Fernandez et al., 2014a; Garrison et al., 2016) and morphology (Garwood and Dunlop, 2014; Legg et al., 2013; Shultz, 2007).

### 21.1. *Fossil specimens*

*Palaeothele montceauensis* Selden, 2000. Museum d'Histoire naturelle, Autun: 51961 (holotype; Fig. 7c), and BMNH 62050, second specimen (not a paratype, Fig. 7d; Selden, 1996). We also compare to *Arthrolycosa* sp. B Selden et al. 2014. PIN 5431/9 (Fig. 7e).

### 21.2. *Phylogenetic justification*

The deep, narrow sternum of *P. montceauensis* (formerly *Eothele montceauensis* Selden, 1996) is shared by extant members of Mesothelae (Selden, 1996). Several other characters that are symplesiomorphic within Araneae, such as spinnerets, suggest a position within crown Araneae, and perhaps on the stem lineage of Mesothelae (Selden, 1996).

### 21.3. *Age justification*

*P. montceauensis* was found in the Montceau Lagerstätte, Montceau-les-Mines, Massif Central, France. The nodule-rich layer is just above the first seam in the Great Seams Formation (late Stephanian) (Perrier and Charbonnier, 2014). The age within the Stephanian has been assigned to Stephanian B, with some biostratigraphic evidence for Stephanian C (Racheboeuf et al., 2002). The Stephanian B/C is a European stage of the Pennsylvanian, straddling the boundary of the globally used Kasimovian and Gzhelian (Richards, 2013). The younger bound of the Gzhelian is 298.9 Ma ± 0.15 Myr, hence the minimum age of the Montceau Lagerstätte is 298.75 Ma.

Soft maximum as for 10.3.

### 21.4 *Discussion*

A possible older spider, *Arthrolycosa* sp. B Selden et al., 2014, is known from the Krasnodonsky Horizon of Rostov Province, Russia (Duckmantian, ~313 Ma). It shares characters with extant Mesothelae, such as the position and morphology of the eye tubercles, but lacks spinnerets, so the inference is largely based on better-known *Arthrolycosa* from other localities (Selden et al., 2014). Because phylogenetic evidence for crown group membership is largely indirect, we maintain *P. montceauensis* as the oldest well-supported Araneae.

## 22. Crown Mandibulata

This clade comprises Myriapoda and Pancrustacea, their last common ancestor and all of its descendants (Fig. 1). Monophyly has been independently demonstrated based on protein-coding genes and microRNAs, as well as morphological data (Rota-Stabelli et al., 2011).

### 22.1. *Fossil specimens*

As for 4.1.

### 22.2. *Phylogenetic justification*

As for 4.2.

### 22.3. *Age justification*

As for 4.3.

### 22.4. *Discussion*

*Wujicaris muelleri* Zhang et al., 2010 has an equal claim to being the earliest record of Mandibulata, but it is of equal age to the holotype of *Y. dianensis.*

## 23. Crown Myriapoda

This is the clade comprising Chilopoda (centipedes) and Progoneata, their last common ancestor and all of its descendants (Fig. 1). Monophyly has been demonstrated by morphology (Edgecombe, 2004; Legg et al., 2013; Rota-Stabelli et al., 2011), nuclear protein-coding genes (Regier et al., 2010; Zwick et al., 2012), transcriptomes (Rehm et al., 2014), and combined analysis of molecules and morphology (Lee et al., 2013).

### 23.1. *Fossil specimens*

*Cowiedesmus eroticopodus* Wilson and Anderson, 2004. AMS F.64845, holotype (Fig. 2e).

### 23.2. *Phylogenetic justification*

Membership of *C. eroticopodus* in Diplopoda is indicated by its strict diplosegmentation, whereas its cuticular mineralization supports membership in the subgroup Chilognatha, and its modified legs on trunk segment 8 support membership in Helminthomorpha. *C. eroticopodus* is resolved as total-group Helminthomorpha in the morphological cladistic analysis of Fernandez et al. (2016) and is accordingly a member of the crown-groups of Chilognatha, Diplopoda, Progoneata and Myriapoda.

### 23.3. *Age justification*

The earliest myriapods in the body fossil record are three species of Diplopoda from the *Dictyocaris* Member of the Cowie Formation at Cowie Harbour, near Stonehaven, Aberdeenshire, Scotland, one of which is *C. eroticopodus* (Wilson and Anderson, 2004). Based on associated spores, the Cowie Formation taxa are late Wenlock to early Ludlow in age (Marshall, 1991; Wellman, 1993) and the Early Ludlow upper boundary (Gorstian-Ludfordian boundary) is 425.6 Ma ± 0.9 Myr, so the minimum age for Myriapoda is 424.7 Ma.

Soft maximum as for 2.3.

### 23.4. *Discussion*

*Albadesmus almondi* and *Pneumodesmus newmani* (both described by Wilson and Anderson, 2004) have an equal claim to being the oldest myriapod, sourced from the same locality and unit as *C. eroticopodus.* The latter was selected because it has been explicitly coded in a morphological cladistic analysis (Fernandez et al., 2016). We have not used trace fossil evidence suggestive of Ordovician diplopods (Wilson, 2006) for dating.

## 24. Crown Progoneata

This clade comprises Diplopoda (millipedes), Pauropoda and Symphyla, their last common ancestor and all of its descendants (Fig. 8). Monophyly is supported by phylogenetic analysis of nuclear protein-coding genes (Regier et al., 2010; Zwick et al., 2012), whole mitochondrial genomes (Brewer et al., 2013), and morphology (Edgecombe, 2004; Legg et al., 2013). Two recent molecular phylogenies reject monophyly of Progoneata in favour of a putative clade of Chilopoda and Diplopoda: one based on three protein-coding genes (Miyazawa et al., 2014) and one on transcriptomes (Rehm et al., 2014). This clade is not recognized in GenBank taxonomy.

**Fig. 8.**
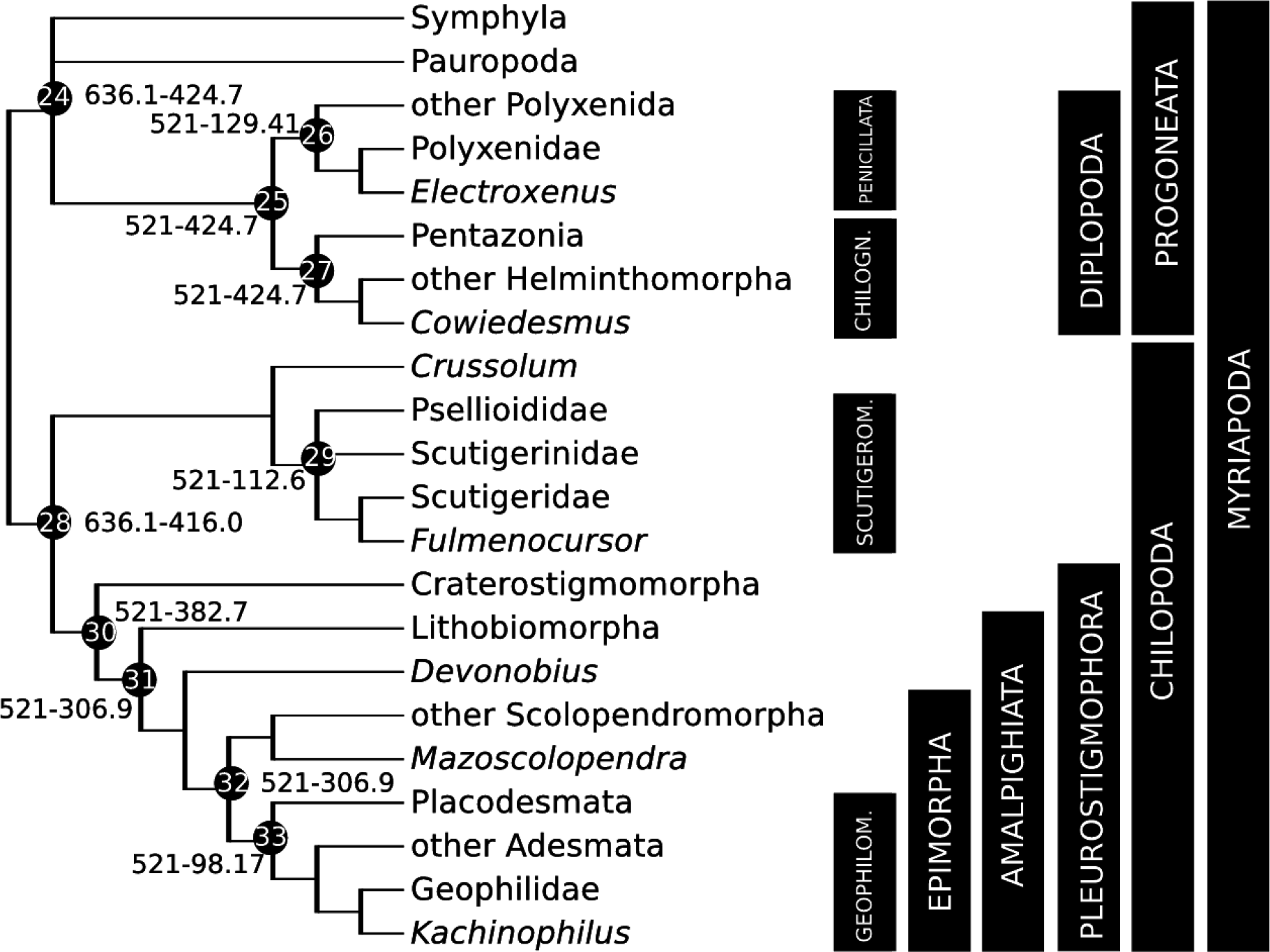
Calibration diagram for Myriapoda (nodes 24–33). Chilogn. = Chilognatha, Scutigerom. = Scutigeromorpha, Geophilom. = Geophilomorpha.

### 24.1. *Fossil specimens*

As for 23.1.

### 24.2. *Phylogenetic justification*

As for 23.2.

### 24.3. *Age justification*

As for 23.3.

### 24.4. *Discussion*

As for 23.4.

## 25. Crown Diplopoda

This clade comprises Penicillata (bristly millipedes) and Chilognatha, their last common ancestor and all of its descendants (Fig. 8). Monophyly is supported by morphological analyses (Blanke and Wesener, 2014), nuclear protein-encoding genes (Regier et al., 2010), and transcriptomes (Fernandez et al., 2016; Rehm et al., 2014).

### 25.1. *Fossil specimens*

As for 23.1.

### 25.2. *Phylogenetic justification*

As for 23.2.

### 25.3. *Age justification*

As for 23.3.

### 25.4. *Discussion*

As for 23.4.

## 26. Crown Penicillata

This clade comprises Polyxenoidea and Synxenoidea, their last common ancestor and all of its descendants (Fig. 8). Monophyly has been defended based on the shared presence of serrate setae arranged in lateral and caudal tufts (Enghoff, 1984).

### 26.1. *Fossil specimens*

*Electroxenus jezzinensis* Nguyen Duy-Jacquemin and Azar, 2004 (Acra collection, provisionally deposited at MNHN: JS 231/1), holotype (Fig. 9a,b), adult in amber (Nguyen Duy-Jacquemin and Azar, 2004, Fig. 1A,B).

**Fig. 9.**
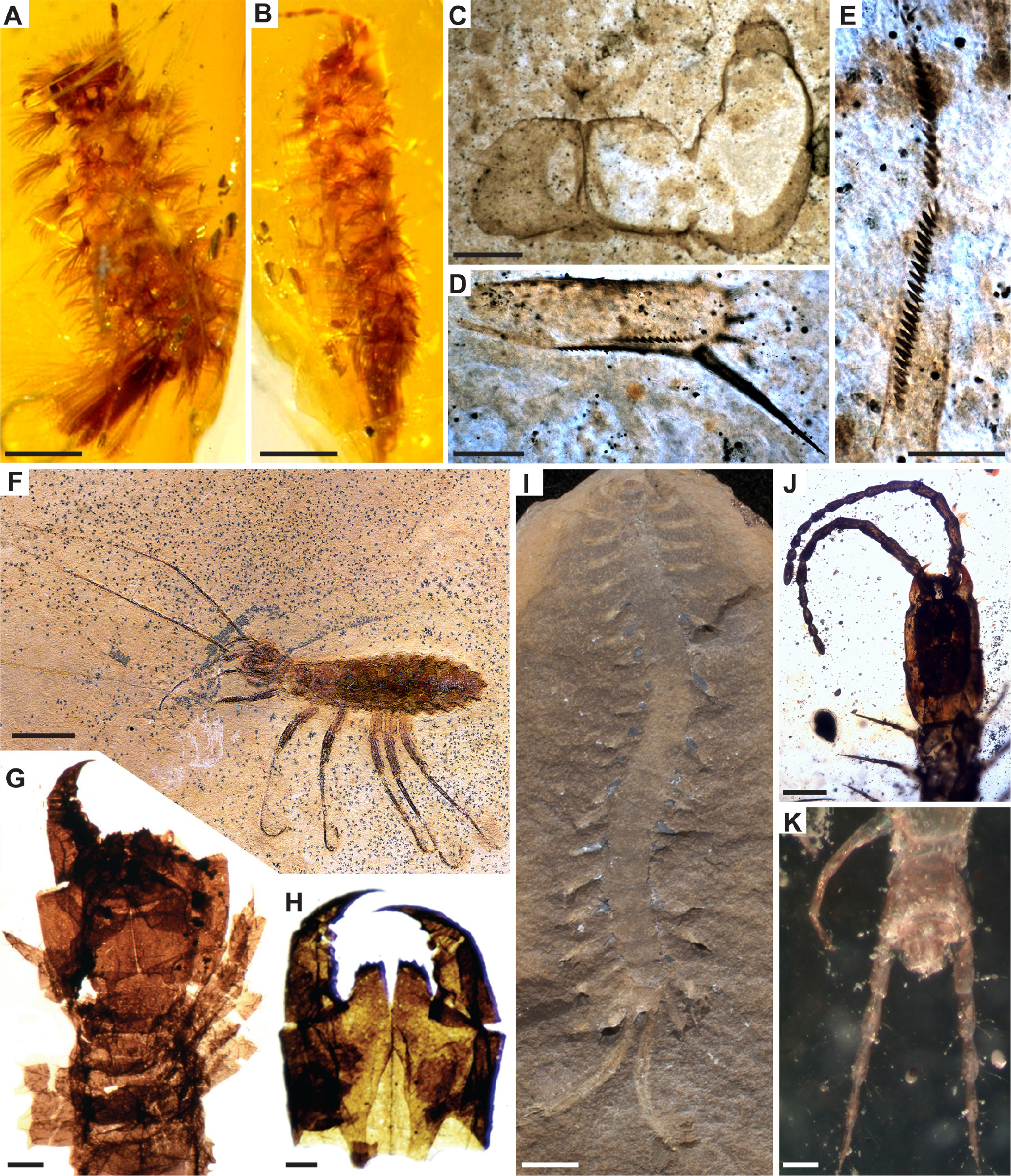
Myriapod fossil calibrations for (A-B) node 26: *Electroxenus jezzinensis,* MNHN JS 231/1, scale bar 0.5mm, image credit D. Azar; (C-E) node 28: *Crussolum* sp., images credit L. Anderson; (C) Forcipular segment, slide AUGD 12308, scale bar 0.5mm; (D) Prefemur of a trunk leg, slide AUGD 12307, scale bar 0.2mm; (E) Tibia of a trunk leg, slide AUGD 12307, scale bar 0.2mm; (F) node 29: *Fulmenocursor tenax,* SMNS 64275, scale bar 5mm, image credit G. Bechly; (G-H) node 30: *Devonobius delta,* scale bars 0.1mm, images credit W. Shear; (G) Head and anterior part of trunk, AMNH slide 411–15-AR18; (H) Forcipular segment, AMNH slide 4329-AR4; (I) nodes 31–32: *Mazoscolopendra richardsoni,* FMNH PE29002, scale bar 5mm, image credit J. Wittry; (J-K) node 33: *Kachinophilus pereirai,* AMNH Bu-Ba41a; (J) Head and anterior part of trunk, scale bar 0.3 mm; (K) Posterior body segments, scale bar 0.1mm.

### 26.2. *Phylogenetic justification*

Cretaceous amber penicillates are readily assigned to two of the three extant families, Polyxenidae and Synxenidae (Nguyen Duy-Jacquemin and Azar, 2004; classification of Penicillata following Short in Enghoff et al., 2015). *E. jezzinensis* preserves diagnostic characters of Polyxenidae such as lateral extensions of the gnathochilarial palps. Membership in an extant family indicates status as crown Penicillata.

### 26.3. *Age justification*

*E. jezzinensis* was discovered in amber from the Jouar Ess-Souss locality, in the Jezzine area, South Lebanon (Azar et al., 2010). Previous work suggested a Neocomian (Valanginian-Hauterivian) age for the Jezzine area (Azar et al., 2010). However, Lebanese stratigraphy has recently been revised; the Jouar Ess-Souss locality is now recognized as part of the lowermost interval of the Grès du Liban (Maksoud et al., 2016). The lower interval lies below a shale layer bearing the echinoid fossil *Heteraster oblongus,* and below a pisolitic interval bearing charyophyte fossils (Maksoud et al., 2016). The charyophyte layer is associated to the *Cruciata-Paucibracteatus* Zone of Martm-Closas et al. (2009) in the late Barremian-early Aptian, but this layer is also older than the Banc de Mrejatt subunit within Lebanon, thus Jezzine amber is older than the Ba2 layer in Fig. 6 of Maksoud et al. (2016). Jezzine amber is therefore no younger than early Barremian. The upper boundary of the early Barremian is proposed to be the first appearance of the ammonite *Ancyloceras vandenheckii* (Ogg et al., 2012). Cyclostratigraphy dates the *A vandenheckii* Zone beginning at 129.41 Ma (Ogg et al., 2012), providing a minimum age for Jezzine Lebanese amber fossils.

A soft maximum age is obtained from the oldest mandibulate, *Y. dianensis,* which was recovered from the Yu’anshan Formation at Xiaotan section, Yongshan, Yunnan Province, attributed to the *Eoredlichia-Wutingaspis* Biozone (Zhang et al., 2007). Chinese Cambrian stratigraphy has been revised substantially and the *Eoredlichia-Wutingaspis* Biozone is no longer recognized (Peng, 2009, 2003). However, *Eoredlichia* is known to co-occur with *Hupeidiscus,* which is diagnostic of the *Hupeidiscus-Sinodiscus* Biozone, which is formally recognised as the second biozone of the Nangaoan Stage of the Qiandongian Series of the Cambrian of China (Peng and Babcock, 2008). The Nangaoan is the proposed third stage of the Cambrian System for the International Geologic Timescale (Peng et al., 2012). Thus, a soft maximum constraint can be established on the age of the lower boundary of the Nangaoan, which has been dated to 521 Ma (Peng et al., 2012; Peng and Babcock, 2008).

### 26.4. *Discussion*

Another species of Polyxenidae from Lebanese amber, *Libanoxenus hammanaensis* Nguyen Duy-Jacquemin and Azar, 2004, is likewise known from a single specimen, from the Mdeiriji/Hammana locality in Central Lebanon. Its age data are similar to those of the more completely known *E. jezzinensis,* so the latter is used for calibration.

## 27. Crown Chilognatha

This clade comprises Pentazonia (pill millipedes) and Helminthomorpha (long-bodied millipedes), their last common ancestor and all of its descendants (Fig. 8). Monophyly is supported by morphological analyses (Blanke and Wesener, 2014), nuclear coding genes (Miyazawa et al., 2014), and transcriptomes (Fernandez et al., 2016). Chilognthan monophyly has rarely been opposed: some analyses of nuclear protein coding genes by Regier et al. (2005) found weak support for an unconventional grouping of Penicillata with Pentazonia, but others retrieved Chilognatha.

### 27.1. *Fossil specimens*

As for 23.1.

### 27.2. *Phylogenetic justification*

As for 23.2.

### 27.3. *Age justification*

Minimum as for 23.3. Soft maximum as for 26.3.

### 27.4. *Discussion*

As for 23.4.

## 28. Crown Chilopoda

This clade comprises Scutigeromorpha (house centipedes) and Pleurostigmophora, their last common ancestor and all of its descendants (Fig. 8). Monophyly is robustly supported by morphological analyses (Edgecombe and Giribet, 2004; Murienne et al., 2010), nuclear protein-coding genes (Miyazawa et al., 2014; Regier et al., 2010; Zwick et al., 2012), and transcriptomics (Fernandez et al., 2016; Rehm et al., 2014).

### 28.1. *Fossil specimens*

*Crussolum* sp. Jeram et al., 1990. DE 1.3.2/50 (Fig. 1N) and DE 3.1.1/88 (Fig. 1P; Jeram et al., 1990). As mentioned below, we also refer to material from the Windyfield Chert (AUGD 12307–12308; Anderson and Trewin, 2003) for morphological details (Fig. 9c-e).

### 28.2. *Phylogenetic justification*

*Crussolum* was resolved as stem-group Scutigeromorpha in the morphological cladistic analysis of Fernandez et al. (2016). Codings were a composite of material described as *Crussolum* sp. from the Windyfield Chert (Pragian) of the Dryden Flags Formation, Aberdeenshire, Scotland (figured by Anderson and Trewin, 2003), and the one formally named species of the genus, *C. crusseratum* (Shear et al., 1998), known from isolated and fragmentary legs from the Middle Devonian Gilboa locality, Schoharie County, New York State (Givetian).

### 28.3. *Age justification*

The oldest examples of *Crussolum* are isolated legs from Ludford Lane in England (Shear et al., 1998), which come from a horizon 0.15–0.20 m above the base of the Ludlow Bone Bed Member, in the Downtown Castle Sandstone Formation. The Ludlow Bone Bed Member is early Pridoli in age (Jeram et al., 1990), that stage having an upper boundary of 419.2 Ma ± 3.2 Myr, providing a minimum age of 416.0 Ma. *Crussolum* as currently delimited crosses the Silurian-Devonian boundary.

Soft maximum as for 2.3.

## 29. Crown Scutigeromorpha

This clade comprises Scutigeridae, Scutigerinidae and Psellioididae, their last common ancestor and all of its descendants (Fig. 8). Monophyly is robustly supported by morphological analyses (Edgecombe and Giribet, 2004), targeted gene sequencing (Murienne et al., 2010), and transcriptomics (Fernandez et al., 2016).

### 29.1. *Fossil specimens*

*Fulmenocursor tenax* Wilson, 2001. SMNS 64275, holotype (Fig. 9f), nearly complete specimen (Wilson, 2001, Pl. 1, Fig. 2).

### 29.2. *Phylogenetic justification*

Wilson (2001) assigned *F. tenax* to the extant family Scutigeridae based on the proportions of its antennal articles and its styliform male gonopods. Paired spine-bristles on the tibia of the second maxilla (synapomorphy of Pselliodidae + Scutigeridae) are consistent with this interpretation (Edgecombe, 2011). These affinities place the genus in crown-group Scutigeromorpha.

### 29.3. *Age justification*

Minimum as for 15.3. Soft maximum as for 26.3.

### 29.5 *Discussion*

A total-group scutigeromorph from the Carboniferous Mazon Creek deposits, *Latzelia primordialis* Scudder, 1890b, cannot be reliably assigned to the scutigeromorph crown group (Edgecombe, 2011; J. T. Haug et al., 2014) and is accordingly not used for dating that clade.

## 30. Crown Pleurostigmophora

This clade comprises Craterostigmomorpha and Amalpighiata, their last common ancestor and all of its descendants (Fig. 8); membership is identical if the internal relationships of the clade are resolved as Lithobiomorpha + Phylactometria. Monophyly is supported by morphological analyses (Edgecombe and Giribet, 2004), nuclear ribosomal and mitochondrial genes, and their combination with morphology (Giribet and Edgecombe, 2006; Murienne et al., 2010), nuclear protein coding genes (Regier et al., 2010), and transcriptomics (Fernandez et al., 2016, 2014b).

### 30.1. *Fossil specimens*

*Devonobius delta* Shear and Bonamo, 1988. AMNH slide 411–15-AR18, holotype (Fig. 9g), complete head with 15 or 16 trunk segments. We also refer to AMNH slide 4329-AR4 (Fig. 9h).

### 30.2. *Phylogenetic justification*

*D. delta* was resolved in a trichotomy with *Craterostigmus* and Epimorpha in the morphological cladistic analysis of (Edgecombe and Giribet, 2004, Fig. 9), and as sister group to extant Phylactometria when those data were combined with sequence data from four genes (Edgecombe and Giribet, 2004, Fig. 14). Published analyses agree on it being more closely related to Epimorpha than to Lithobiomorpha (Shear and Bonamo, 1988, Fig. 1; Murienne et al., 2010, Fig. 2; Fernandez et al., 2016) and it is accordingly crown Pleurostigmophora.

### 30.3. *Age justification*

*D. delta* occurs in the Middle Devonian Gilboa locality, Schoharie County, New York State, USA. Fossils come from the upper part of the Panther Mountain Formation, dated to the Tioughniogan regional Stage, Givetian in the global time scale. Palynomorphs are consistent with a Givetian age (Richardson et al., 1993). Accordingly, minimum date for the end of the Givetian/base of the Frasnian is applied (382.7 Ma).

Soft maximum as for 26.3.

## 31. Crown Amalpighiata

This clade comprises Lithobiomorpha (stone centipedes) and Epimorpha, their last common ancestor and all of its descendants (Fig. 8). Monophyly is supported by targeted gene sequencing (Regier et al., 2010) and transcriptomics (Fernandez et al., 2016, 2014b).

### 31.1. *Fossil specimens*

*Mazoscolopendra richardsoni* Mundel, 1979. FMNH PE 22936, holotype, nearly complete specimen in siderite concretion (J. T. Haug et al., 2014; Mundel, 1979). We also refer to FMNH PE 29002 (Fig. 9i).

### 31.2. *Phylogenetic justification*

*M. richardsoni* was coded by Fernandez et al. (2016) for its morphological data based on descriptions and figures of Mundel (1979) and Haug et al. (J. T. Haug et al., 2014), and personal observation by G.D.E. of type and other material in the Field Museum. It was resolved as total-group Scolopendromorpha based on possession of autapomorphies of that order (e.g. single tergite on the forcipular/first leg-bearing segments, 21 pairs of trunk legs) but cannot be placed more precisely with regards to whether it is a stem-or crown-group scolopendromorph. Nonetheless, its membership in Scolopendromorpha assigns it to crown Amalpighiata. The same calibration would apply were the putative clade Phylactometria endorsed in lieu of Amalpighiata.

### 31.3. *Age justification*

Specimens are derived from the Francis Creek Shale Member of the Carbondale Formation, Mazon Creek, Illinois, of Westphalian D age (Baird et al., 1985; Shabica and Hay, 1997). The Westphalian D is equivalent to the uppermost Moscovian stage of the Pennsylvanian (Richards, 2013). As the upper boundary of the Moscovian is 307.0 Ma ± 0.1 Myr, this provides a minimum age of 306.9 Ma.

Soft maximum as for 26.3.

## 32. Crown Epimorpha

This clade comprises Scolopendromorpha and Geophilomorpha, their last common ancestor and all of its descendants (Fig. 8). Monophyly is supported by morphological analyses (Fernandez et al., 2016; Murienne et al., 2010) and transcriptomics (Fernandez et al., 2016, 2014b).

### 32.1. *Fossil specimens*

As for 31.1.

### 32.2. *Phylogenetic justification*

As for 31.2.

### 32.3. *Age justification*

As for 31.3.

## 33. Crown Geophilomorpha

This clade comprises Placodesmata and Adesmata, their last common ancestor and all of its descendants (Fig. 8). Monophyly is supported by morphological analyses, targetted sequencing, and combination of molecular and morphological data (Bonato et al., 2014a; Fernandez et al., 2014b; Murienne et al., 2010) and transcriptomics (Fernandez et al., 2016).

### 33.1. *Fossil specimens*

*Kachinophilus pereirai* Bonato et al., 2014b. AMNH Bu-Ba41a, holotype (Fig. 9j,k), complete adult male in amber (Bonato et al., 2014b, Fig. 1A-B, 2).

### 33.2. *Phylogenetic justification*

*K. pereirai* was originally assigned to the extant family Geophilidae based on a combination of characters that is unique to that family. More precisely it compares most closely to a subgroup within Geophilidae that has been classified as subfamilies Chilenophilinae or Pachymeriinae. *K. pereirai* was coded by Fernandez et al. (2016) for its morphological data based on original observations on the type material. It was resolved as more closely related to extant Geophilidae *(sensu* Bonato et al., 2014a) than to members of any of the other sampled geophilomorph family,as predicted by its original classification. Thus it is unambiguously a member of crown Adesmata and accordingly crown Geophilomorpha.

### 33.3. *Age justification*

Minimum as for 12.3. Soft maximum as for 26.3.

### 33.4. *Discussion*

A total-group geophilomorph from the Upper Jurassic of Germany, *Eogeophilus jurassicus* Schweigert and Dietl, 1997 (refigured by J. T. Haug et al., 2014), is too inadequately known to establish whether or not it is a member of the geophilomorph crown-group.

## 34. Crown Pancrustacea

This clade comprises Oligostraca, Multicrustacea, and Allotriocarida, their last common ancestor and all of its descendants (Fig. 1). The inclusion of Hexapoda in a paraphyletic ‘Crustacea’ (and hence, erection of the clade Pancrustacea; Zrzavy and Stys, 1997) has been supported by numerous phylogenetic analyses, including those based on nuclear protein-coding genes (Regier et al., 2010, 2005), transcriptomes (Andrew, 2011; Meusemann et al., 2010; Rota-Stabelli et al., 2011; von Reumont et al., 2012), morphology (Legg et al., 2013; Schram and Koenemann, 2004; Strausfeld and Andrew, 2011) and combined morphological and molecular data (Oakley et al., 2013).

This clade has also been named Tetraconata (Dohle, 2001) referring to the shared apomorphy of four cone cells within the compound eye; however this character is absent in many members of the clade, with multiple possible reconstructions of homology (Oakley, 2003; T. Oakley, pers. comm.). Terminology that does not rely on the homology of cone cell arrangement is thus preferred. More recently, an amended version of ‘Crustacea’ has been proposed (Haug and Haug, 2015) to avoid a different application of the ‘Pan-’ prefix (Lauterbach, 1989). While this concept of Crustacea is in our view valid, for this purpose we favour the original use of Pancrustacea referring to the crown group members only (Zrzavy and Stys, 1997). Haug and Haug (2015) argue that fossils such as Phosphatocopina would need to be included within Pancrustacea, however recent phylogenetic analyses show the sister group to crown Pancrustacea is in fact crown Myriapoda, with all other fossils outside (Legg et al., 2013). Pancrustacea is the clade name implemented in GenBank, and is the most commonly used name among molecular workers.

### 34.1. *Fossil specimens*

As for 4.1.

### 34.2. *Phylogenetic justification*

As for 4.2.

### 34.3. *Age justification*

As for 4.3.

### 34.4. *Discussion*

As for 22.4.

## 35. Crown Oligostraca

This clade comprises Ostracoda (seed shrimp), Branchiura (fish lice), Pentastomida (tongue worms), and Mystacocarida, their last common ancestor and all of its descendants (Fig. 10).

**Fig. 10.**
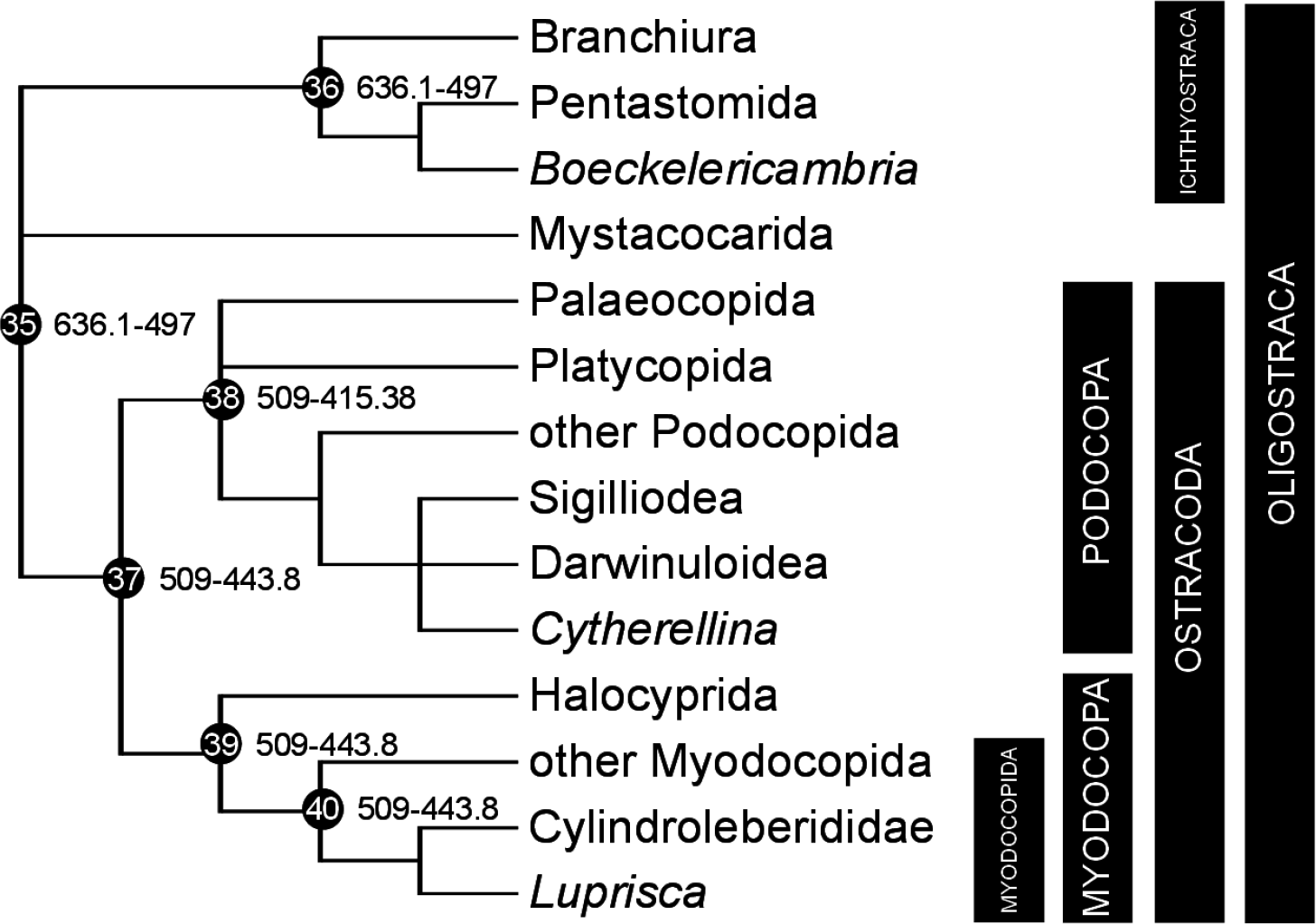
Calibration diagram for Oligostraca (nodes 35–40).

Monophyly of this clade has been demonstrated with nuclear protein-coding genes (Regier et al., 2010; Zwick et al., 2012) and combined phylogenetic analysis of molecules and morphology (Lee et al., 2013; Oakley et al., 2013; Zrzavy et al., 1998). GenBank taxonomy does not recognize this clade. Instead GenBank prefers the Maxillopoda hypothesis (Branchiura, Pentastomida, Mystacocarida, Thecostraca and Copepoda), which has not been recovered in molecular analyses (Abele et al., 1992; Regier et al., 2005) despite support from morphology (Legg et al., 2013).

### 35.1. *Fossil specimens*

*Boeckelericambria pelturae* Walossek and Müller, 1994. UB W116, holotype, consisting of a complete larva (Fig. 11a).

**Fig. 11.**
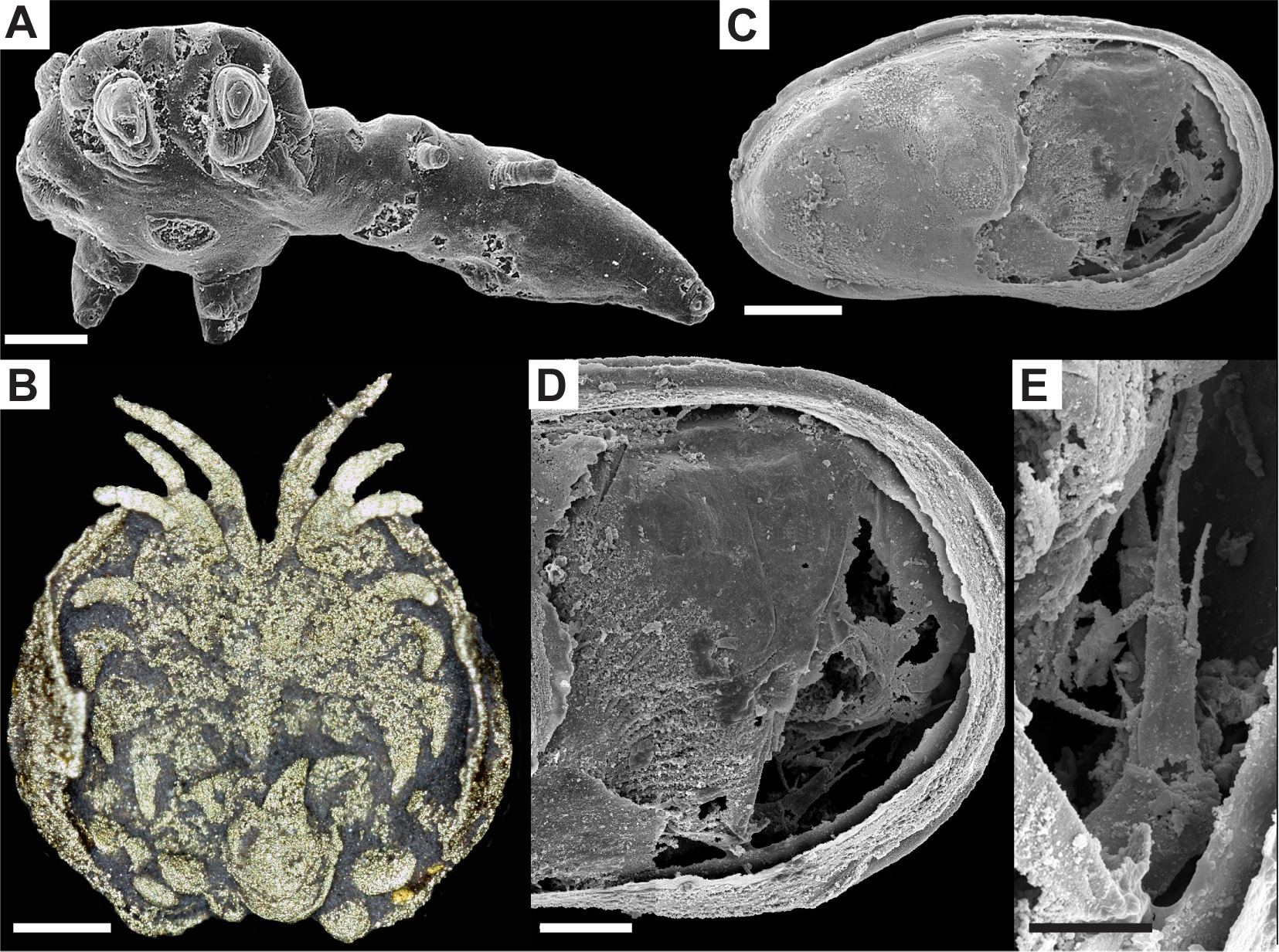
Oligostraca fossil calibrations for (A) nodes 35–36: *Boeckelericambria pelturae,* UB W116, scale bar 50pm, image credit D. Waloszek; (B) nodes 37, 39–40: *Luprisca incuba,* YPM IP 307300, scale bar 500pm, image credit D. Siveter, G. Tanaka, U. Farrell, M. Martin, D. Siveter & D. Briggs; (C-E) node 38: *Cytherellina submagna,* ZPAL 0.60/001, images credit E. Olempska; (C) steinkern left lateral view, scale bar 200 pm; (D) soft anatomy of posterior region, scale bar 100pm; (E) walking legs and presumed furca, scale bar 50pm.

### 35.2. *Phylogenetic justification*

*B. pelturae* is likely a stem group pentastomid, based especially on the diagnostic synapomorphy of a head with two pairs of grasping hooks (similar to the extant *Reighardia* larva; Walossek and Müller, 1994, Fig. 25a). This species is a member of the round headed (as opposed to hammer headed) morphotype (Walossek and Müller, 1994). It was resolved in the pentastomid stem-group in a cladistic analysis that sampled the extant genera by Almeida and Christoffersen (Almeida and Christoffersen, 1999). Its pentastomid identity is not dependent on whether pentastomids are interpreted as Ichthyostraca (Møller et al., 2008; Regier et al., 2010; Sanders, 2010). These studies vary in how well they have adhered to best practices for selecting calibration fossils, as many previous calibrations assume that fossil taxonomy accurately reflects phylogeny. Compounding the issue is the expansion of divergence time studies for a variety of comparative questions far beyond systematics and biogeography, including habitat shifts (Letsch et al., 2016; Lins et al., 2012; Rota-Stabelli et al., 2013a; Yang et al., 2013), genome evolution (Cao et al., 2013; Schwarz et al., 2014; Starrett et al., 2013; Wissler et al., 2013; Yuan et al., 2016), origins of novel characters and behaviours (Rainford et al., 2014; Sanggaard et al., 2014; Wheat and Wahlberg, 2013), evolution of parasites and disease (Ibarra-Cerdena et al., 2014; Palopoli et al., 2014; Rees et al., 2014; Zhou et al., 2014), rate of diversification and its relationship to morphology and ecology (Lee et al., 2013; Wiens et al., 2015), coevolution (Kaltenpoth et al., 2014; Shelomi et al., 2016; Wilson et al., 2013), conservation (Owen et al., 2015), and the use of arthropods as a model for methodological development (O’Reilly et al., 2015; Ronquist et al., 2012; Warnock et al., 2012; Zhang et al., 2016).

### 35.3. *Age justification*

The Orsten fossils come from the lowest zone of the late Cambrian Alum Shale, formally called the *Agnostus pisiformis* Zone or Zone 1, previously corresponding to the Uppermost Zone of the Middle Cambrian (Babcock et al., 2005). The *Agnostus pisiformis* Zone was recently redefined as the uppermost zone of the Guzhangian, at the upper boundary of Cambrian Series 3 (Nielsen et al., 2014). This age of the uppermost stage of the Cambrian Series 3 is 499 Ma ± 2 Myr. Thus the minimum age applied to Oligostraca is 497 Ma.

Soft maximum as for 2.3.

## 36. Crown Ichthyostraca

This clade comprises Branchiura and Pentastomida, their last common ancestor and all of its descendants (Fig. 10). It was first proposed based on sperm ultrastructure (Wingstrand, 1972). Subsequent analyses of morphology combined with protein-coding genes (Zrzavy et al., 1998) or with transcriptomes (Oakley et al., 2013), as well as molecular data on their own (Møller et al., 2008; Regier et al., 2010) have supported monophyly of this clade.

### 36.1. *Fossil specimens*

As for 35.1.

### 36.2. *Phylogenetic justification*

As for 35.2.

### 36.3. *Age justification*

As for 35.3.

## 37. Crown Ostracoda

This clade comprises Myodocopa, Podocopa (including Palaeocopida), their last common ancestor and all its descendants (Fig. 10). Monophyly of this clade has been demonstrated by phylogenetic analysis of a dataset including nuclear protein-coding genes, transcriptomes, and morphology (Oakley et al., 2013). Additional phylogenetic analyses of morphology alone (Legg et al., 2013; Wolfe and Hegna, 2014) also support monophyly.

### 37.1. *Fossil specimens*

*Luprisca incuba* Siveter et al., 2014. YPM IP 307300, holotype, consisting of a complete pyritized specimen in ventral aspect (Fig. 11b).

### 37.2. *Phylogenetic justification*

To date, *L. incuba* is yet to be included in an extensive phylogenetic analysis, but a number of features confirm both its placement within Myodocopida, and therefore Myodocopa. Specifically, the arrangement of setae on the antenula of *L. incuba* is currently only observed amongst extant myodocopid ostracods (Kornicker, 1981).

### 37.3. *Age justification*

The holotype of *L. incuba* was collected from siltstone of the Original Trilobite Bed, Walcott Quarry of Beecher’;s Trilobite Bed, in the Frankfort Shale of upstate New York (Siveter et al., 2014). Beecher’;s site within the Frankfort Shale is within the Lorraine Group, part of the regional Maysvillian Stage of the Cincinnatian Series (Farrell et al., 2011, 2009). Globally, the Maysvillian *(Amplexograptus manitoulinensis* Graptolite Zone) corresponds to the lower Katian Stage, from the base of the *Diplacanthograptus caudatus* Graptolite Zone to the base of the *Pleurograptus linearis* Graptolite Zone (Bergstrom et al., 2009). The upper boundary of the Katian is 445.2 Ma ± 1.4 Myr, providing a minimum age of 443.8 Ma.

As in Oakley et al. (2013), we suggest the maximum age of ostracods must be 509 Ma, the age of the Burgess Shale. Myodocope ostracods possess bivalved, calcified carapaces, which are preserved from many other Burgess Shale arthropods. There is no taphonomic reason why they would not have been preserved from ostracods. The Burgess Shale type locality is from Unit 3 of the Collins Quarry on Mount Stephen in the Canadian Rocky Mountains, British Columbia, which falls within the Kicking Horse Shale Member of the “thick” Stephen Formation (Aitkin, 1997; Caron et al., 2010; Stewart, 1991), also referred to as the Burgess Shale Formation (Fletcher and Collins, 2003, Fletcher and Collins 1998). This unit yields trilobites from the *Polypleuraspis insignis* Subzone of the *Glossopleura* Zone (Fletcher and Collins, 1998), and is the oldest soft-bodied fossil excavation of the Burgess Shale sites. The age of the *Glossopleura* Zone corresponds to the Cambrian Series 3 Stage 5, giving a maximum constraint of 509 Ma.

### 37.4. *Discussion*

Older fossils, from the Tremadocian (~40 Myr older) have been reported from numerous localities across the current and Ordovician world: Argentina, Australia, China, Iran, Norway, Sweden, and the UK (Williams et al., 2008). However, all of these fossils are known solely from carapaces, which are known to be highly homoplastic (Siveter et al., 2013). The affinities of Tremadocian ostracods are therefore ambiguous. Other bivalved crustacean-like taxa from the Cambrian, such as bradoriids and phosphatocopines, are demonstrably not closely related to ostracods, or even Oligostraca, based on phylogenetic analyses (Hou et al., 2010; Legg et al., 2013; Oakley et al., 2013; Wolfe and Hegna, 2014).

## 38. Crown Podocopa

This clade comprises Palaeocopida, Podocopida and Platycopida, their last common ancestor and all of its descendants (Fig. 10). Monophyly has been demonstrated by phylogenetic analysis of morphology (Horne et al., 2005), and morphology combined with nuclear protein-coding genes and transcriptomes (Oakley et al., 2013). Although the sole living representative of Palaeocopida, *Manawa staceyi,* has to date only been represented by a single ribosomal sequence and morphology, bootstrap support for its position as a sister to the remaining Podocopa is strong (Oakley et al., 2013).

### 38.1. *Fossil specimens*

*Cytherellina submagna* Krandijevsky, 1963. For phylogenetically relevant details, we refer to ZPAL O.60/001 (Fig. 11c-e), preserving soft anatomy, and ZPAL O.60/002, preserving adductor muscle scars (Olempska et al., 2012).

### 38.2. *Phylogenetic justification*

*C. submagna* is very similar to modern podocopes, particularly sigilloideans and darwinuloideans, with which it shares a particular adductor muscle scar pattern, long terminal seta on the seventh limb pair, and a furca with large distal setae (Olempska et al., 2012).

### 38.3. *Age justification*

Specimens of *C. submagna* were recovered from two localities in Podolia, Ukraine: Kasperovcy village, left border of the river Seret (type locality), and from the right escarpment of the River Dniester near the village Ivanye Zlote (Olempska et al., 2012). The type locality, part of the Chortkov/Chortkiv Horizon, underlies the second locality, which is part of the Ivanye Horizon (Filipiak et al., 2012; Olempska et al., 2012). Thus we must use the Chortkiv age as a minimum constraint on the age of *C. submagna*, to provide the narrowest interval of clade divergence. The Chortkiv Horizon comfortably lies within the middle Lochkovian stage of the Lower Devonian (Filipiak et al., 2012; Małkowski et al., 2009; Plotnick, 1999). Conodont biostratigraphy places the upper boundary of the Chortkiv Horizon at the end of the *Caudicriodus postwoschmidti* Biozone, the oldest conodont Biozone within the Devonian (Drygant and Szaniawski, 2012). Spline fits on radiometric ages for the Devonian place the *C. postwoschmidti* Biozone at 417.27 Ma with a duration of 1.89 Myr (Becker et al., 2012). Thus the end of the *C. postwoschmidti* Biozone, and a minimum age for the first appearance of *C. submagna*, is 415.38 Ma.

Soft maximum as for 37.3.

### 38.4. *Discussion*

Although isolated ostracod carapace valves are incredibly abundant in the fossil record, the morphology of carapaces has been shown to have little systematic value (Siveter et al., 2013). For this reason we have selected a taxon with soft-tissue preservation, of which *C. submagna* is the oldest recognized example, although other species of *Cytherellina* are known from older deposits including the later Silurian of Ludlow, England, with only the carapaces preserved (Olempska et al., 2012). These older species cannot be ruled out as myodocopes or stem members of any of Podocopa, Myodocopa, or even Ostracoda, as they lack diagnostic soft parts.

## 39. Crown Myodocopa

This clade comprises Myodocopida and Halocyprida, their last common ancestor and all of its descendants (Fig. 10). Monophyly has been demonstrated by phylogenetic analysis of morphology (Horne et al., 2005), and morphology combined with nuclear protein-coding genes and transcriptomes (Oakley et al., 2013).

### 39.1. *Fossil specimens*

As for 37.1.

### 39.2. *Phylogenetic justification*

As for 37.2.

### 39.3. *Age justification*

As for 37.3.

## 40. Crown Myodocopida

This clade comprises Cylindroleberidoidea, Cypridinoidea and Sarsielloidea, their last common ancestor and all of its descendants (Fig. 10). Monophyly has been demonstrated by phylogenetic analysis of morphology (Horne et al., 2005), and morphology combined with nuclear protein-coding genes and transcriptomes (Oakley et al., 2013).

### 40.1. *Fossil specimens*

As for 37.1.

### 40.2. *Phylogenetic justification*

As for 37.2.

### 40.3. *Age justification*

As for 37.3.

## 41. Crown Altocrustacea

This clade comprises Multicrustacea and Allotriocarida, their last common ancestor and all of its descendants (Fig. 1). Monophyly has been supported by phylogenetic analysis of nuclear protein-coding genes (Regier et al., 2010; Zwick et al., 2012), transcriptomes (von Reumont et al., 2012), and combined analysis of morphology and nuclear protein-coding genes (Lee et al., 2013) or morphology and transcriptomes (Oakley et al., 2013). However, this clade has been challenged as paraphyletic (containing Oligostraca) by Rota-Stabelli et al. (2013b), and has not been supported by morphological data alone. It is not recognized in GenBank taxonomy.

### 41.1. *Fossil specimens*

As for 4.1.

### 41.2. *Phylogenetic justification*

As for 4.2.

### 41.3. *Age justification*

As for 4.3.

### 41.4. *Discussion*

A series of disarticulated Small Carbonaceous Fossils (Harvey and Butterfield, 2008) from the early Cambrian Mount Cap Formation, Northwest Territories, Canada, have been cited as calibration fossils within Altocrustacea or even Allotriocarida (e.g. Rehm et al., 2011; Sun et al., 2015). These fossils were argued by Harvey (2008) to comprise part of the feeding apparatus of a single crustacean taxon. The Mount Cap arthropod fossils would have represented structures each specialized for precise feeding functions. The fossil species may have initially scraped food with saw-toothed and hooked setae, further processed particles with filter plates and other delicate setal associations, then macerated with mandibular molar surfaces and passed to the mouth with long fringing setae (Harvey and Butterfield, 2008). Estimates of the body size of the animal, based on regression of body length versus molar surface length (from extant crustaceans) suggest the Mount Cap arthropod was, in total, about the same size as an adult cladoceran or anostracan (Harvey and Butterfield, 2008). While direct synapomorphies linking the Mount Cap arthropod to crown group branchiopods are lacking, the evidence together suggests affinity along the stem lineage of Altocrustacea (Harvey, 2008, Fig. 5.6).

## 42. Crown Multicrustacea

This clade comprises Copepoda, Thecostraca (barnacles) and Malacostraca, their last common ancestor and all of its descendants (Figs. 12 and 13). Monophyly was first demonstrated by nuclear protein-coding genes (Regier et al., 2010) and supported by transcriptomes (von Reumont et al., 2012) and combined analysis of molecular and morphological data (Lee et al., 2013; Oakley et al., 2013). This clade has, however, not been recovered in any morphology-only phylogenetic analyses, presumably owing to widespread support for Malacostraca as sister to much of the rest of Pancrustacea (the Entomostraca hypothesis, e.g. Walossek and Müller, 1998). See Wolfe and Hegna (2014) for a morphological deconstruction of Entomostraca. Neither Multicrustacea nor Entomostraca is recognized in GenBank taxonomy.

**Fig. 12.**
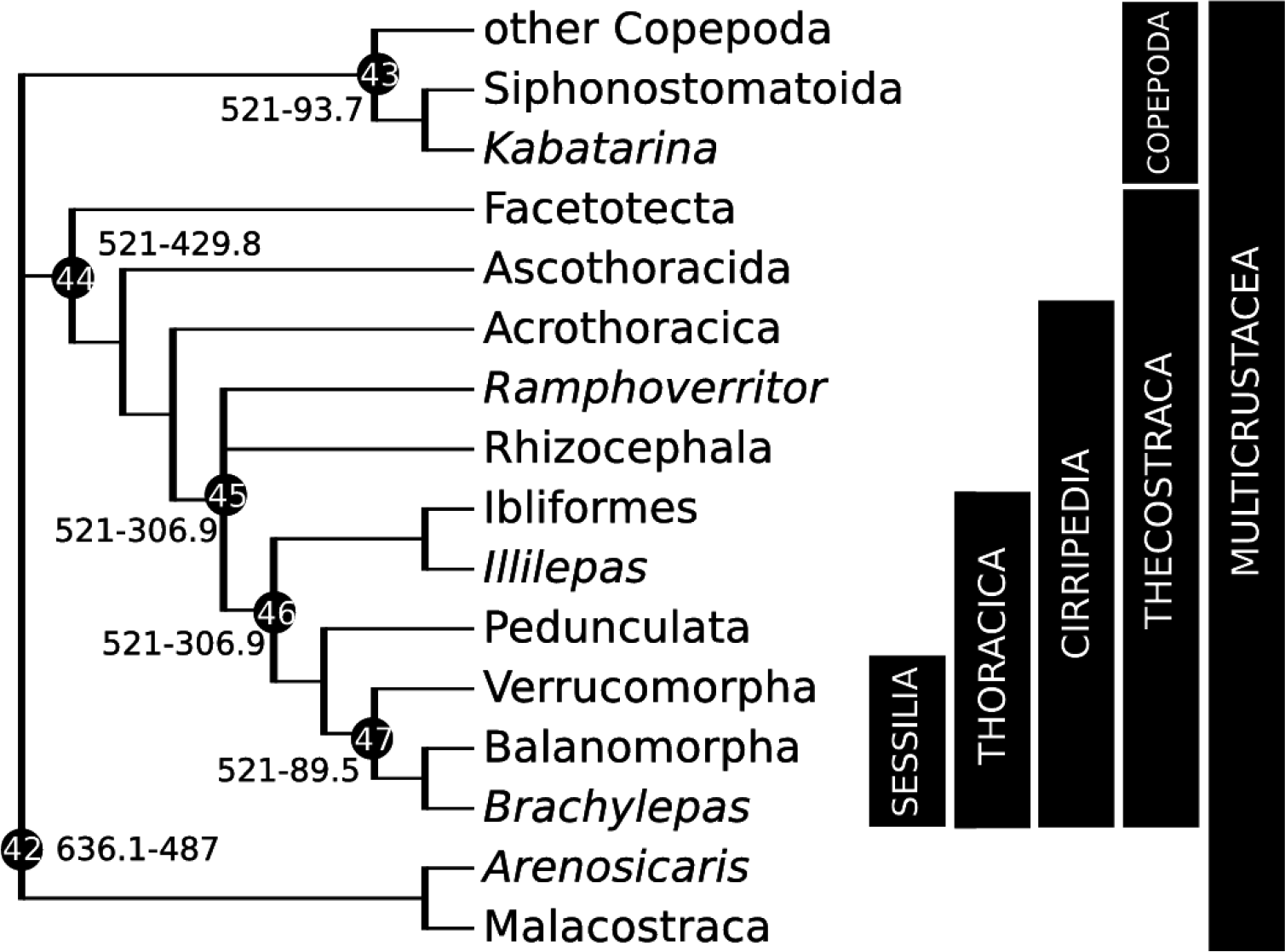
Calibration diagram for Copepoda and Thecostraca (nodes 42–47).

**Fig. 13.**
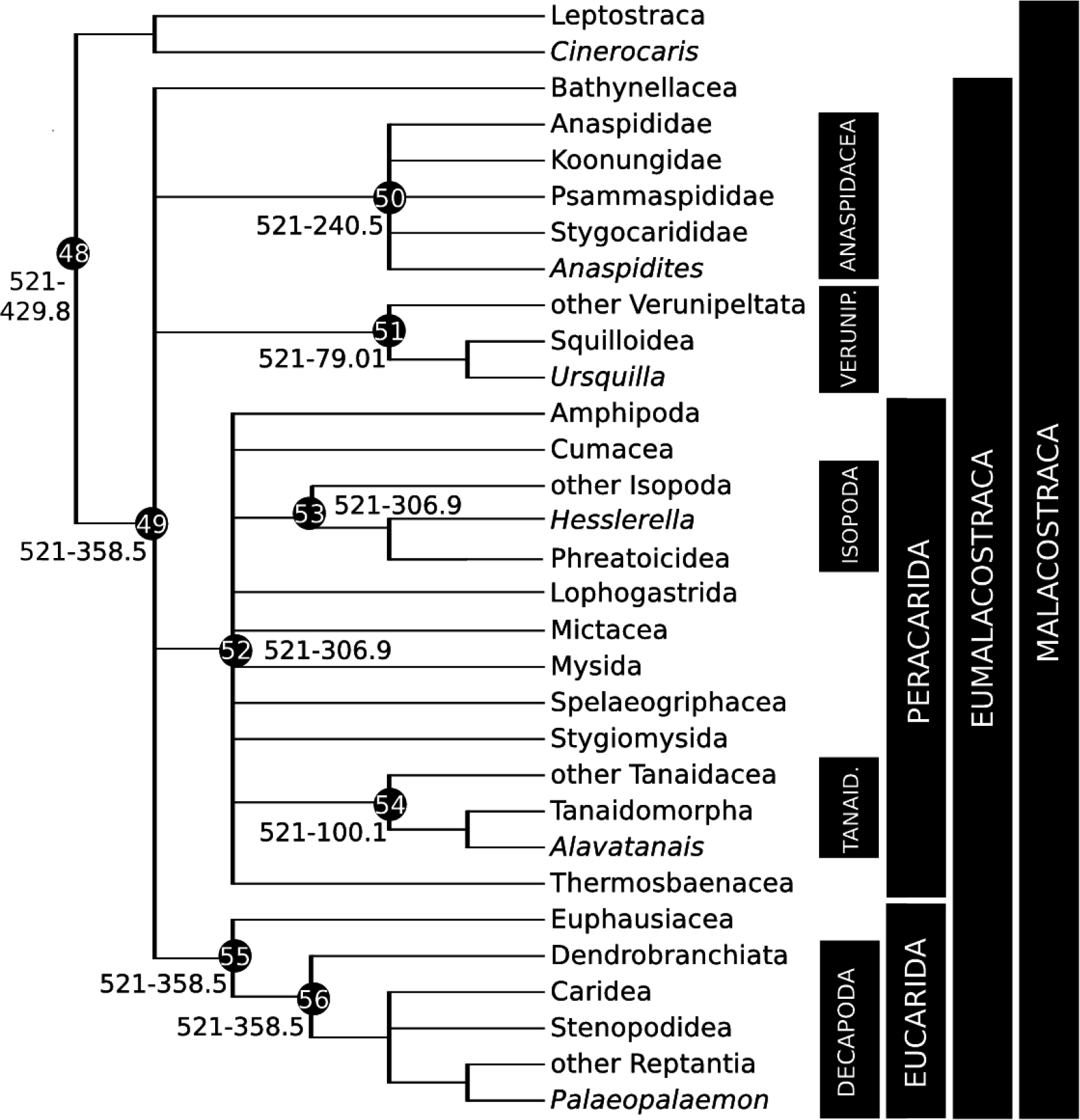
Calibration diagram for Malacostraca (nodes 48–56). Verunip. = Verunipeltata, Tanaid = Tanaidacea.

### 42.1. *Fossil specimens*

*Arenosicaris inflata* Collette and Hagadorn, 2010a. PRI 10130), holotype from the Elk Mound Group (Fig. 14a), which is difficult to date (see 42.3). Therefore, we refer to a second specimen, UWGM 745.

**Fig. 14.**
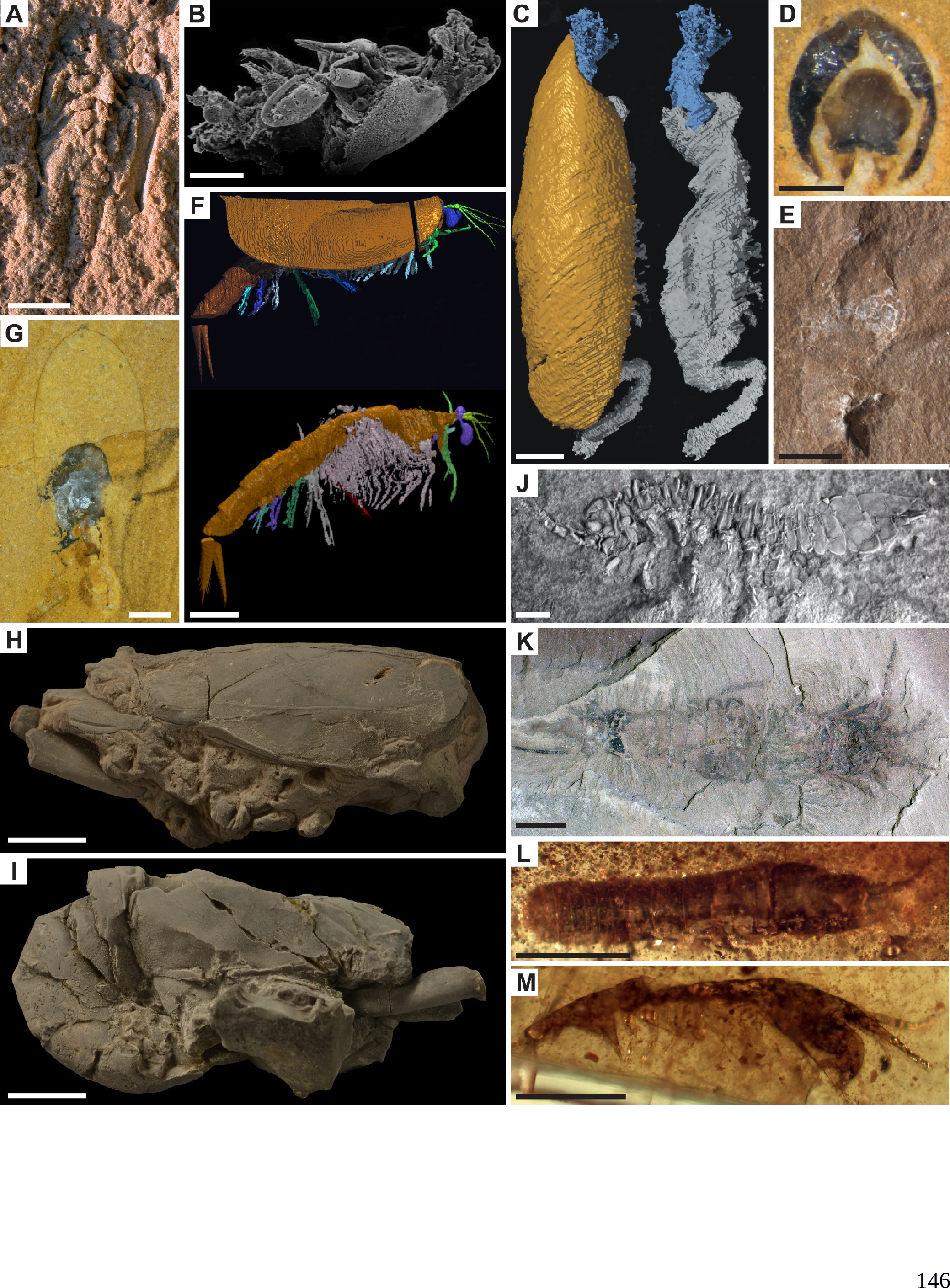
Multicrustacea fossil calibrations for (A) node 42: *Arenosicaris inflata,* PRI 10130, scale bar 10mm, image credit J. Collette; (B) node 43: *Kabatarina pattersoni,* NHMUK 63466, scale bar 100pm, image credit G. Boxshall; (C-D) node 44: *Rhamphoverritor reduncus,* OUM C.29587, scale bars 500pm, image credit D. Briggs, M. Sutton, D. Siveter & D. Siveter; (C) lateral views with (left) and without (right) head shield; (D) transverse section before serial grinding; (E) nodes 45–46: *Illilepas damrowi,* FMNH P32055, scale bar 5mm, image credit J. Wittry; (F-G) node 48: *Cinerocaris magnifica,* images credit D. Briggs, M. Sutton, S. Siveter & D. Siveter; (F) OUM C.29566, reconstruction in lateral view with (top) and without (bottom) head shield, scale bar 2mm; (G) holotype 29565, sub-transverse section, scale bar 1mm; (H-I) nodes 49, 55–56: *Palaeopalaemon newberryi,* KSU 3484, scale bars 5mm, image credit W. Jones; (H) left view; (I) right view; (J) nodes 52–53: *Hesslerella shermani,* FMNH PE 16527, latex cast whitened with ammonium chloride, scale bar image credit T. Hegna; (K) node 50: *Anaspidites antiquus*, AMS F64765, scale bar 5mm, image credit S. Ahyong; (L-M) node 54: *Alavatanais carabe,* scale bars 500 pm, images credit A. Sanchez-Garcia; (L) holotype MCNA 9537; (M) MCNA 13888 lateral view.

### 42.2. *Phylogenetic justification*

*A inflata* was identified within the Archaeostraca, the likely fossil sister group to extant Leptostraca (together comprising ‘Phyllocarida’) and included in the morphological cladistic analysis of Collette and Hagadorn (2010b). In that analysis, the exceptionally preserved fossils *Nahecaris stuertzi* Jaeckel, 1921 and *Cinerocaris magnifica* Briggs et al., 2004 were also included within Archaeostraca (Collette and Hagadorn, 2010b). A separate extensive combined molecular and morphological analysis placed *C. magnifica* within crown Malacostraca (as is traditionally assumed for members of Archaeostraca), while *N. stuertzi* was equivocally stem Leptostraca or stem Malacostraca (Oakley et al., 2013), suggesting non-monophyly of Archaeostraca. *C. magnifica* was also crown Malacostraca in another morphological analysis that omitted *N. stuertzi* (Legg et al., 2013). *A inflata* is within the same archaeostracan clade as *N. stuertzi* (Pephricaridina + Rhinocaridina) while *C. magnifica* is in a separate archaeostracan group (Echinocaridina; Collette and Hagadorn, 2010b). Given the uncertainty of crown affinities and potential monophyly of Archaeostraca, we conservatively assign *A inflata* to crown-group Multicrustacea, but not Malacostraca.

### 42.3. *Age justification*

*A inflata* is found in the Elk Mound Group (holotype) and in the Lodi Member of the St. Lawrence Formation, both in Wisconsin (Collette and Hagadorn, 2010a). Although the Elk Mound Group is the older of these, no biostratigraphically useful fossils co-occur with *A. inflata,* limiting the ability to determine the formation to which they belong (Collette and Hagadorn, 2010a). The St. Lawrence Formation is younger, containing *Saukia* Zone trilobites, which mark it as late Sunwaptan within the Furongian (Collette and Hagadorn, 2010a; Raasch, 1951). The Sunwaptan is the second latest stage of the Furongian, postdated by the Skullrockian (which extends into the Early Ordovician; Peng et al., 2012). The Sunwaptan-Skullrockian boundary is determined by the appearance of conodonts in the *Hirsutodontus hirsutus* Subzone of the *Cordylodus proavus* Zone (Peng et al., 2012). Based on the correlation diagram of Peng et al. (2012), the Sunwaptan-Skullrockian boundary is approximately 487 Ma, providing a minimum age estimate.

Soft maximum as for 2.3.

## 43. Crown Copepoda

This clade comprises Calanoida, Cyclopoida, Gelyelloida, Harpacticoida (benthic copepods), Misophrioida, Mormonilloida, Platycopioida and Siphonostomatoida (fish parasites), their last common ancestor and all of its descendants (Fig. 12). Members of Calanoida, Cyclopoida, and Siphonostomatoida were included in a large combined analysis of transcriptomes and morphology, forming a monophyletic group (Oakley et al., 2013). Phylogenetic analysis of morphology (Huys and Boxshall, 1991; Ho, 1994), mitochondrial genes (Minxiao et al., 2011), and ribosomal genes combined with morphology (Huys et al., 2007) suggest this sampling covers distant lineages of Copepoda, although omitting Platycopioida, the presumed most basal order (Huys and Boxshall, 1991). Molecular data remain unavailable from Platycopioida, although comparative morphological investigations support copepod monophyly (Dahms, 2004).

### 43.1. *Fossil specimens*

*Kabatarina pattersoni* Cressey and Boxshall, 1989. BMNH IN. 63466, holotype, preserving the cephalothorax, mouthparts, oral cone, and first and second thoracic limbs (Fig. 14b). This specimen likely represents an adult female, recovered from the gills of a fossil teleost fish (Cressey and Boxshall, 1989).

### 43.2. *Phylogenetic justification*

*K. pattersoni* has not been included in a formal phylogenetic analysis. Cressey and Boxshall (1989) detail one apomorphy shared by the fossil and recent members of the family Dichelesthiidae, which is a medial groove delimiting the distal part of the maxillary claw. A number of other characters are shared more generally with copepods, such as the shape of the oral cone (typical for fish parasitic copepods), and biramous thoracic limbs with a 2-segmented protopod, joined by the intercoxal plate (Cressey and Boxshall, 1989). Dichelesthiidae is a family belonging to the Siphonostomatoida. Even a position for *K. pattersoni* on the stem of Dichelesthiidae or stem Siphonostomatoida would be within crown group Copepoda.

### 43.3. *Age justification*

*K. pattersoni* was found in the Cretaceous Santana Formation, Serra do Araripe, Ceara, Brazil (Cressey and Boxshall, 1989), which is mainly famous for concretions enclosing fossil fishes. The age of the Santana Formation is poorly constrained (as it lacks biostratigraphic index fossils and igneous rocks for radiometric dating); dates have been suggested that range from the Aptian to the Albian or Cenomanian (Martill, 2007). In order to ensure a minimum date, the upper boundary of the Cenomanian, which is 93.7 Ma (from 93.9 Ma ± 0.2 Myr; Ogg et al., 2012), is used.

Soft maximum as for 26.3.

### 43.4. *Discussion*

Despite their overwhelming abundance in modern aquatic ecosystems, copepods have a poor fossil record, likely due to their small size, unsclerotized cuticle, and planktonic ecology. Apart from *K. pattersoni*, putative copepod appendages have been reported from much older sediments in the Pennsylvanian (Selden et al., 2010) and even the Cambrian (Harvey et al., 2012; Harvey and Pedder, 2013; These fossils each bear one to four characters found in crown copepods, from partial maxillae (Selden et al., 2010) and partial or complete mandibular gnathal edges (Harvey et al., 2012; Harvey and Pedder, 2013). Relationships between feeding habits and mandibular morphology have been observed in extant copepods (Michels and Schnack-Schiel, 2005), and variation may occur among closely related species (Sano et al., 2015). It is therefore extremely unlikely that mandibular characters have not experienced any homoplasy since the Cambrian.

Nevertheless, divergence time analyses suggest Devonian-Carboniferous (calibrated with *K. pattersoni;* (Rota-Stabelli et al., 2013a), Carboniferous (calibrated with external fossils from other pancrustacean clades; Oakley et al., 2013), or Permian (external fossils; Wheat and Wahlberg, 2013) origins for crown Copepoda. These analyses do not conflict stratigraphically with a crown assignment for the Pennsylvanian fossils (Selden et al., 2010). Therefore we must caution that a Cretaceous age is likely a severe underestimate of crown copepod antiquity.

## 44. Crown Thecostraca

This clade comprises Facetotecta (y-larvae), Ascothoracida, and Cirripedia, their last common ancestor and all of its descendants (Fig. 12). Monophyly of Thecostraca has been demonstrated by separate analysis of nuclear housekeeping genes and morphology (Perez-Losada et al., 2009a). All included clades have complex (sometimes only partially known) life histories, but particularly strong morphological support comes from a shared larval stage, the cyprid.

More recently, the enigmatic parasitic Tantulocarida were added within Thecostraca based on analysis of a single ribosomal gene (Petrunina et al., 2014), however, other relationships within Pancrustacea differed significantly from those outlined herein. In light of the paucity of data other than ribosomal genes, we remain ambivalent about including Tantulocarida. As tantulocarids have no known fossil record, if further evidence supports their position within Thecostraca, this calibration may be modified to include them as well.

### 44.1. *Fossil specimens*

*Rhamphoverritor reduncus* Briggs et al., 2005. OUM C.29587, holotype (Fig. 14c,d), preserving a cyprid larva in a volcaniclastic concretion. As the reconstruction of Herefordshire fossils requires serial grinding and photography of 20 μm sections (Sutton et al., 2002), the holotype (figured in Briggs et al., 2005: Fig. 1) was thus destroyed in preparation. Morphological data for Herefordshire fossils are published as 3D models of the thin section photographs.

### 44.2. *Phylogenetic justification*

In a phylogenetic analysis of morphology, *R. reduncus* is the sister group of two species of Thoracica (both are members of Balanomorpha) (Legg et al., 2013). This is a position likely within the crown group of Thecostraca, however, no other thecostracans (such as Facetotecta and Ascothoracida) were included. Generally, the cyprid larval morphotype (with an elongated head shield, six swimming thoracopods, and robust modified antennules) is considered a synapomorphy of Thecostraca (Høeg et al., 2004). *R. reduncus* differs from cirripede cyprids as it lacks attachment discs, and its abdomen extends past the head shield; a differentiated abdomen is a condition of Ascothoracida (Briggs et al., 2005; Høeg et al., 2009a). Based on the presence of five shell plates in a juvenile specimen, Høeg et al. (2009b) suggested that *R. reduncus* may be placed on the cirripede stem lineage. In combination with the phylogenetic analysis of Legg et al. (2013), these apomorphies indicate that *R. reduncus* is very likely a member of crown group Thecostraca, and likely on the stem of Cirripedia.

### 44.3. *Age justification*

Minimum as for 6.3. Soft maximum as for 26.3.

## 45. Crown Cirripedia

This clade comprises Acrothoracica, Rhizocephala and Thoracica (barnacles), their last common ancestor and all of its descendants (Fig. 12). Monophyly has been demonstrated by separate phylogenetic analysis of nuclear housekeeping genes and morphology (Perez-Losada et al., 2009a).

### 45.1. *Fossil specimens*

*Illilepas damrowi* Schram, 1986. FMNH P32055, holotype (Fig. 14e).

### 45.2. *Phylogenetic justification*

Schram (1975) described this fossil as *Praelepas damrowi*, a congener of *P. jaworskii* Chernyshev, 1930. Restudy led to the transfer of this fossil to the new genus *Illilepas* under the combination *I. damrowi*, as the original description of a carina more closely resembled a tergum, and the original tergum was more likely an enlarged spine along the margin of the aperture, similar to that seen in Ibliformes (Thoracica) (Buckeridge and Newman, 2006; Schram, 1986). However, both molecular and morphological data place *Ibla* as the most basal clade of Thoracica (Perez-Losada et al., 2009a, 2008), which, if *I. damrowi* is on the Ibliformes stem lineage, would still situate the fossil within the crown group of Cirripedia.

### 45.3. *Age justification*

As for 31.3.

### 45.4. *Discussion*

A possible Lower Ordovician stalked barnacle (Pedunculata: Lepadomorpha?) was illustrated in Fig. 2c and S3h of Van Roy et al. (2010). It has not been formally described and its affinities are unclear.

*P. jaworskii* is in some ways a more appropriate fossil calibration than *I. damrowi.* It is approximately coeval to *I. damrowi,* and was coded in a morphological phylogenetic analysis (Glenner et al., 1995), where it was placed within Thoracica (sister to all other Thoracica except *Ibla). P. jaworskii* has been used as a calibration fossil for Thoracica and Cirripedia (Perez-Losada et al., 2009b, 2008; Rees et al., 2014). However, no specimen information was available in the original publication, nor was any significant stratigraphic data beyond Carboniferous of the Kusnetzk Basin, Russia (Chernyshev, 1930). As the papers using *P. jaworskii* for calibration estimate its age at 306.5–311.7 Ma (e.g. Rees et al., 2014), substitution of the slightly younger *I. damrowi* will not significantly violate the minimum age.

## 46. Crown Thoracica

This clade comprises Ibliformes, ‘Pedunculata’ (goose barnacles) and Sessilia (acorn barnacles), their last common ancestor and all of its descendants (Fig. 12). Pedunculata is no longer supported as monophyletic (Buckeridge and Newman, 2006; Perez-Losada et al., 2009a; Rees et al., 2014). Ibliformes is identified as the sister group of all other Thoracica (Perez-Losada et al., 2009a). Monophyly has been established by phylogenetic analysis of nuclear housekeeping genes (Perez-Losada et al., 2009a), although a morphological phylogenetic analysis in the same paper resolves all studied members of Rhizocephala and Thoracica together in a polytomy. This is because only larval characters can be scored for Rhizocephala, with missing data for all adult characters due to their parasitic lifestyle.

### 46.1. *Fossil specimens*

As for 45.1.

### 46.2. *Phylogenetic justification*

As for 45.2.

### 46.3. *Age justification*

As for 31.3.

### 46.4. *Discussion*

As for 45.4.

## 47. Crown Sessilia

This clade comprises Verrucomorpha and Balanomorpha, their last common ancestor and all of its descendants (Fig. 12). Monophyly is supported by phylogenetic analysis of nuclear protein-coding and ribosomal genes (Perez-Losada et al., 2008; Rees et al., 2014).

### 47.1. *Fossil specimens*

*Brachylepas fallax* Withers, 1935. For calibration, we refer to the stratigraphically oldest specimen (SM 704275), which is undescribed (A. Gale, pers. comm.).

### 47.2. *Phylogenetic justification*

This species was originally described from disarticulated material by Darwin (1851) as *Pollicipes fallax.* Withers (1935, 1914) recognized it was a sessile, rather than pedunculate barnacle, and that it had similarities to the basal sessilian genus *Pycnolepas* in overall form and plate development. Synapomorphies shared by *B. fallax* and all crown Sessilia include: the absence of a peduncle, presence of an operculum, and absence of all lateral plates (Gale and Sørensen, 2015). *B. fallax* shares with crown Balanomorpha a low, hemiconical carina and rostrum (Gale and Sørensen, 2015). On the basis of these characters, a recent cladogram depicts *B. fallax* as one of the most distant stem lineages of Balanomorpha, which is therefore a position within the crown group of Sessilia (Gale and Sørensen, 2015).

### 47.3. *Age justification*

The oldest known locality from which *B. fallax* has been recovered is Pit No. 125 (Brydone, 1912), close to junction of Barnet Side Lande and King’;s Lane, Froxfield, Hampshire, England (A. Gale, pers. comm.). This locality bears fossils of *Holaster (Sternotaxis) planus,* and is thus part of the *S. planus* echinoid zone, which is the uppermost zone of the Turonian in English Chalk (Gale, 1996; Mortimore, 2011). The GSSP defining the global upper boundary of the Turonian remains debated (due to difficulty in identifying its index ammonite fossil, *Forresteria (Harleites) petrocoriensis* (Ogg et al., 2012). Currently, the upper boundary of the Turonian is dated to 89.8 Ma ± 0.3 Myr, providing a minimum age of 89.5 Ma.

Soft maximum as for 26.3.

### 47.4. *Discussion*

The Albian-Cenomanian *Verruca withersi* Schram and Newman, 1980 has been shown not to be a cirripede (Jagt and Buckeridge, 2005) and cannot be used as a minimum. *Proverruca* (coded at the genus level) was included in a morphological phylogenetic analysis, where it was placed in a polytomy with the fossil *Eoverruca* and the crown family Verrucidae (Glenner et al., 1995), and subsequently used as a calibration fossil for divergence time analysis (Perez-Losada et al., 2014, Perez-Losada, 2008). However, the placement of *Proverruca* was based on similarities to the extant genus *Neoverruca*, which was shown in molecular analyses to fall outside Sessilia entirely, instead within the Scalpelliformes (Perez-Losada et al., 2008; Rees et al., 2014). More recent morphological phylogenetic analyses confirm the convergence between *Neoverruca* + *Proverruca* and crown Verrucomorpha (Gale, 2015; Gale and Sørensen, 2015).

The Albian *Pycnolepas rigida* Sowerby, 1836 was included in a morphological phylogenetic analysis, where it was placed on the stem lineage of Verrucomorpha (Gale, 2015). However, that analysis did not include members of Balanomorpha, so the topology did not explicitly exclude a position on the stem lineage of Sessilia. In fact, this species was referred to stem Sessilia (Gale, 2015, p. 770). Unlike crown group Sessilia, it retains the pedunculate character of a lateral plate, the tall upper latus (Gale, 2015).

*Verruca tasmanica tasmanica* (Buckeridge, 1983), a previously used calibration fossil at the base of Verrucomorpha (Herrera et al., 2015; Linse et al., 2013; Perez-Losada et al., 2014, Perez-Losada, 2008; Rees et al., 2014), is known from the Santonian Gingin Chalk Formation of Dandaragan, Western Australia (as well as type material from younger Oligocene strata of Oamaru, New Zealand; Buckeridge, 1979). As it is both younger than *B. fallax* and has not been studied in a phylogenetic context, it is not used herein.

## 48. Crown Malacostraca

This clade comprises Leptostraca and Eumalacostraca, their last common ancestor and all of its descendants (Fig. 13). Its monophyly is one of the least contested matters in arthropod phylogeny; it has been demonstrated by phylogenetic analysis of morphology (Legg et al., 2013; Wills et al., 1998), nuclear ribosomal and protein-coding genes (Giribet et al., 2001; Regier et al., 2010, 2005), transcriptomes (von Reumont et al., 2012), and combined analysis of molecular and morphological data (Lee et al., 2013; Oakley et al., 2013).

### 48.1. *Fossil specimens*

*Cinerocaris magnifica* Briggs et al., 2004. OUM C.29565 (holotype; Fig. 14g), and OUM C.29566 (serially ground and reconstructed specimen; Fig. 14f). Morphological data for Herefordshire fossils are published as 3D models of thin section photographs.

### 48.2. *Phylogenetic justification*

*C. magnifica* was found to be a member of the stem group of Leptostraca (therefore crown group Malacostraca) in analyses of morphology (Legg et al., 2013) and morphology combined with molecules (Oakley et al., 2013).

### 48.3. *Age justification*

As for 44.3.

### 48.4. *Discussion*

The position of other fossil phyllocarids with respect to extant malacostracans (and the monophyly of phyllocarids themselves) have not been significantly investigated. The position of the Devonian phyllocarid *Nahecaris stuertzi* in a phylogenetic analysis (Oakley et al., 2013) was equivocally stem Leptostraca or stem Malacostraca, casting doubt on the position of at least rhinocarid phyllocarids within crown Malacostraca. As *N. stuertzi* has the same relationship to crown Malacostraca as *A inflata* (at least in the analysis of Collette and Hagadorn, 2010b), neither fossil can be confidently placed within crown-group Malacostraca. Recent reinvestigation of *Ceratiocaris* cf. *macroura* (related to *A. inflata* and *N. stuertzi* in the analysis of Collette and Hagadorn, 2010b) suggests this Silurian ‘phyllocarid’ may be a stem eumalacostracan due to possession of an antennal scale, casting further doubt on the monophyly of fossil phyllocarids (Jones et al., 2015). The *C. macroura* study, however, assumes malacostracan identity of phyllocarids (partly defined by biramous antennules), which, as discussed above (section 42.2), may not be robust to phylogenetic analysis. If, however, *Ceratiocaris* is within crown Malacostraca (either alone or with other archaeostracans), the oldest Malacostraca would be amended to *Ceratiocaris winneshiekensis* Briggs et al., 2015 from the Darriwilian Winneshiek Lagerstätte of Iowa (~14 Myr older than *C. magnifica).*

Thylacocephalans are an enigmatic fossil arthropod clade ranging from the Silurian to the Cretaceous (C. Haug et al., 2014; Schram, 2014). Some Cambrian fossils have been proposed as thylacocephalans, but their membership is generally discounted (Schram, 2014; Vannier et al., 2006). Thylacocephalans have been compared to several extant arthropod clades, including the malacostracan stomatopods and decapods (Schram, 2014; Schram et al., 1999; Secretan and Riou, 1983; Vannier et al., 2016), which would extend the minimum age of Malacostraca slightly older within the Silurian (~433 Ma). The Jurassic thylacocephalan *Dollocaris ingens* Van Straelen, 1923 was coded in a morphological matrix, and found to be a stem eumalacostracan (therefore crown malacostracan) by phylogenetic analysis (Legg et al., 2013). However, their malacostracan affinities have been questioned, especially by C. Haug et al. (2014) studying a Silurian species, noting divergent trunk tagmosis and similarities to remipedes. Continued uncertainty over thylacocephalan affinites make them poor calibration fossils.

## 49. Crown Eumalacostraca

This clade comprises Verunipeltata (mantis shrimp, partial total group called Stomatopoda), Peracarida, ‘Syncarida’ (itself comprising Anaspidacea and Bathynellacea) and Eucarida, their last common ancestor and all of its descendants (Fig. 13). Monophyly is demonstrated by phylogenetic analysis of morphology (Legg et al., 2013; Richter and Scholtz, 2001), nuclear ribosomal and protein-coding genes (Regier et al., 2010), transcriptomes (von Reumont et al., 2012), and combined analysis of molecular and morphological data (Lee et al., 2013; Oakley et al., 2013; Wills et al., 2009). Although stomatopods and/or syncarids were not sampled in some of the above analyses, the best taxon sampling still resulted in eumalacostracan monophyly (Legg et al., 2013; Wills et al., 2009).

### 49.1. *Fossil specimens*

*Palaeopalaemon newberryi* Whitfield, 1880. The holotype at the AMNH, figured by Schram et al. (1978: Plate 3 #1–3), has been lost (B. Hussaini, pers. comm.), thus we refer to specimen KSU 3484 (Fig. 14h,i).

### 49.2. *Phylogenetic justification*

Schram and Dixon (2004) coded *P. newberryi* into the morphological matrix of Dixon et al. (2003), finding it sister to a clade including Anomura, Brachyura, and Achelata to the exclusion of Astacidea, Axiidea, Gebiidea and Glypheidea. This position is within the crown group of Reptantia, hence within the crown groups of Decapoda and Eumalacostraca.

### 49.3. *Age justification*

The specimen of *P. newberryi* was found in gray shale near “Paine’;s Creek,” LeRoy, Lake County, Chagrin (Erie) Shale, northeastern Ohio, USA (Feldmann et al., 1978; Schram et al., 1978). The Chagrin Shale is dated to the late Fammenian based on presence of the index alga *Foerstia* (Murphy, 1973), which in Ohio lies 40–70 m below surface outcrops (Feldmann et al., 1978). The upper boundary of the Fammenian is 358.9 Ma ± 0.4 Myr, giving a minimum of 358.5 Ma.

Soft maximum as for 26.3.

## 50. Crown Anaspidacea

This clade comprises the families Anaspididae, Koonungidae, Psammaspididae and Stygocarididae, their last common ancestor and all of its descendants (Fig. 13). Phylogenetic studies including Anaspidacea are extremely rare, but morphology (Schram, 1984) and mitochondrial 16S sequences (Camacho et al., 2002) indicated monophyly of the clade.

### 50.1. *Fossil specimens*

*Anaspidites antiquus* Chilton, 1929. AMS F64765, holotype, complete specimen (Fig. 14k).

### 50.2. *Phylogenetic justification*

Although not included in a formal phylogenetic analysis, Schram (1984) justified the membership of *A. antiquus* as essentially indistinguishable from living Anaspididacea. Fusion of the first thoracomere into the cephalon, uniramous pleopods, and absence of an antennal scale are noted as diagnostic characters (Schram, 1984). As the fossil lacks preservation of diagnostic mouthparts, exact family affinities within total-group Anaspidacea are uncertain.

### 50.3. *Age justification*

*A antiquus* was found in the Hawkesbury Sandstone at the former Brookvale Brick Quarry, New South Wales, Australia (Schram, 1984). The Hawkesbury Sandstone overlies the Narrabeen Group and underlies the Wianamatta Group (Herbert, 1997). Sequence stratigraphy places the Hawkesbury in Sequence F, including the appearance of *Aratrisporites parvispinosus* spores (Helby, 1973; Herbert, 1997). The *A parvispinosus* spore zone indicates an age during the Anisian (middle Triassic) (Herbert, 1997). The younger bound of the Anisian is estimated at 241.5 ± 1 Ma (Ogg, 2012), providing a minimum age at 240.5 Ma.

Soft maximum as for 26.3.

### 50.4. *Discussion*

‘Syncarida’, the traditional taxon containing Anaspidacea, is purposely excluded from this review, as its monophyly has been substantially challenged. In phylogenetic analyses, syncarid monophyly was only supported by the morphological dataset of Richter and Scholtz (2001), emphasizing the absence of a dorsal carapace in Anaspidacea and Bathynellacea (‘Syncarida’). Each of a partial mitochondrial gene (Camacho et al., 2002), two mitochondrial genes and morphology (Jenner et al., 2009) and morphology including putative syncarid fossils (Schram and Hof, 1998; Wills et al., 2009) failed to recover a sister group relationship bewteen Anaspidacea and Bathynellacea. Wills et al. (2009) suggested that a paraphyletic grade of ‘Syncarida’, including fossils *(Acanthotelson, Palaeocaris*) represented the first divergences of Eumalacostraca.

## 51. Crown Verunipeltata

This is the clade comprising Bathysquilloidea, Erythrosquilloidea, Eurysquilloidea, Parasquilloidea, Gonodactyloidea, Lysiosquilloidea and Squilloidea, their last common ancestor and all of its descendants (Fig. 13). Although the extant members are often referred to Stomatopoda, that clade includes a number of extinct members forming a paraphyletic grade (Fig. 1 of Haug et al., 2010). As membership of fossil species within a crown group is only possible if they branch along the stem lineage of a living clade that is part of the crown itself, we agree with the assertion that crown ‘Stomatopoda’ is equivalent to the clade Verunipeltata *sensu* Haug et al. (2010). The stomatopod clade as a whole may be defined by several apomorphies, such as five pairs of maxillipeds, and modification of the first two pleopods as gonopods (Richter and Scholtz, 2001).

All living members form a clade in analyses of morphology including both fossil and extant taxa (Ahyong, 1997; Ahyong and Harling, 2000). Analysis of combined molecular and morphological data, with limited sampling of verunipeltatan clades, strongly supports monophyly of those members (Jenner et al., 2009). As well, representatives of Gonodactyloidea, Lysiosquilloidea and Squilloidea were sequenced for six housekeeping genes; these were monophyletic in a diverse sample of Pancrustacea (Bybee et al., 2011).

### 51.1. *Fossil specimens*

*Ursquilla yehoachi* Remy and Avnimelech, 1955. For calibration, we refer to two specimens. Based on new phylogenetically relevant details, we use a new specimen (SMNS 67703) from Fig. 1 and 2 of Haug et al. (2013). The SMNS specimen, however, lacks locality and stratigraphic information beyond “Upper Cretaceous Negev Desert, Israel” as it was privately donated (Haug et al., 2013). We also therefore refer to the holotype (MNHN R. 62691).

### 51.2. *Phylogenetic justification*

The uropod morphology of *U. yehoachi* indicates its membership in the crown group of Verunipeltata. The uropodal exopod of *U. yehoachi* specimen SMNS 67703 is bipartite, a synapomorphy of Verunipeltata (Haug et al., 2013). Furthermore, *U. yehoachi* shares several characters with Squillidae (Squilloidea), such as prelateral lobes, submedian teeth with fixed apices and a subquadrate telson (Haug et al., 2013). As *U. yehoachi* has not been included in a phylogenetic analysis, it is uncertain whether it falls within crown Squilloidea (Haug et al., 2013) or on its stem; either position would remain within the crown group of Verunipeltata.

### 51.3. *Age justification*

The holotype of *U. yehoachi* was collected from a chert bank in the Chert Member of the Mishash Formation, near the city of Arad, Israel (Hof, 1998). The chert banks are within the *Hoplitoplacenticeras marroti* ammonite biiozone (Reiss et al., 1986). *H. marroti* co-occurs with *Baculites* sp. (smooth) (Lehmann and Murphy, 2001), which is dated to the uppermost early Campanian. Based on a spline-fit of interbedded bentonites, the base of the Tethyan *Baculites* sp. (smooth) Zone is dated to 79.64 Ma, with a duration of 0.63 Myr (Ogg et al., 2012), thus the minimum age of *U. yehoachi* is 79.01 Ma.

Soft maximum as for 26.3.

## 52. Crown Peracarida

This clade comprises Amphipoda (scuds/beach fleas), Isopoda (wood lice/pill bugs), Cumacea, Lophogastrida, Bochusacea, Mictacea, Mysida (opossum shrimp), Stygiomysida, Spelaeogriphacea, Tanaidacea and Thermosbaenacea, their last common ancestor and all of its descendants (Fig. 13). Monophyly has been demonstrated by phylogenetic analysis of morphology (Jones et al., 2016; Poore, 2005; Richter and Scholtz, 2001; Wills et al., 2009; Wirkner and Richter, 2010), and combined morphology and molecular data (Jenner et al., 2009). Molecular phylogenies based on ribosomal genes reject the inclusion of Mysida within Peracarida (Jenner et al., 2009; Meland and Willassen, 2007; Spears et al., 2005), while Lophogastrida and Stygiomysida are removed from Mysidacea and found comfortably within Peracarida (Meland and Willassen, 2007; Meland et al., 2015). Note ribosomal genes alone are insufficient markers for deep divergences (Giribet and Ribera, 2000), so we cautiously include Mysida within Peracarida pending multilocus investigations.

### 52.1. *Fossil specimens*

*Hesslerella shermani* Schram, 1970. FMNH PE 16527, holotype, lateral view (illustrated in Wilson, 2012 and Fig. 14j).

### 52.2. *Phylogenetic justification*

*H. shermani* was included in a morphological cladistic analysis by Wilson (2012). It occurred at the base of Phreatoicidea (Wilson, 2012), a position within the crown group of Isopoda and thus, crown Peracarida. Note that the assignment of our calibration fossil to crown Isopoda means that exclusion of Mysida from Peracarida (Jenner et al., 2009; Meland and Willassen, 2007; Spears et al., 2005) will not change the date assessed for Peracarida.

### 52.3. *Age justification*

As for 31.3.

### 52.4. *Discussion*

Pygocephalomorpha is a eumalacostracan fossil clade with a number of symplesiomorphic characters. Pygocephalomorpha were proposed as members of Peracarida; however, a recent phylogeny depicted them in a polytomy in any of three positions: sister to Mysidacea (Mysida + Lophogastrida), sister to all non-Mysidacea peracarids, or on the peracarid stem lineage (Jones et al., 2016). Note also the molecular analyses discussed above that exclude Mysida from Peracarida (e.g. Jenner et al., 2009), which would mean two of three equally parsimonious positions for Pygocephalomorpha may be outside the peracarid crown group. If Pygocephalomorpha were shown to be more likely within the crown group of Peracarida (e.g. as sister to non-mysid peracarids), one species in this clade, *Tealliocaris walloniensis* Gueriau et al., 2014, would become the oldest peracarid (from the Fammenian stage of the Devonian ~50 Myr older; Gueriau et al., 2014).

Within Peracarida, several orders have putative Mesozoic calibration fossils that do not fully meet our requirements. Putative Lophogastrida fossils were described from the middle Triassic Falang Formation of China (Taylor et al., 2001) and Gres a Voltzia, France (Bill, 1914). These were attributed to the extant family Eucopiidae (Taylor et al., 2001), although they resolved outside crown Lophogastrida in an older phylogeny including only the French species (Taylor et al., 1998). Putative Mysida were described from the Late Jurassic Solnhofen Plattenkalk of Germany, but are poorly preserved (Schram, 1986). Fossils separately attributed to each of Lophogastrida and Mysida have also been described from the Middle Jurassic La Voulte-sur-Rhone of France (Secrétan and Riou, 1986). Other fossils noted as “mysidaceans” have been mentioned, but not yet described, from the middle Triassic Luoping Biota of China (Feldmann et al., 2015;. None of these fossils have been evaluated in the context of molecular discoveries, i.e. they assume a sister relationship between Mysida and Lophogastrida. Therefore, none can be used as calibrations until their relationships with respect to potential polyphyletic Mysidacea have been assessed.

Fossils allied to Spelaeogriphacea have been described from the Cretaceous Yixian Formation of China section 60.3 for revised stratigraphy) and Las Hoyas of Spain (Jaume et al., 2013). However, poor preservation of diagnostic characters indicates these are stem spelaeogriphaceans (Jaume et al., 2013).

Although several Mesozoic fossils have been assigned to Amphipoda, none have sufficient or accurate morphological placement (Starr et al., 2016; Vonk and Schram, 2007). Thus no known Mesozoic fossil qualifies as a crown group amphipod.

## 53. Crown Isopoda

This clade comprises Asellota, Phoratopodidea, Cymothoida, Microcereberidea, Limnoridea, Sphaeromatidea, Valvifera, Oniscidea (wood lice/pill bugs), Phreatoicidea, Calabozoidea and Tainisopidea, their last common ancestor and all of its descendants (Fig. 13). The list of isopod suborders is derived from Boyko et al. (2008). Isopod monophyly was recovered in phylogenetic analysis of four housekeeping genes (Lins et al., 2012) and of combined ribosomal genes and morphology (Wilson, 2009).

### 53.1. *Fossil specimens*

As for 52.1.

### 53.2. *Phylogenetic justification*

As for 52.2.

### 53.3. *Age justification*

As for 31.3.

## 54. Crown Tanaidacea

This clade comprises Apseudomorpha, Neotanaidomorpha and Tanaidomorpha, their last common ancestor and all of its descendants (Fig. 13). Phylogenetic analysis of three molecular loci demonstrated monophyly of Apseudomorpha and Tanaidomorpha; Neotanaidomorpha could not be included (Drumm, 2010). More expansive outgroup sampling (without Neotanaidomorpha) did not recover monophyletic Tanaidacea with ribosomal and morphological data (Wilson, 2009). However, a nuclear ribosomal analysis including Neotanaidomorpha supported monophyly (Kakui et al., 2011).

### 54.1. *Fossil specimens*

*Alavatanais carabe* Vonk and Schram, 2007. MCNA 9537, holotype in amber, male (Fig. 14l). For additional morphological details, we also refer to MCNA 9846a and MCNA 13888 (Fig. 14m), both males.

### 54.2. *Phylogenetic justification*

Spanish amber tanaids were originally misidentified as amphipods (Alonso et al., 2000). The fossils were placed in a new family, Alavatanaidae, part of the superfamily Paratanaoidea within Tanaidomorpha (Sanchez-Garria et al., 2015; Vonk and Schram, 2007). Characters supporting affinity within Tanaidomorpha include the presence of an articulated ischium, articulation of the last two pleopods (may be reduced in males), and seven or fewer antennal articles (Sanchez-Garria et al., 2015).

### 54.3. *Age justification*

Amber inclusions bearing arthropod fossils were discovered from the Peñacerrada I outcrop, Basque-Cantabrian Basin, Alava, Spain (Alonso et al., 2000; Penalver and Delclos, 2010). The Peñacerrada I outcrop itself is divided into three intervals, with the lowest bearing the amber (Barron et al., 2015). Earlier palynological study assigned Peñacerrada I to the Escucha Formation, in the late Aptian (Barron et al., 2001). Recent restudy, however, amended this outcrop to the Utrillas Group (Barron et al., 2015). The presence of marine palynomorphs characterized by *Chichaouadinium vestitum* and *Palaeohystrichophora infusorioides,* and the terrestrial *Distaltriangulisporites mutabilis* and *Senectotetradites varireticulatus* together constrain a late Albian age for the Penacerrada I (Barron et al., 2015). The upper boundary of the Albian stage is 100.5 Ma ± 0.4 Myr (Ogg et al., 2012), providing a minimum estimate for Alava amber fossils at 100.1 Ma.

Soft maximum as for 26.3.

## 55. Crown Eucarida

This clade comprises Euphausiacea (krill) and Decapoda (crabs, shrimp, lobsters), their last common ancestor and all of its descendants (Fig. 13). Monophyly of Euphausiacea and Decapoda is supported by phylogenetic analysis of transcriptomes (von Reumont et al., 2012), ribosomal genes (Spears et al., 2005), and combined molecular and morphological data (Jenner et al., 2009; Wills et al., 2009). Amphionidacea was represented as its own order within Eucarida based on morphology (Jenner et al., 2009; Wills et al., 2009), but recently the first molecular sequence data have become available, indicating that Amphionidacea are larval stages of Caridea (i.e. within Decapoda) (De Grave et al., 2015).

### 55.1. *Fossil specimens*

As for 49.1.

### 55.2. *Phylogenetic justification*

As for 49.2.

### 55.3. *Age justification*

As for 49.3.

## 56. Crown Decapoda

This clade comprises Dendrobranchiata (shrimp/prawns) and Pleocyemata (caridean shrimp, mud shrimp, true crabs, hermit and king crabs, lobsters, spiny lobsters, etc.), their last common ancestor and all of its descendants (Fig. 13). Decapod monophyly is established by phylogenetic analysis of protein-coding genes (Bracken et al., 2009; Bybee et al., 2011; Timm and Bracken-Grissom, 2015), morphology (Legg et al., 2013; Richter and Scholtz, 2001), and combined morphology and molecular data (Jenner et al., 2009). Analyses of whole mitochondrial genomes place Euphausiacea (krill) within Decapoda (Shen et al., 2015), a result congruent with acquisition of a nauplius larval stage (though this is accepted as convergent: (Jirikowski et al., 2013; Scholtz, 2000). We apply caution in interpreting deep splits inferred from mitochondrial DNA (Simon and Hadrys, 2013); hence, we accept decapod monophyly to the exclusion of krill.

### 56.1. *Fossil specimens*

As for 49.1.

### 56.2. *Phylogenetic justification*

As for 49.2.

### 56.3. *Age justification*

As for 49.3.

## 57. Crown Allotriocarida

This clade comprises Branchiopoda, Cephalocarida, Remipedia, and Hexapoda, their last common ancestor and all of its descendants (Fig. 15). Monophyly of this clade was proposed by a combined phylogenetic analysis of transcriptomes, nuclear protein-coding genes, and morphology (Oakley et al., 2013). As of this writing, a transcriptome remains to be sequenced for Cephalocarida. This clade is not recognized in GenBank taxonomy.

**Fig. 15.**
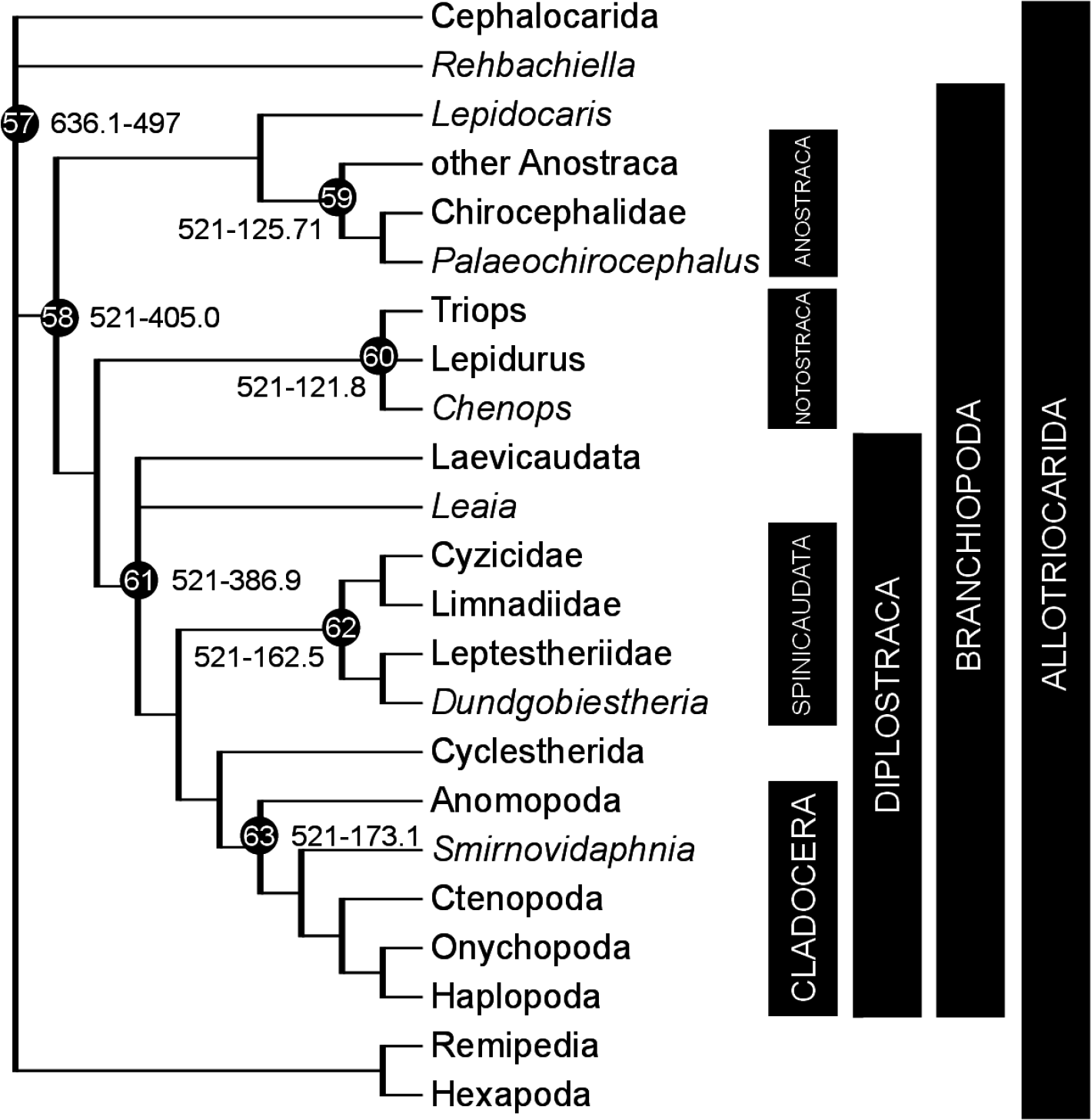
Calibration diagram for Allotriocarida (nodes 57–63).

### 57.1. *Fossil specimens*

*Rehbachiella kinnekullensis* Müller, 1983. UB 644, holotype, consisting of a complete larva (Fig. 16a). This species has been extensively documented by Walossek (1993).

**Fig. 16.**
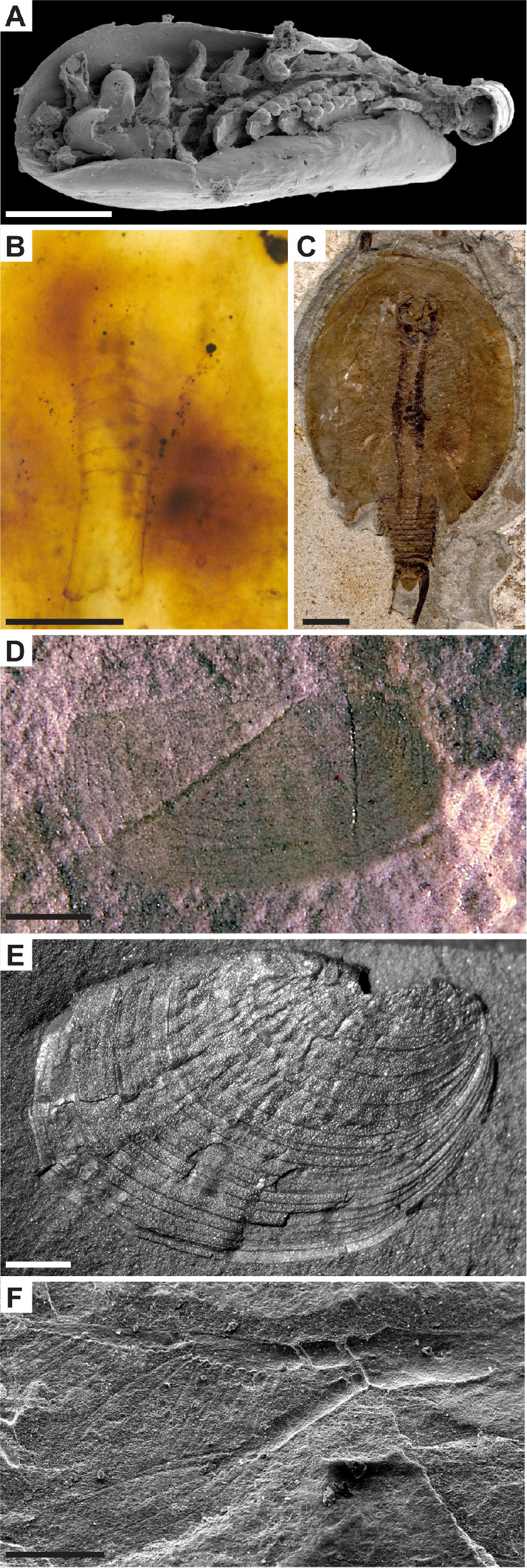
Allotriocarida fossil calibrations for (A) node 57: *Rehbachiella kinnekuUensis,* UB 611, scale bar 200pm, image credit D. Waloszek; (B) node 58: *Lepidocaris rhyniensis,* NHMUK In. 24493, scale bar 200pm, image credit NHMUK; (C) node 60: *Chenops yixianensis,* CNU-CAL-HP-2009001, scale bar 10mm, image credit T. Hegna; (D) node 61: *Leaia chinensis,* NIGP 51786, scale bar 1mm, image credit Y. Shen; (E) node 62: *Dundgobiestheria mandalgobiensis,* ED-A-14–1, scale bar 1 mm, image credit G. Li; (F) node 63: *Smirnovidaphnia smirnovi,* PIN 1873/100, scale bar 200 pm, image credit A. Kotov.

### 57.2. *Phylogenetic justification*

Recent phylogenetic analyses including *R. kinnekullensis* have strongly indicated a position on the branchiopod stem lineage (morphology: Legg et al., 2013; combined morphology and molecular: Oakley et al., 2013) or the cephalocarid stem lineage (morphology: Wolfe and Hegna, 2014). Either relationship (or a strict consensus position on the stem of Branchiopoda + Cephalocarida) would still be within the crown group of Allotriocarida.

### 57.3. *Age justification*

As for 35.3.

## 58. Crown Branchiopoda

This clade comprises Anostraca (fairy/brine shrimp), Notostraca (tadpole shrimp) and Diplostraca, their last common ancestor and all of its descendants (Fig. 15). Monophyly of this clade is established by phylogenetic analysis of protein-coding genes (Regier et al., 2010; Spears and Abele, 2000), transcriptomes (von Reumont et al., 2012), morphology (Legg et al., 2013), and combined molecular and morphological data (Oakley et al., 2013).

### 58.1. *Fossil specimens*

*Lepidocaris rhyniensis* Scourfield, 1926. BMNH IN. 24493, holotype (Fig. 16b).

### 58.2. *Phylogenetic justification*

*L. rhyniensis* has been included in several phylogenetic analyses, coded from a composite of holotype and paratype material (Scourfield, 1926). With morphology only, *L. rhyniensis* is sister to extant Anostraca (Hegna, 2012; Legg et al., 2013); with morphology and molecular data, it is within Anostraca (Oakley et al., 2013). Therefore, it is unequivocally a crown group member of Branchiopoda.

### 58.3. *Age justification*

Minimum as for 10.3. Soft maximum as for 26.3.

### 58.4. *Discussion*

*R. kinnekullensis* has been frequently used to polarize the evolution of Branchiopoda. Its fossils are known only from larval stages, which may confound discussions of its morphology (Wolfe and Hegna, 2014). Recent phylogenetic analyses have indicated a position on the branchiopod stem lineage (Legg et al., 2013; Oakley et al., 2013) or the cephalocarid stem lineage (Wolfe and Hegna, 2014), excluded from the branchiopod crown. Nevertheless, divergence time analyses suggest Cambrian origins for crown Branchiopoda (Oakley et al., 2013), such that molecular clock estimates do not conflict with branchiopod affinities for Cambrian filter plate fossils (Harvey and Butterfield, 2008) or *R. kinnekullensis*.

## 59. Crown Anostraca

This clade comprises Artemiidae, Branchinectidae, Branchipodidae, Chirocephalidae, Parartemiidae, Streptocephalidae, Tanymastigidae and Thamnocephalidae, their last common ancestor and all of its descendants (Fig. 15). Monophyly of five sampled families is established by phylogenetic analysis of six housekeeping genes and morphology (Richter et al., 2007). Full sampling of families produced monophyletic Anostraca in an analysis of one ribosomal gene (Weekers et al., 2002).

### 59.1. *Fossil specimens*

*Palaeochirocephalus rasnitsyni* Trussova, 1975. TsGM 7a/10303 and 9/10303, preserving male antennae, and TsGM 2/10303, preserving a female body. The holotype does not preserve any diagnostic characters for the Anostraca crown group.

### 59.2. *Phylogenetic justification*

*P. rasnitsyni* (formerly *Chirocephalus rasnitsyni* Trussova, 1971) has not been included in a phylogenetic analysis. Taxonomic placement of its family, Palaeochirocephalidae, implicitly relates them to the extant family Chirocephalidae, though this family is considered *incertae sedis* by Rogers (2013). Morphological characters (shared with Chirocephalidae) supporting this relationship include 11 thoracic appendages bearing two pre-epipodites, the nine-segmented abdomen, and the basally separated two-segmented antennae in males (Trussova, 1971). A possible position on the stem lineage of Chirocephalidae would therefore place *P. rasnitsyni* within the crown group of Anostraca.

### 59.3. *Age justification*

The type locality of *P. rasnitsyni,* briefly described by Trussova (1971), is the left bank of Daya River, upstream from Shiviya Falls, in eastern Transbaikal, Russia. This locality, within the Unda-Daya Basin, has been assigned to the Glushkovo Formation (Sinitshenkova, 2005). The age of the Glushkovo Formation is poorly constrained, suggested as Late Jurassic (Sinitsa and Starukhina, 1986), Early Cretaceous (Sinitshenkova, 2005; Zherikhin et al., 1998), or perhaps at the Jurassic/Cretaceous boundary (Rasnitsyn and Quicke, 2002). However, *P. rasnitsyni* itself (along with palaeopteran insects such as *Proameletus caudatus* and *Equisetum undense)* correlates the Glushkovo Formation to the Baigul locality, also in Transbaikalia (Ignatov et al., 2011). The Baigul locality preserves fossil *Bryokhutuliinia jurassica,* one of only five known genera of Jurassic mosses (Ignatov et al., 2011).

Thus Baigul can be correlated to the Ulugey Formation of Mongolia, which also preserves *Bryokhutuliinia* fossils (Ignatov, 1992). The Ulugey Formation, in turn, is correlated to the La Cabrúa (Sierra del Montsec, Pyrenees, Spain) locality based on the shared presence of the coleopteran genus *Gobicar* (Gratshev and Zherikhin, 2000; Legalov, 2010; Soriano et al., 2006). Fossil charophyte algae *(Atopochara trivolvis triquetra)* indicate an age of late Hauterivian-early Barremian for the freshwater deposits of La Cabrúa (Gomez et al., 2002; Martm-Closas and Lopez-Moron, 1995). Although it has been proposed that a minimum age of the Montsec limestone may be as young as the end Maastrichtian (O’Reilly et al., 2015), recent biostratigraphic work proposes the last appearance of *A trivolvis triquetra* is correlated to the *Deshayesites weissi* ammonite Zone at its youngest (Martín-Closas et al., 2009). Revision of Tethyan ammonite dates indicates the *D. weissi* Zone, now the *D. forbesi* Zone, had an upper boundary of 125.71 Ma (Ogg et al., 2012a). This age is early Aptian, and provides a minimum for the correlated Glushkovo Formation.

Soft maximum as for 26.3.

### 59.4. *Discussion*

Oakley et al. (2013) placed the Early Devonian *L. rhyniensis* in the crown group of Anostraca, having sampled extant members of Artemiidae and Streptocephalidae. Earlier descriptions (Sanders, 1963; Scourfield, 1940a, 1926; Walossek, 1993) support this position. However, the most extensively sampled morphological analyses of Hegna (2012) consistently place *L. rhyniensis* as sister to all extant Anostraca. The Late Devonian *Haltinnaias serrata* Gueriau et al., 2016, described from both sexes, likely also belongs to the total group of Anostraca.

Other fossils from the Late Jurassic Daohugou Beds of China (Huang et al., 2006; Shen and Huang, 2008) and Early Cretaceous Koonwarra Formation of Australia are likely to belong to the crown group of Anostraca, as they have been included in a morphological phylogenetic analysis (Hegna, 2012), but these have not been described in detail.

## 60. Crown Notostraca

This clade comprises two extant genera, *Triops* and *Lepidurus*, their last common ancestor and all of its descendants (Fig. 15). Monophyly is established by phylogenetic analysis of housekeeping genes (Mathers et al., 2013), morphology (Lagebro et al., 2015), and combined morphological and molecular data (Richter et al., 2007).

### 60.1. *Fossil specimens*

*Chenops yixianensis* Hegna and Ren, 2010. CNU-CAL-HP-2009001 (part; Fig. 16c) and CNU-CAL-HP-2009002 (counterpart), holotype.

### 60.2. *Phylogenetic justification*

In the morphological phylogenetic analysis of Lagebro et al. (2015), *C. yixianensis* was in a polytomy with other crown Notostraca, including *Lepidurus batesoni*. The equal size of thoracic endopods and fourth and fifth endites may exclude *C. yixianensis* from crown Notostraca (Hegna and Ren, 2010). However, in previous morphological phylogenies (Hegna, 2012), *C. yixianensis* was sister taxon to the extant *L. batesoni*. This is because *L. batesoni* lacks elongated endites on the first thoracic appendage, suggesting a synapomorphy between *C. yixianensis* and the extant species, and perhaps membership in an entirely different genus (Hegna, 2012; Hegna and Ren, 2010). If indeed *C. yixianensis* is, based on phylogenies and endite morphology, a sister group of *L. batesoni,* it would remain within the crown Notostraca.

### 60.3. *Age justification*

This fossil was discovered in the Yixian Formation of northeastern China (Hegna and Ren, 2010). The Yixian Formation lies between the overlying Jiufotang Formation and underlying Tuchengzi Formation; together they comprise the Jehol Group (e.g. Chang et al., 2009a; Zhou, 2006). Dating of the Jehol Group has been contentious, varying from Late Jurassic to Early Cretaceous based on biostratigraphic and radiometric techniques. Recent 40Ar/39Ar dates yielded ages of 129.7 Ma ± 0.5 Myr for basaltic lava from the bottom of the Yixian Formation and 122.1 Ma ± 0.3 Myr for tuff layers at the bottom of the overlying Jiufotang Formation (Chang et al., 2009a). Other age estimates have fallen within this range (reviewed by Zhou, 2006). This debate underscores the point that reasonably precise radiometric dates may still be quite inaccurate. We conservatively use the younger of these estimates, so a minimum age for Yixian fossils is 121.8 Ma.

Soft maximum as for 26.3.

### 60.4. *Discussion*

*Strudops goldenbergi* Lagebro et al., 2015 was recently described from the Late Devonian, which would be the oldest notostracan. In a morphological phylogenetic analysis, however, it could only be identified as a member of the total group (Lagebro et al., 2015), and thus cannot assign a minimum age to crown Notostraca.

Morphological conservatism (i.e. Permian and Triassic fossils erroneously attributed to the extant species *Triops cancriformis)* has led to the misleading name ‘living fossil’ for Notostraca (Hegna, 2012; Mathers et al., 2013). Once touted as the ‘oldest living species,’ *T. cancriformis permiensis,* from the Permian of France, is more similar to the co-occurring *Lepidurus occitaniacus* than extant *Triops* (Lagebro et al., 2015). Decay experiments on living *T. cancriformis* confirm that carapace characters and elongated endites of the first thoracic limb are phylogenetically meaningful, thus rejecting a referral of any fossil to the extant species (Hegna, 2012).

## 61. Crown Diplostraca

This clade comprises Laevicaudata, Spinicaudata, Cyclestherida (these three collectively: ‘clam shrimp—) and Cladocera (water fleas), their last common ancestor and all of its descendants (Fig. 15). Monophyly is supported by phylogenetic analysis of transcriptomes (von Reumont et al., 2012) and 62 nuclear protein-coding genes (Regier et al., 2010), and combined molecules and morphology (Oakley et al., 2013) although Cyclestherida was not sampled in these analyses. Smaller molecular analyses (Richter et al., 2007 with indel costs) and morphological analyses (Hegna, 2012; Legg et al., 2013; Olesen, 2009, 1998; Richter et al., 2007) including data for Cyclestherida also support monophyly of Diplostraca.

### 61.1. *Fossil specimens*

*Leaia chinensis* Shen, 1983. NIGP 51786, holotype, preserving a left valve (Fig. 16d).

### 61.2. *Phylogenetic justification*

*L. chinensis* has not been treated in a phylogenetic context. It is placed within the fossil family Leaiidae primarily based on carapace shape, including the nearly straight ventral edge (Shen, 1978; Tasch, 1987). However, congeneric fossils, *L. gondwanella* Tasch, 1987 and *L. canadensis* Shen and Schram, 2014, may have preserved soft parts (Shen and Schram, 2014). These include biramous antennae, mandibles, as well as putative shell glands and digestive tubes, and radiating carinae (Shen and Schram 2014). The short and delicate antennal flagella and radiating carinae of *L. chinensis,* in particular, are similar to extant Spinicaudata and Cyclestherida; the presence of growth lines is only known from Spinicaudata (Shen and Schram 2014). However, the head shape of *L. chinensis* is more similar to Laevicaudata. This suggests phylogenetic positions either on the stem of Onychocaudata (Spinicaudata, Cyclestherida, and Cladocera) or on the stem of Laevicaudata. A position as stem Diplostraca would mean growth lines are ancestral for all diplostracans, and have been lost twice (in the ancestors of Laevicaudata and Cyclestherida + Cladocera), which is unparsimonious. Hence we support *L. chinensis* within crown Diplostraca.

### 61.3. *Age justification*

*L. chinensis* was found in sediments of the Guitou Group, near Chengma village, Hepu county, Guangxi Zhuang Autonomous Region, southern China (Shen, 1978). The upper subgroup of the Guitou Group, bearing conchostracan fossils, has purple-grey sandy shales within mudstone lamination, and is overlain by the Tungkangling Formation (Shen, 1978). The brachiopod *Stringocephalus,* the stromatoporoid corals *Endophyllum* and *Sunophyllum,* and the ostracods *Samarella crassa* and *Tuberokloedenia bituberculata* together indicate a Givetian age for the upper Tungkangling Formation (Liao and Ruan, 2003). The underlying Yingtang Formation (as well as the lower Tungkangling Formation) are correlated to the Eifelian Stage by the ostracods *Bairdocypris biesenbachi* and *Flatiella subtrapezoidalis* (Liao and Ruan, 2003), the brachiopod *Bornhardtina* and the conodonts *Polygnathus australis, P. costatus,* and *P. partitus* (Ma et al., 2009). As the conchostracan-bearing sediments underlie the Tungkangling Formation, they are no younger than Eifelian in age. The upper boundary of the Eifelian is 387.7 Ma ± 0.8 Myr, providing a minimum age of 386.9 Ma.

Soft maximum as for 26.3.

## 62. Crown Spinicaudata

This clade comprises Leptestheriidae, Cyzicidae and Limnadiidae, their last common ancestor and all of its descendants (Fig. 15). Clade names are defined in Schwentner et al. (2009). Monophyly is established by phylogenetic analysis of morphology (Hegna, 2012; Olesen, 1998; Richter et al., 2007), three housekeeping genes (Schwentner et al., 2009), and six housekeeping genes plus morphology (Richter et al., 2007).

### 62.1. *Fossil specimens*

*Dundgobiestheria mandalgobiensis* Li et al., 2014. ED-A-14–1, holotype (Fig. 16e).

### 62.2. *Phylogenetic justification*

Phylogenetic analysis of spinicaudatan fossils is rare, owing to difficulty in character identification (Astrop and Hegna, 2015; see also 62.3). Members of Leptestheriidae appear to share emergence of dendritic reticulation and anastomizing ridge ornamentation with the fossil spinicaudatan family Loxomegaglyptidae (Astrop and Hegna, 2015; Shen, 1994). *D. mandalgobiensis* is perhaps the oldest definitive Loxomegaglyptidae, based on large-sized reticulate ornamentation and weakly defined growth lines, shared with other members of the family (Li et al., 2014). Due to the above ornamentation characters (Astrop and Hegna, 2015; T. Astrop, pers. comm.), it can be placed on the stem lineage of Leptestheriidae, and thus within crown Spinicaudata.

### 62.3. *Age justification*

*D. mandalgobiensis* is from the Eedemt Formation, Khootiin Khotgor coal mine region, Dundgobi Province, southeast Mongolia (Li et al., 2014). The spinicaudatan genus *Triglypta* (cooccurring with *D. mandalgobiensis*) provides a biostratigraphic constraint on the Eedemt Formation, as *Triglypta* species also occur in both the (older) *Euestheria ziliujingensis* fauna and (younger) *Sinokontikia* fauna in lacustrine sequences of northwestern China (Li et al., 2014; Li and Matsuoka, 2012). First, the *E. ziliujingensis* fauna is distributed throughout east Asia, dated to a Bajocian-Bathonian (Jurassic) age by the occurrence of *Euestheria trotternishensis* (Chen et al., 2007). *E. trotternishensis* co-occurs in the *Skyestheria* spinicaudatan fauna of Skye, Scotland; the Bajocian-Bathonian date for this locality comes from ammonite and palynological index fossils (Chen and Hudson, 1991). Although the *Sinokontikia* fauna was once thought stratigraphically equal to the late *E. ziliujingensis* fauna (Chen et al., 2007; Li and Matsuoka, 2012), *Sinokontikia* has been determined to be younger based on occurrence in the Qiketai Formation of the Turpan Basin, northwest China. The Qiketai Formation is constrained only to the Callovian. As Chinese *Triglypta* (the index genus for the Eedemt Formation) is absent from any higher strata, a minimum age comes from the upper boundary of the *Sinokontikia* fauna (Li et al., 2014). The upper boundary of the Callovian is 163.5 Ma ± 1.0 Myr, giving a minimum age for the Eedemt Formation of 162.5 Ma.

Soft maximum as for 26.3.

### 62.4. *Discussion*

Preservation of the calcified spinicaudatan carapace is extremely common (some fossil species are used as biostratigraphic indices), but characters diagnostic for extant taxa (i.e. soft parts) are rarely preserved (Hegna, 2012; Orr and Briggs, 1999). Uniquely among ‘conchostracans’; (and indeed most arthropods), Spinicaudata do not moult their carapace, instead preserving growth lines. The number of growth lines necessarily increases through ontogeny, so it is a poor character to demonstrate phylogenetic relationships. Therefore relationships among fossil taxa have been determined based on ornamentation of the carapace (e.g. Gallego, 2010); however, these characters have not yet been integrated with morphological study of extant carapaces. Ongoing work seeks to demonstrate the utility of carapace ornamentation as valid phylogenetic characters (Astrop and Hegna, 2015). Furthermore, integration of past descriptive work is hindered by language barriers between different workers (Chinese, Russian, and English; summarized by Astrop and Hegna, 2015).

For example, some poorly known fossils from the Middle Devonian of Antarctica, described as members of the genus *Cyzicus (Euestheria),* may be assigned to Spinicaudata (Tasch, 1987), possibly on the stem lineage of ‘non-Australian Cyzicidae’ *sensu* Schwentner et al. (2009). Characters linking the Antarctic fossils to the living clade, however, are dubious. There are reports of uncertain Late Devonian stem group members for Limnadiidae (Liu and Gao, 1985), and much more likely Permian stem Limnadiidae (Novojilov, 1970), but these fossils are insufficiently described, leaving any specific crown spinicaudatan character states debatable. Therefore we must caution that Jurassic age is likely a severe underestimate of crown spinicaudatan antiquity. Finally, the Sinemurian *Afrolimnadia sibiriensis* Tasch, 1987 was used to calibrate crown Spinicaudata in a divergence time analysis, but with limited justification of characters for Limnadiidae (Bellec and Rabet, 2016).

## 63. Crown Cladocera

This clade comprises Anomopoda, Ctenopoda, Onychopoda and Haplopoda, their last common ancestor and all of its descendants (Fig. 15). Monophyly of this clade is supported by phylogenetic analysis of housekeeping genes alone (Stenderup et al., 2006), morphology (Hegna, 2012; Olesen, 2009, 2007, 1998), and six housekeeping genes plus morphology (Richter et al., 2007).

### 63.1. *Fossil specimens*

*Smirnovidaphnia smirnovi* Kotov, 2007, illustrated by two specimens: PIN 1873/100 (holotype, preserving the second antenna; Fig. 16f) and PIN 1873/105 (paratype, preserving both second antennae and mandibles).

### 63.2. *Phylogenetic justification*

The setal armature of the second antenna is similar to that found in extant Ctenopoda (Kotov, 2007). In morphological phylogenetic analyses, *S. smirnovi* was either found in a basal polytomy with all of crown Cladocera, or it was sister to all Cladocera except Anomopoda (Hegna, 2012). The polytomy could indicate a stem cladoceran position, but the analyses in which *S. smirnovi* was within crown Cladocera included a lesser amount of missing data, and thus may be more robust. Furthermore, Ctenopoda itself was not recovered as monophyletic in these analyses, but *S. smirnovi* was related to taxa that have been previously included in Ctenopoda (Hegna, 2012).

### 63.3. *Age justification*

*S. smirnovi* was found on the right bank of the Angara River, downstream of Ust’-Baley village in the Olonkovsky District of Asian Russia (Kotov, 2007). According to the presence of *Osmundacidites-type* plant spores, the Ust’-Baley outcrop of the Cheremkhovo or Cheremkhovskaya Formation is correlated to the late Toarcian stage of the Early Jurassic (Akulov et al., 2015). The younger boundary of the Toarcian is 174.1 Ma ± 1.0 Myr, therefore giving a minimum age of 173.1 Ma.

Soft maximum as for 26.3.

### 63.4. *Discussion*

*Ebullitiocaris oviformis* Anderson et al., 2003 from the Devonian Rhynie Chert and *E. elatus* Womack et al., 2012 from Carboniferous chert have both been described as Cladocera. Fragmentation patterns of both fossils are inconsistent with those observed from decay experiments, therefore casting doubt on their cladoceran membership (Hegna, 2012).

Other Jurassic/Cretaceous members of the fossil family Prochydoridae are known from Mongolia and Asian Russia (Kotov, 2009). However, the Prochydoridae have been proposed as a member of the stem lineage of Anomopoda, the stem of all non-Anomopoda Cladocera, as well as the stem of Cladocera itself (Kotov, 2013, Kotov, 2009). Thus crown-group affinity cannot be confirmed.

## 64. Crown Hexapoda

This clade comprises Collembola (springtails), Protura (cone heads), Diplura (two-pronged bristletails) and Insecta (insects), their last common ancestor and all of its descendants (Fig. 17). Monophyly of Hexapoda is established by phylogenetic analysis of nuclear protein-coding genes (Regier et al., 2010, 2005; Sasaki et al., 2013), transcriptomes (Dell’Ampio et al., 2014; Misof et al., 2014), and morphology (Legg et al., 2013).

**Fig. 17.**
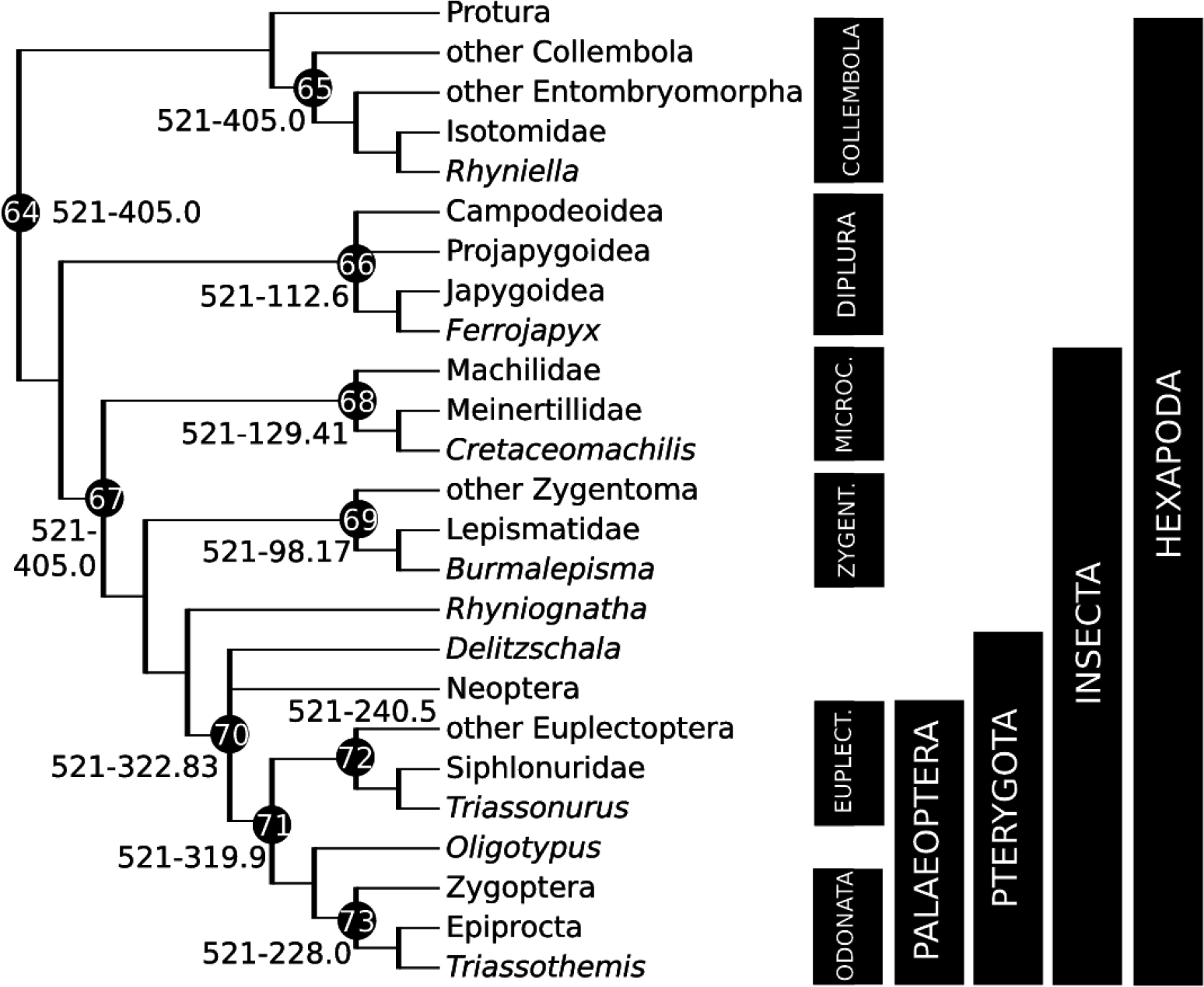
Calibration diagram for non-neopteran Hexapoda (nodes 64–73). Microc. = Microcoryphia, Zygent. = Zygentoma, Euplect. = Euplectoptera.

### 64.1. *Fossil specimens*

*Rhyniella praecursor* Hirst and Maulik, 1926. NHMUK IN. 27765, lectotype *(fide* Ross and York, 2003), head capsule with mouthparts in chert (Fig. 2f). Paralectyotypes NHMUK IN. 3822538227.

### 64.2. *Phylogenetic justification*

The Rhynie Chert taxon *R. praecursor* Hirst and Maulik, 1926, was originally identified as an insect. Re-identification as a poduromorph collembolan was made by Tillyard (1928). Additional material was examined by Scourfield (1940a, 1940b), who considered *R. praecursor* to be a possible entomobryomorph rather than a poduromorph. Subsequent studies, including some additional specimens, were made by Massoud (1967), Whalley and Jarzembowski (1981), Greenslade (1988), and Greenslade and Whalley (1986). The most recent of these investigations favour membership of at least some specimens assigned to *R. praecursor* in the extant entomobryomorph family Isotomidae (Greenslade and Whalley 1986). Greenslade (1988) indicated that three species could be represented in the sample that is currently assigned to *R. praecursor*. Identification as an entomobryomorph underpins an assignment to the crown groups of Collembola and Hexapoda. Other taxa from the Rhynie Chert *(Rhyniognatha* and *Leverhulmia;* Engel and Grimaldi, 2004; Fayers and Trewin, 2005) and thus as old as *R. praecursor* have also been identified as Hexapoda, and more precisely as Insecta.

### 64.3. *Age justification*

As for 58.3.

## 65. Crown Collembola

This clade comprises Entomobryomorpha, Neelipleona, Poduromorpha, Symphypleona and Tomoceroidea, their last common ancestor and all of its descendants (Fig. 17). Most studies with substantial outgroup sampling are based on only one or two subclades of Collembola; however, phylogenetic analysis of ribosomal genes including all subclades (Gao et al., 2008; Xiong et al., 2008) demonstrated monophyly. Additional support with limited sampling of Collembola subclades comes from transcriptomes (Dell’Ampio et al., 2014), mitochondrial genomes (Carapelli et al., 2007), and morphology (D’Haese, 2003).

### 65.1. *Fossil specimens*

As for 64.1.

### 65.2. *Phylogenetic justification*

As for 64.2.

### 65.3. *Age justification*

As for 58.3.

## 66. Crown Diplura

This clade comprises Projapygoidea, Japygoidea and Campodeoidea, their last common ancestor and all of its descendants (Fig. 17). Monophyly of Diplura has been demonstrated by phylogenetic analysis of ribosomal genes (Gao et al., 2008), whole mitochondrial genomes (Chen et al., 2014), and morphology (Koch, 1997).

### 66.1. *Fossil specimens*

*Ferrojapyx vivax* Wilson and Martill, 2001. SMNS 64276, holotype.

### 66.2. *Phylogenetic justification*

*F vivax* is “morphologically indistinguishable” (Wilson and Martill, 2001) from extant Japygoidea, sharing characters such as 40-segmented antennules, abdominal tergites 1–8 with a median suture, abdominal tergite 9 significantly shorter, conical abdominal styli, and forcipate cerci with curved inner margins and lacking obvious denticles (Wilson and Martill, 2001). Monophyly of Japygoidea has been considered “beyond doubt” (Koch, 2009; Fig. 3 therein for cladogram), so a phylogenetic position for this fossil either within or closely related to that clade would place it within crown Diplura.

### 66.3. *Age justification*

As for 29.3.

## 67. Crown Insecta

This clade comprises Microcoryphia (jumping bristletails), Zygentoma (silverfish) and Pterygota (winged insects), their last common ancestor and all of its descendants (Fig. 17). Monophyly is established by phylogenetic analysis of nuclear protein-coding genes (Regier et al., 2010, 2005; Sasaki et al., 2013), transcriptomes (Dell’Ampio et al., 2014; Misof et al., 2014), and morphology (Legg et al., 2013).

### 67.1. *Fossil specimens*

*Rhyniognatha hirsti* Tillyard, 1928. BMNH IN. 38234, holotype, preserving the mandibles and their articulation (Fig. 18d). Redescribed and imaged by Engel and Grimaldi (2004).

**Fig. 18.**
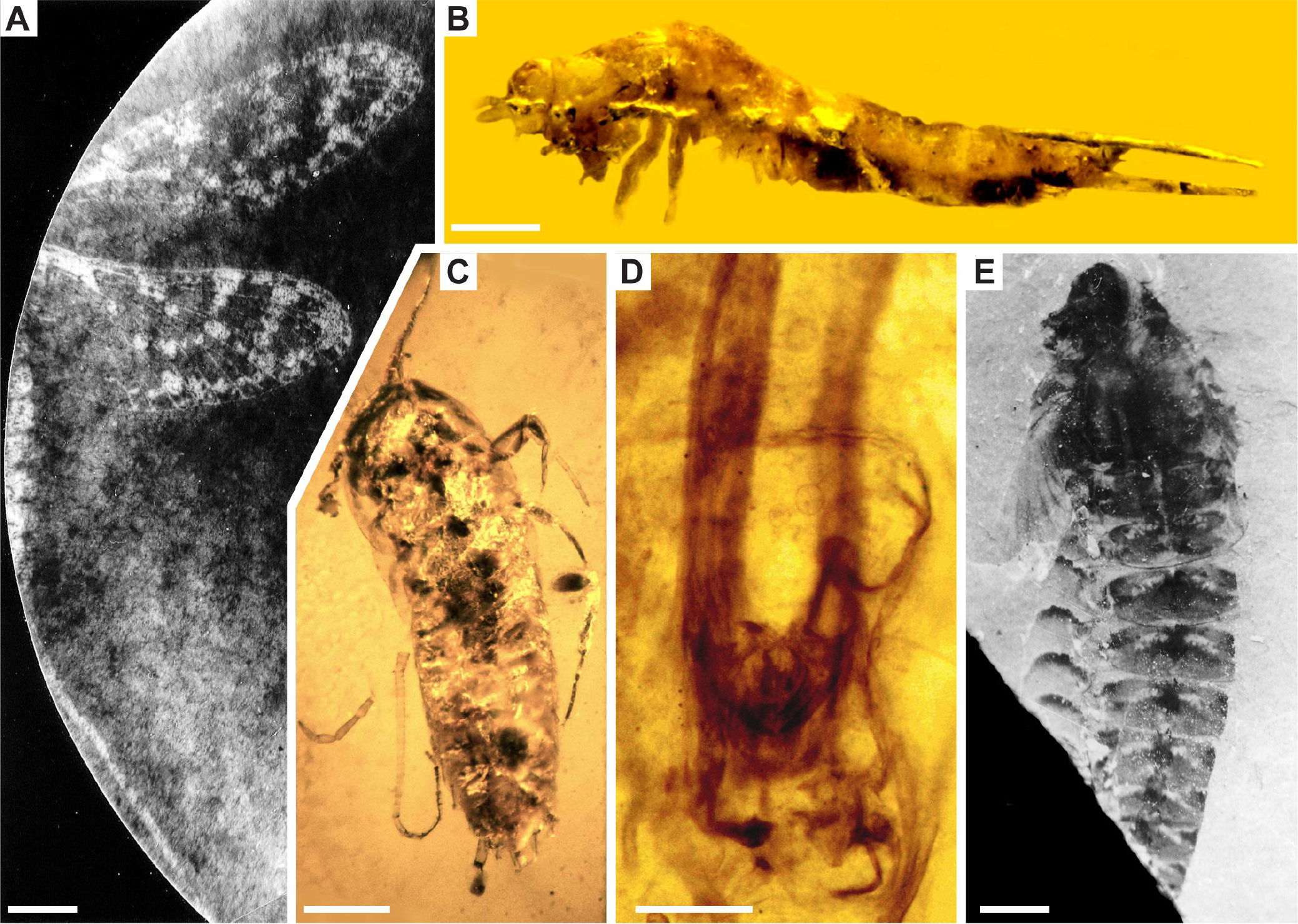
Hexapoda fossil calibrations for (A) node 70: *Delitzschala bitterfeldensis,* BGR X 9216, scale bar 2mm, image credit C. Brauckmann; (B) node 68: *Cretaceomachilis libanensis,* Milki No. 194/35, scale bar 500μm, image credit G. Poinar; (C) node 69: *Burmalepisma cretacicum,* B-TH-1, scale bar 500μm, image credit G. Poinar; (D) node 67: *Rhyniognatha hirsti,* BMNH IN. 38234, scale bar 200pm, image credit NHMUK; (E) node 72: *Triassonurus doliiformis,* Louis Grauvogel collection No. 9304, scale bar 10mm, image credit L. Grauvogel-Stamm.

### 67.2. *Phylogenetic justification*

The only known specimen (the holotype) of *R. hirsti* demonstrates that the preserved pair of mandibles articulate at two points, i.e. are dicondylic (Engel and Grimaldi, 2004). Dicondylic mandibles are a diagnostic synapomorphy of Insecta (including Microcoryphia; Blanke et al., 2015). Although *R. hirsti* has not been included in numerical phylogenetic analyses, its mandibular shape implies “metapterygote” affinities (Engel and Grimaldi, 2004) and accordingly crown group membership within Insecta.

### 67.3. *Age justification*

As for 58.3.

### 67.4. *Discussion*

A complete body fossil of a putative insect, *Strudiella devonica* Garrouste et al., 2012, was described from the Fammenian (372–359 Ma) of Strud, Belgium. Apomorphies supporting an insect affinity (such as the structure of the mandibles and the number of legs), however, are poorly preserved and potentially over-interpreted (Hornschemeyer et al., 2013).

## 68. Crown Microcoryphia

This clade comprises the families ‘Machilidae’ and Meinertillidae, their last common ancestor, and all of its descendants (Fig. 17). This clade is commonly referred to as Microcoryphia in taxonomic literature and Archaeognatha in phylogenetic literature (Gaju-Ricart et al., 2015). Recently it was clarified that Archaeognatha includes the completely extinct order Monura (compound eyes not fused, paracercus only) and Microcoryphia (very small head vertex due to their fused compound eyes, terminalia composed of a median paracercus and two lateral cerci), and thus the crown group refers to Microcoryphia only (Gaju-Ricart et al., 2015). Monophyly is supported by phylogenetic analysis of transcriptomes (Misof et al., 2014), nuclear protein-coding genes (Regier et al., 2010), and morphology (Legg et al., 2013). Synapomorphies are discussed and mapped on cladograms by Larink (1997) and Koch (2003).

### 68.1. *Fossil specimens*

*Cretaceomachilis libanensis* Sturm and Poinar, 1998 (collection Milki No. 194/35, deposited at the American University of Beirut, Beirut, Lebanon), holotype, male in amber (Fig. 18b).

### 68.2. *Phylogenetic justification*

C. *libanensis* shares morphological characters with the extant family Meinertillidae. These include the the absence of scales on the scapus, pedicellus and flagellum, and the presence of a characteristic hook near the distal end of article 2 of the male maxillary palp (Sturm and Poinar, 1998). The latter character is a synapomorphy of crown Meinertillidae (Sturm and Poinar, 1998), therefore the fossil is within crown Microcoryphia.

### 68.3. *Age justification*

*C. libanensis* was discovered in Cretaceous amber, from a locality in Lebanon that was not recorded (Sturm and Poinar, 1998). We therefore use a minimum age constraint from the youngest of the several known Lebanese amber localities, which all bear the same age within the early Barremian (Maksoud et al., 2016). The upper boundary of the early Barremian is proposed to be the first appearance of the ammonite *Ancyloceras vandenheckii* (Ogg et al., 2012). Cyclostratigraphy dates the *A vandenheckii* Zone beginning at 129.41 Ma (Ogg et al., 2012), providing a minimum age for Jezzine Lebanese amber fossils.

Soft maximum as for 26.3.

### 68.4. *Discussion*

Body imprint and trackway trace fossils from the Pennsylvanian have been attributed to both Archaeognatha and Zygentoma (Getty et al., 2013). Experiments with extant species of both clades indicate that archaeognathans produced fossil body imprints, as well as trackways exhibiting opposite symmetry (Getty et al., 2013). However, it is not possible to distinguish specific crown group apomorphies for these traces.

Dasyleptidae, a diverse fossil group known from the Carboniferous-Triassic, has been proposed as the sister group of Ectognatha (Collembola, Diplura, Protura) as well as part of Archaeognatha. Recent classifications place Dasyleptidae in Monura, a separate extinct suborder of Archaeognatha, and thus outside its crown group (Bechly and Stockar, 2011; Gaju-Ricart et al., 2015).

## 69. Crown Zygentoma

This clade comprises the families Lepismatidae, Nicoletiidae, Lepidotrichidae, Maindroniidae, and Protrinemuridae, their last common ancestor and all of its descendants (Fig. 17). Phylogenetic analyses of transcriptomes (Misof et al., 2014), protein-coding genes (Regier et al., 2010; Sasaki et al., 2013), morphology (Blanke et al., 2014), and combined molecular and morphological data (Kjer et al., 2006) with partial taxon sampling support monophyly.

### 69.1. *Fossil specimens*

*Burmalepisma cretacicum* Mendes and Poinar, 2008 (specimen B-TH-1 deposited in the Poinar amber collection maintained at Oregon State University; B-TH refers to Burma-Thysanura), holotype, a female in amber (Fig. 18c).

### 69.2. *Phylogenetic justification*

Although it has not been included in a phylogenetic analysis, *B. cretacicum* bears several morphological similarities to the extant family Lepismatidae. These include the presence of compound eyes, absence of ocelli, coxal, femur, and tarsal morphology, absence of a pronotal setal collar, and presence of only smooth macrochaetae (Mendes and Poinar, 2008). A phylogenetic position either on the stem or within the crown of Lepismatidae is within the crown group of Zygentoma.

### 69.3. *Age justification*

As for 33.3.

### 69.4 *Discussion*

A possible Zygentoma fossil has been recorded from the older Aptian Crato Formation of Brazil (Sturm, 1998), but is not sufficiently characterized to be a calibration fossil. Another fossil from Burmese amber, *Cretolepisma kachinicum* Mendes and Wunderlich, 2013, is also identified as Lepismatidae. This fossil is of equal age and affinity, and is therefore also an acceptable calibration for Zygentoma.

## 70. Crown Pterygota

This clade comprises Palaeoptera (mayflies, dragonflies) and Neoptera (flexible winged insects), their last common ancestor and all of its descendants (Fig. 17). Monophyly is established based on phylogenetic analysis of nuclear protein-coding genes (Regier et al., 2010, 2005; Sasaki et al., 2013), transcriptomes (Misof et al., 2014; Simon et al., 2012), and morphology (Legg et al., 2013).

### 70.1. *Fossil specimens*

*Delitzschala bitterfeldensis* Brauckmann and Schneider, 1996 (BGR X 9216), holotype, preserving a forewing, a hindwing, cerci, and part of the abdomen (Fig. 18a).

### 70.2. *Phylogenetic justification*

*D. bitterfeldensis* is considered a member of the family Spilapteridae, in the clade Palaeodictyopterida (Brauckmann and Schneider, 1996). Morphological characters supporting this relationship include the concave anterior wing margin and deeply bifurcate MA vein ending with two long branches (Brauckmann and Schneider, 1996; Li et al., 2014). Other fossils of Spilapteridae have also preserved the distinctive colour patterns and long cerci observed in *D. bitterfeldensis* (Li et al., 2014). Palaeodictyoptera have previously assumed to be related to extant Palaeoptera as they share the inability to fold their wings over the abdomen (a character observed in *D. bitterfeldensis*). However, a recent morphological phylogenetic analysis controversially recognized Palaeodictyopterida as the fossil sister group of Neoptera (Sroka et al., 2015). In this evolutionary scenario, palaeopterous wings are presumably a symplesiomorphy. Nonetheless, the position of Palaeodictyoptera suggested by Sroka et al. (2015) remains within crown Pterygota.

### 70.3. *Age justification*

*D. bitterfeldensis* was preserved in a core extracted from the locality Bore WISABAW 1315, in the Sandersdorf Formation of Bitterfeld, Germany (Brauckmann and Schneider, 1996). The core was correlated with the E2 ammonite zone, i.e. Arnsbergian (middle Namurian A) based on the cooccurrence of ostracods and conchostracans (Brauckmann et al., 1994). The Arnsbergian is further correlated to the late Serpukhovian stage at the end of the Mississippian (Pointon et al., 2012). The upper boundary of the Serpukhovian (base of the Pennsylvanian) is estimated at 323.23 Ma ± 0.4 Myr (Davydov et al., 2012), giving a minimum age of 322.83 Ma.

Soft maximum as for 26.3.

### 70.4. *Discussion*

An undescribed Namurian A (latest Mississippian) member of Archaeorthoptera was initially attributed to Orthoptera (Prokop et al., 2005). It was noted that the veins are strongly deformed, enough so that the authors were reluctant to make a formal systematic description (Prokop et al., 2005). Therefore, we regard the fossil as insufficiently characterized for dating. See section 67.4 for refutation of the even older *Strudiella devonica* (Garrouste et al., 2012).

## 71. Crown Palaeoptera

This clade comprises Euplectoptera (mayflies) and Odonata (dragonflies), their last common ancestor and all of its descendants (Fig. 17). Monophyly of this group has been challenged by morphology (Kristensen 1981) and some transcriptome data (Simon et al., 2012, 2009), though recent analyses of nuclear protein-coding genes (Regier et al., 2010) and transcriptomes have supported Palaeoptera (Misof et al., 2014; von Reumont et al., 2012), the former weakly, only in maximum likelihood analyses of the total dataset. Recent detailed morphological analyses of head morphology corroborate the monophyly of Palaeoptera (Blanke et al., 2013, 2012). Although a morphological phylogenetic analysis (Sroka et al., 2015) claims to reject palaeopteran monophyly, its constituent extant members, Euplectoptera and Odonata, are each others’; closest living relative and named therein as a new clade, Euhydropalaeoptera. We therefore cautiously endorse Palaeoptera as a clade of interest for dating.

### 71.1. *Fossil specimens*

*Oligotypus huangheensis* Li et al., 2013a (CNU-NX2006003), holotype, a forewing.

### 71.2. *Phylogenetic justification*

This species was originally described as *Sinomeganeura huangheensis* within the family Meganeuridae, part of Protodonata (Ren et al., 2008). Meganeurids include the “giant” dragonflies (with wings up to 710 mm), though *O. huangheensis* is much smaller at 70 mm (Ren et al., 2008). Despite the size difference, wing venation characters are consistent with classification in Protodonata, including the fusion of stems of CuP and CuA to a single oblique vein, distinctly stronger than the crossveins (Ren et al., 2008). This character, previously assumed to be synapomorphic for Meganeuridae, is more widespread within Protodonata (Li et al., 2013a). The group “Protodonata” itself is a paraphyletic stem group to crown Odonata, together within the total group Odonatoptera, defined by the simple MP vein (Sroka et al., 2015). Regardless of the precise relationship of Protodonata to crown Odonata, its members are definitively within crown Palaeoptera.

### 71.3. *Age justification*

The insect beds where this species was located are near Xiaheyan Village in the Qilianshan Mountains, Zhongwei County, Ningxia Huizu Autonomous Region, northwest China (Zhang et al., 2013). The insect fossil deposits are within the uppermost unit of the upper Tupo Formation (synonyms Hongtuwa or Zhongwei Formation). The presence of the ammonoids *Reticuloceras reticulatum, Gastrioceras listeri* and *G. montgomeryense* and conodonts *Declingnathodus noduliferous* and *Neognathodus symmetricus* indicate a Namurian B/C age (Xie et al., 2004; Yang, 1987; Yang et al., 1988; Zhang et al., 2013). The (upper) Namurian-(lower) Westphalian boundary is defined by the earliest occurrence of the goniatite *Gastrioceras subcrenatum* (Waters and Davies, 2006), but lacks a precise isotopic date. Pointon et al. (2012) estimated an age of c. 319.9 Ma for the base of the Westphalian (top of the Namurian, only slightly younger than the Marsdenian) based on Milankovitch cycles of sedimentation, giving a minimum age for Xiaheyan fossils.

Soft maximum as for 26.3.

### 71.4. *Discussion*

Due to the controversial status of Palaeoptera as a clade, there are several fossil groups that have been considered for membership: Palaeodictyopterida, Paoliida, stem mayflies (Ephemeroptera, see section 72), and stem dragonflies (Protodonata, including Geroptera) (Grimaldi and Engel, 2005). Each is discussed below.

Palaeodictyopterida, an abundant clade from the Carboniferous and Permian, have been assumed to be related to extant Palaeoptera as they share the inability to fold their wings over the abdomen. The oldest member, *Delitzschala bitterfeldensis,* predates all other Palaeoptera (and all other Pterygota with preserved wings), as it is from the Mississippian (Brauckmann et al., 1994; Pointon et al., 2012). In cladodograms in which Palaeodictyopterida is the fossil sister group of Neoptera (Sroka et al., 2015), palaeopterous wings are presumably a symplesiomorphy. The presence of nymphal wing pads is probably also a symplesiomorphy of Pterygota (Haug et al., 2016). Therefore Palaeodictyopterida may be outside the crown and even stem group of Palaeoptera.

Paoliida is an extinct clade known mainly from wings of the Westphalian A stage of the Pennsylvanian. The group has been attributed to the Palaeodictyoptera and later removed (Prokop et al., 2012; Prokop and Nel, 2007). It has been subsequently proposed that Paoliida is the fossil sister group of Dictyoptera (Prokop et al., 2014), and thus are within crown Polyneoptera. This would refute a relationship with Palaeoptera or any of its members.

A full body impression of a putative Carboniferous (Westphalian B/C) mayfly is subject to controversy (Benner et al., 2013; Knecht et al., 2011; Marden, 2013a, 2013b). *Bojophlebia prokopi* Kukalová-Peck, 1985 (Westphalian C) is likely outside of Palaeoptera, according to descriptions and phylogenetic analysis (Prokop et al., 2010; Sroka et al., 2015; Staniczek et al., 2011). The Stephanian B/C fossil *Triplosoba pulchella* Brongniart, 1883, originally described as a mayfly, has been redescribed as Palaeodictyopterida (Prokop and Nel, 2009). The oldest body fossils belonging to the mayfly stem group are therefore members of the Syntonopteroidea. The oldest Syntonopteroidea are *Lithoneura lameerei* Carpenter, 1938 and *Syntonoptera schucherti* Handlirsch, 1911, from the Westphalian D Mazon Creek (younger than Xiaheyan) (Nicholson et al., 2015; Prokop et al., 2010).

Putative older members of Odonatoptera are Geroptera, represented by the species *Eugeropteron lunatum* and *Geropteron arcuatum* (both Riek and Kukalová-Peck, 1984), are known from the upper Namurian of Argentina (Gutierrez et al., 2000; Riek and Kukalová-Peck, 1984). *G. arcuatum* was included in a morphological phylogenetic analysis, but was resolved outside Protodonata, in a polytomy with Odonata, Ephemeroptera, and Neoptera, thus outside Palaeoptera (Bybee et al., 2008). It need, however, be noted that morphological characters were polarized *a priori* with respect to *G. arcuatum* (Bybee et al., 2008).

## 72. Crown Euplectoptera

This clade comprises the families Siphluriscidae, Baetidae, Isonychiidae, Ametropodidae, Acanthametropodidae, Coloburiscidae, Siphlaenigmatidae, Ameletopsidae, Heptageniidae, Metretopodidae, Siphlonuridae, Oniscigastridae, Rallidentidae, Nesameletidae, and the larger clades Carapacea and Furcatergalia, their last common ancestor and all of its descendants (Fig. 17). A variety of different classification schemes have been proposed (Kluge, 2004, 1998; McCafferty, 1991; Ogden and Whiting, 2005), but we follow the consensus based on combined phylogenetic analysis of two nuclear genes, two mitochondrial genes, and morphology (Ogden et al., 2009). Although the extant members are often referred to as Ephemeroptera, that clade includes a number of extinct members forming a paraphyletic grade (Kluge, 2004, 1998). As membership of fossil species within a crown group is only possible if they branch along the stem lineage of a living clade that is part of the crown itself, we treat crown ‘Ephemeroptera’ as equivalent to the clade Euplectoptera *sensu* Tillyard (1932).

Monophyly has been supported by the aforementioned combined data study (Ogden et al., 2009), as well as analyses of transcriptomes (with limited but representative taxon sampling: Misof et al., 2014) and morphology (Ogden et al., 2009 Fig. 4: note this is monophyletic, but the root is shown intervening).

### 72.1. *Fossil specimens*

*Triassonurus doliiformis* Sinitshenkova et al., 2005 (part of the private Louis Grauvogel collection, Ringendorf, Bas-Rhin, France, No. 9304), holotype, an incomplete nymph (Fig. 18e).

### 72.2. *Phylogenetic justification*

*T. doliiformis* has not been explicitly included in a phylogenetic analysis. It bears several morphological characters indicating affinity with the extant family Siphlonuridae: a large nymph with a non-flattened body, head longer than short prothorax, massive mesothorax with considerably shorter metathorax, large and wide forewing pads almost completely covering the hind ones, short and slender legs, abdominal segments without sharp denticles, large rounded tergaliae, and cerci and paracercus long (Sinitshenkova et al., 2005). The family Siphlonuridae is not accepted as monophyletic by all authors (J.-D. Huang et al., 2008; Kluge, 2004; McCafferty, 1991) although it is supported in total evidence and morphological analyses of Ogden et al. (2009). Siphlonuridae is within the crown of Euplectoptera, in the clade of families outside Carapacea + Furcatergalia (Ogden et al., 2009). Therefore, *T. doliiformis* is also crown Euplectoptera.

### 72.3. *Age justification*

The fossil is derived from the silt-clay Grès à Meules unit (lowermost layer) of the Grès-a-Voltzia Formation, Vosges, northeastern France (Gall and Grauvogel-Stamm, 1993). Grès à Meules represents the last stage of the fluviatile facies prevalent in the Buntsandstein (Gall, 1985). Based on sequence stratigraphy, Bourquin et al. (2007, 2006) correlate the Grès-a-Voltzia Formation to the middle Anisian stage of the Triassic. Globally, the uppermost boundary of the Anisian is 241.5 ± 1 Ma (Ogg, 2012), providing a minimum age at 240.5 Ma.

Because the monophyly of Palaeoptera is under debate (Simon et al., 2012, 2009), we assign a soft maximum age that allows its constituent extant orders to be older than the Rhynie Chert, as for 26.3.

### 72.4. *Discussion*

The Permian fossils *Protereisma permianum* Sellards, 1907 and *Kukalova americana* Demoulin, 1970 were coded in the morphology matrix (larval and adult characters) of Ogden et al. (2009), but their relationships to the crown remain uncertain. *P. permianum* was resolved on the stem lineage of the extant families Euthyplociidae, Ephemeridae, and Potamanthidae with morphology only, but outside crown Euplectoptera in a total evidence analysis (Ogden et al., 2009). More generally, Kluge (Kluge, 2004, 1998) considered the Permian-Triassic members to form a plesion, Permoplectoptera, outside the crown group Euplectoptera. The relationship of Permoplectoptera to the crown group remains to be tested by morphological phylogenetics.

Furthermore, many mayfly fossils are of nymphs, and linking characters between fossil and extant nymphs (or fossil nymphs and extant adults) is challenging, though not impossible (e.g. Haug et al., 2016; Wolfe and Hegna, 2014). The nymphal fossils include *Fuyous gregarius* and *Shantous lacustris* (both Zhang and Kluge, 2007), two species from the Jurassic Daohugou biota that have been used as crown group calibrations in previous divergence time analyses (Thomas et al., 2013). They are, however, younger than *T. doliiformis.*

## 73. Crown Odonata

This clade comprises Epiprocta (dragonflies; comprising Anisozygoptera and Anisoptera) and Zygoptera (damselflies), their last common ancestor and all of its descendants (Fig. 17). Monophyly of Odonata is supported by phylogenetic analysis of transcriptomes (Misof et al., 2014), combined morphology and housekeeping genes (Bybee et al., 2008), and a supertree including morphological and molecular input trees (Davis et al., 2011).

### 73.1. *Fossil specimens*

*Triassothemis mendozensis* Carpenter, 1960. MACN No. 18040, holotype, preserving the distal portion of two wings.

### 73.2. *Phylogenetic justification*

*T. mendozensis* is the oldest known member of the fossil family Triassolestidae (Nel et al., 2002; Nicholson et al., 2015). A family-level supertree, incorporating molecular and morphological input trees, found Triassolestidae within crown group Epiprocta (Davis et al., 2011, largest tree in their Fig. 1). This fossil is therefore also a member of crown group Odonata. Of all the fossil families included in the supertree analysis and placed within crown Odonata (Davis et al., 2011), Triassolestidae (represented by *T. mendozensis)* has the oldest member. Furthermore, this family (and its approximate date) was used to calibrate Odonata in multiple recent divergence time analyses (Rota-Stabelli et al., 2013a; Thomas et al., 2013).

### 73.3. *Age justification*

*T. mendozensis* was discovered in dark grey siltstone of the Potrerillos Formation, at Quebrada del Durazno, Mendoza Province, Argentina (Martins-Neto et al., 2008). The insect-bearing beds are from the upper part of the Potrerillos Formation. U-Pb SHRIMP dates have been measured for zircons from tuff layers in the middle section of the Potrerillos Formation. The uppermost tuff layer estimated an age of 230.3 Ma ± 2.3 Myr (Spalletti et al., 2009), corresponding to the Carnian, late Triassic. Thus a minimum age of *T. mendozensis* is 228.0 Ma.

Because the monophyly of Palaeoptera is under debate (Simon et al., 2009, 2012), we assign a soft maximum age that allows its constituent extant orders to be older than the Rhynie Chert, as for 26.3.

### 73.4. *Discussion*

The internal taxonomy and placement of odonate fossils is highly contentious. Triassic members of the fossil family Triadophlebiidae are of approximately equal age to *T. mendozensis* (i.e. Carnian; Nicholson et al., 2015), however, they appear outside the crown group of Odonata in a supertree (Davis et al., 2011). Triadophlebiidae were not included in the total evidence analysis of Bybee et al. (2008). Most fossils in the latter analysis that had membership within crown Odonata were Jurassic or younger (Bybee et al., 2008).

The Madygen fauna of Kyrgyzstan yields two possible calibrations for crown group Odonata. *Triassolestodes asiaticus* Pritykina, 1981 (family Triassolestidae, preserving a hindwing) was recently justified as the oldest odonate by Kohli et al. (2016). As well, fossil members of Lestidae, crown group Odonata in the analysis of Bybee et al. (2008), are inferred from oviposition scars on plant fossils from Madygen (Moisan et al., 2012). Aside from challenges associated with interpreting crown group affinities of trace fossils, the Madygen fauna is, according to our stratigraphy, dated to the Carnian (see section 90.3), substantially younger than the 237 Ma age argued by Kohli et al. (2016) based on megaflora. Our age would give a minimum age of 226.4 Ma for Madygen fossils, which is very slightly younger than the 228.0 Ma estimated for *T. mendozensis.* Note, however, that the mean age estimate for Madygen is 228.4 Ma (also younger than the mean of 230.3 Ma for the Potrerillos Formation).

*Triassolestes epiophlebioides* Tillyard, 1918, a member of Triassolestidae used as a calibration fossil by Rota-Stabelli et al. (2013a), is also known from the Carnian (Nicholson et al., 2015). Another fossil, *Pseudotriassothemis nipponensis* Bechly, 1997 (formerly *Triassoneura okafujii),* from the Carnian of Japan is also approximately coeval. We select *T. mendozensis* because its preservation is better, and radiometric dates for the Potrerillos Formation may be more precise.

## 74. Crown Neoptera

This clade comprises Polyneoptera (Figs. 19 and 20) and Eumetabola, their last common ancestor and all of its descendants (Fig. 21). Monophyly is supported by phylogenetic analysis of transcriptomes (Letsch and Simon, 2013; Misof et al., 2014), protein-coding genes (Ishiwata et al., 2011; Sasaki et al., 2013), and combined molecular and morphological data (Terry and Whiting, 2005).

**Fig. 19.**
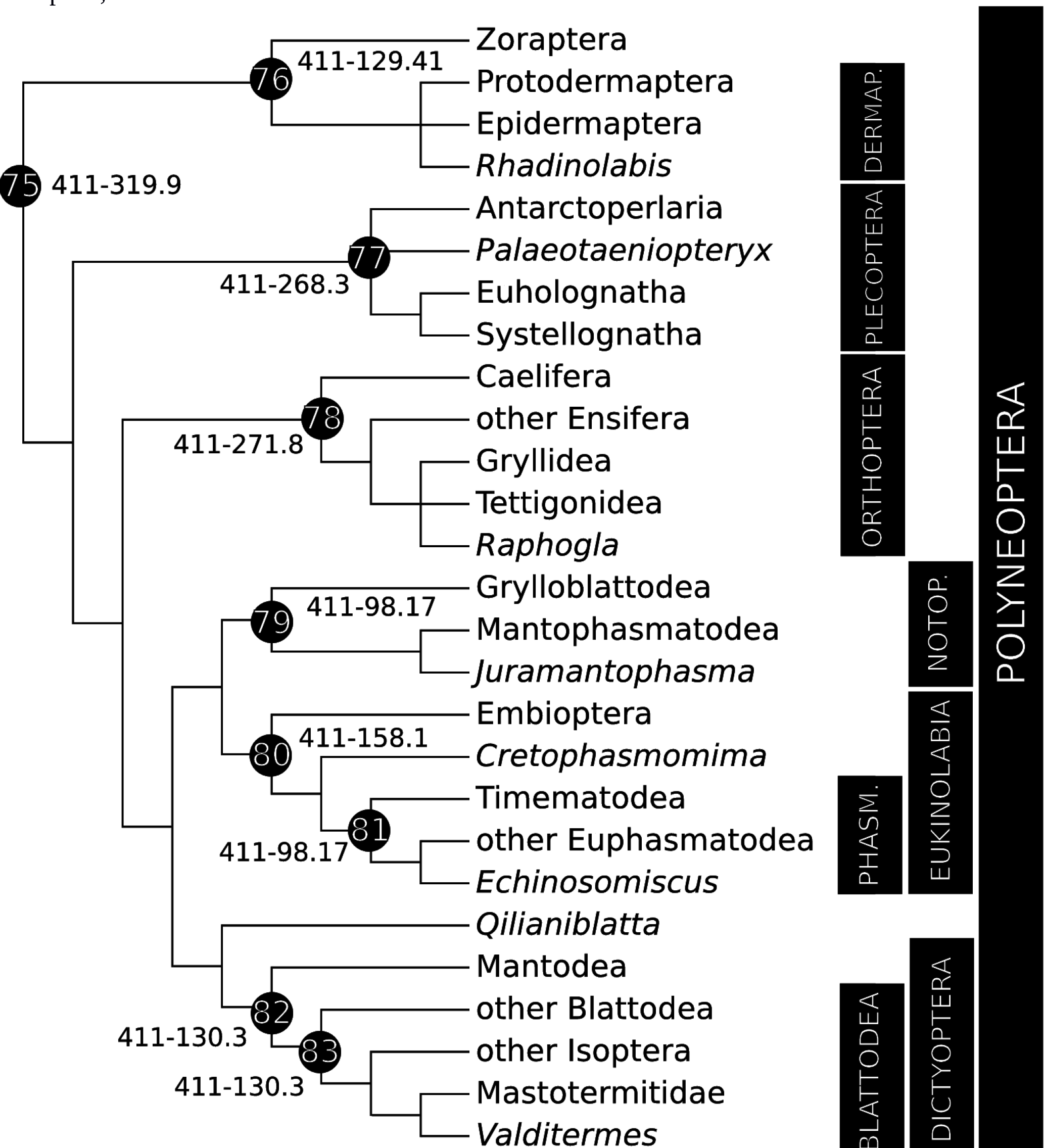
Calibration diagram for Polyneoptera (nodes 75–83). Dermap. = Dermaptera, Notop. = Notoptera, Phasm. = Phasmatodea.

**Fig. 20.**
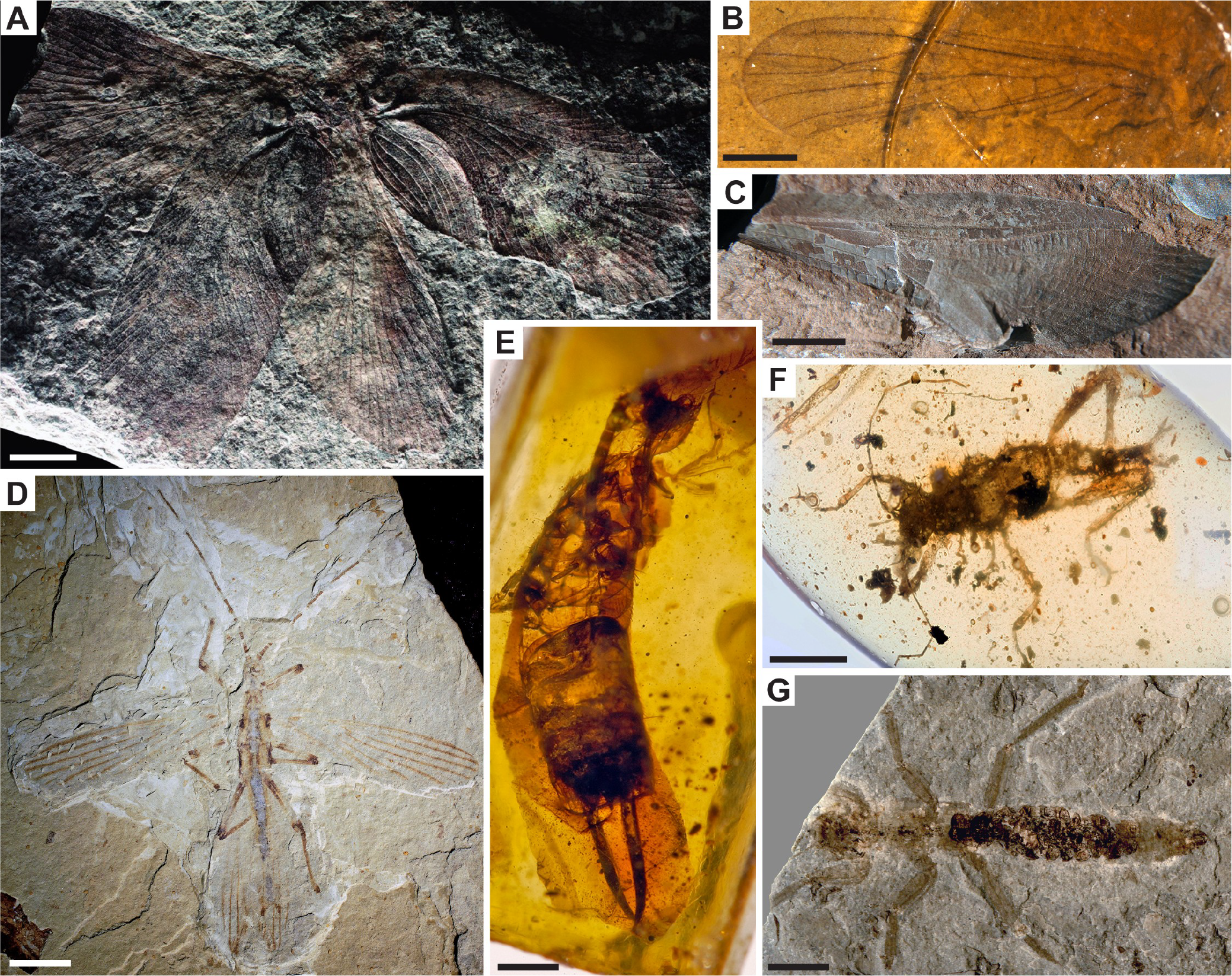
Polyneoptera fossil calibrations for (A) node 75: *Qilianiblatta namurensis,* CNU-NX1–303, scale bar 5mm, image credit D. Ren; (B) node 77: *Palaeotaeniopteryx elegans,* PIN 1197/333, scale bar 1mm, image credit N. Sinitshenkova; (C) node 78: *Raphogla rubra,* Ld LAP 415 B, scale bar 5 mm, image credit S. Fouche; (D) node 80: *Cretophasmomima melanogramma,* CNU-PHA-NN2012002, scale bar 10mm, image credit D. Ren; (E) node 76: *Rhadinolabis phoenicica,* Azar collection 1013, scale bar 500μm, image credit M. Engel;; (F) node 81: *Echinosomiscus primoticus,* NIGP 163536, scale bar 2mm, image credit M. Engel; (G) node 79: *Juramantophasma sinica,* NIGP 142171, scale bar 10mm, image credit D. Huang.

**Fig. 21.**
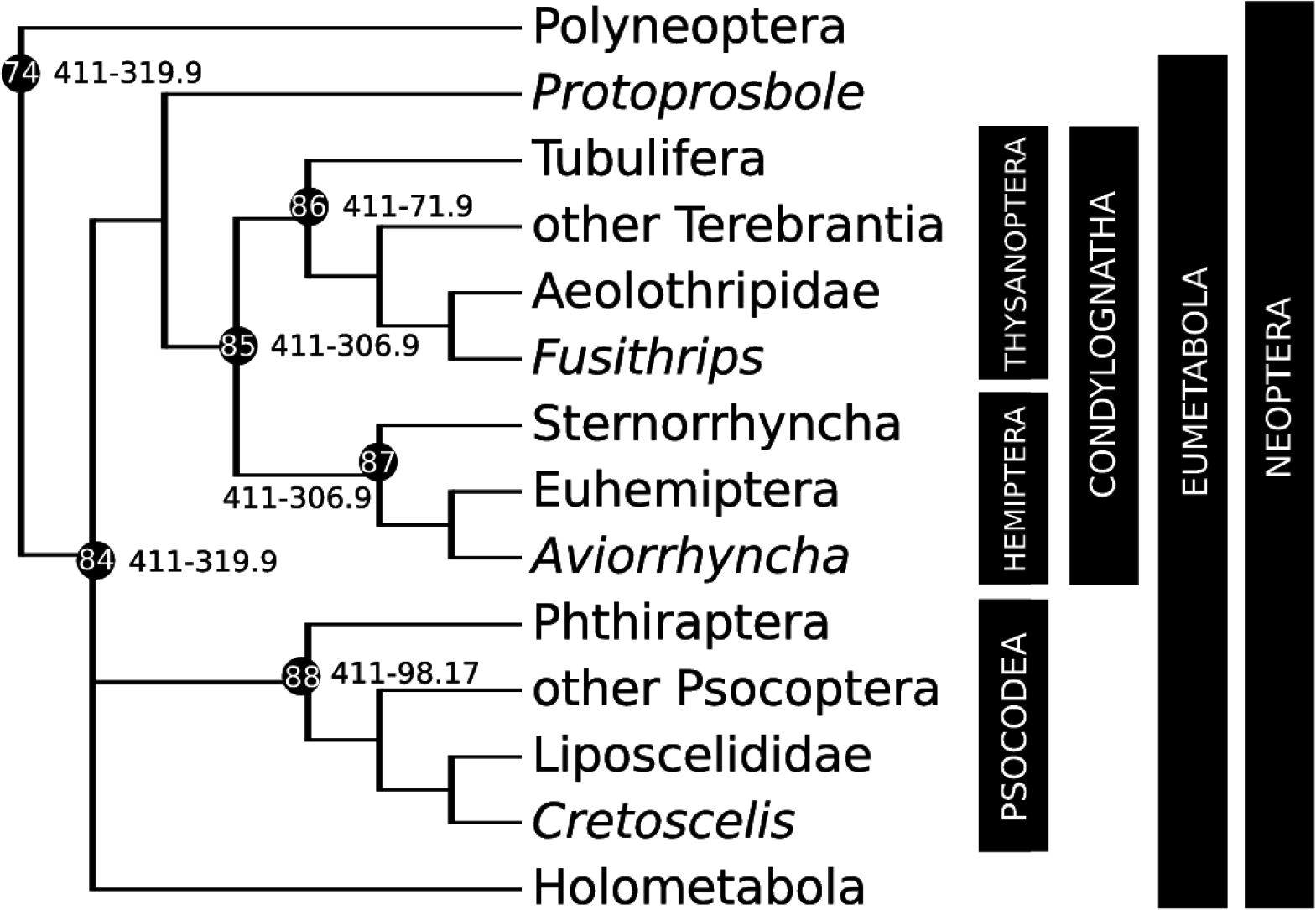
Calibration diagram for Condylognatha and Psocodea (nodes 74, 84–88).

### 74.1. *Fossil specimens*

*Protoprosbole straeleni* Laurentiaux, 1952. IRSNB a9885, holotype, preserving a forewing. Figured in Nel et al. (A. Nel et al., 2012), Fig. 3h.

### 74.2. *Phylogenetic justification*

The original description of *P. straeleni* as a member of Hemiptera by Laurentiaux (1952) has been rejected several times (Hennig, 1981; A. Nel et al., 2012; Shcherbakov, 1995). Nel et al. (2012) summarize the phylogenetic position of *P. straeleni* as being within Paraneoptera (a clade unsupported by recent molecular studies, but comprising Psocodea and Condylognatha). The cua-cup contact with CuP and the flexion or nodal line following the course of RA are both similar to those observed in extant Hemiptera (A. Nel et al., 2012). However, the presence of three veins in the anal area is argued as a hemipteran autapomorphy that is lacking in *P. straeleni* (A. Nel et al., 2012). Conservatively, this fossil species can be thus assigned to the stem group of Condylognatha, and thus crown Eumetabola and Neoptera.

### 74.3. *Age justification*

*P. straeleni* was discovered in Charbonnage de Monceau-Fontaine, Charleroi Coal Basin, Belgium (Brauckmann et al., 1994). The specimen was likely found in uppermost Marsdenian strata about 3 m below the base of the Yeadonian (discussed by Brauckmann et al., 1994). The fossil-bearing deposits are assigned to the late Namurian B (Marsdenian) based on the *Bilinguites superbilinguis* R2c2 subzone of goniatite ammonoid stratigraphy (Brauckmann et al., 1994). The (upper) Namurian-(lower) Westphalian boundary is defined by the earliest occurrence of the goniatite *Gastrioceras subcrenatum* (Waters and Davies, 2006), but lacks a precise isotopic date. Pointon et al. (2012) estimated an age of c. 319.9 Ma for the base of the Westphalian (top of the Namurian, only slightly younger than the Marsdenian) based on Milankovitch cycles of sedimentation, giving a minimum age for *P. straeleni.*

Soft maximum as for 26.3.

### 74.4. *Discussion*

*Qilianiblatta namurensis* Zhang et al., 2013 has an equal claim to being the earliest record of Neoptera (as *Q. namurensis* is a member of crown Polyneoptera), but it is of equal age to *P. straeleni*. The Qilianshan entomofauna at Xiaheyan, China, yields many other likely neopterans (e.g. Béthoux et al., 2011; Liu et al., 2009a; Prokop and Nel, 2007; Zhang et al., 2013).

*Ctenoptilus elongatus* Béthoux and Nel, 2005 from the Stephanian B/C of France has been used as the putative oldest calibration fossil for crown Neoptera (e.g. O’Reilly et al., 2015). However, the Stephanian is a European stage of the Pennsylvanian, corresponding to the globally used Gzhelian, which is substantially younger than the Bashkirian age of both *P. straeleni* and *Q. namurensis* (with an upper bound of 298.75 Ma).

## 75. Crown Polyneoptera

This clade comprises Zoraptera (ground lice), Dermaptera (earwigs), Plecoptera (stoneflies), Orthoptera (crickets, katydids), Notoptera, Eukinolabia and Dictyoptera, their last common ancestor and all of its descendants (Fig. 19). Monophyly has been demonstrated by phylogenetic analysis of transcriptomes (Letsch and Simon, 2013; Misof et al., 2014), protein-coding genes (Ishiwata et al., 2011), and wing morphology (Yoshizawa, 2011). This clade is not recognized in GenBank taxonomy.

### 75.1. *Fossil specimens*

*Qilianiblatta namurensis* Zhang et al., 2013. GMCB 04GNX1001–1, holotype, preserving the right forewing. We also refer to a second specimen (CNU-NX1–303; Fig. 20a), preserving paired forewings and hindwings (Guo et al., 2013).

### 75.2. *Phylogenetic justification*

*Q. namurensis* has not been included in formal phylogenetic analysis, but the fossil exhibits some apomorphic characters uniting it with crown group Blattodea, specifically presence of a deeply concave CuP in the forewing (Prokop et al., 2014). Additional data from forewings of the CNU specimen shows the RA with branches translocated to RP, as in cockroaches, suggesting this species could be stem Blattodea (Guo et al., 2013). However, there has not been a cladistic analysis of wing venation characters for both fossil and extant Blattodea and/or Dictyoptera, thus we agree with the more conservative view (Kjer et al., 2015; Legendre et al., 2015; Prokop et al., 2014) that roachoids likely represent a diverse and speciose fauna on the stem group of Dictyoptera. See also section 82.4. Nonetheless, placement of *Q. namurensis* on the stem lineage of Dictyoptera identifies it as a member of crown Polyneoptera.

### 75.3. *Age justification*

Minimum as for 71.3.

A soft maximum age is estimated from *R. praecursor,* the oldest hexapod, from the Lower Devonian (Pragian) Rhynie Chert of Aberdeenshire, Scotland. Spore assemblages of the Windyfield and stratigraphically underlying Rhynie Chert are dated to the early but not earliest Pragian to early (earliest?) Emsian (polygonalis-emsiensis Spore Assemblage Biozone) (Parry et al., 2011). Radiometric dating of the underlying Milton of Noth Andesite at ca. 411 Ma (Parry et al., 2013, 2011) has been subject to a dispute over its temporal relationship to hot spring activity associated with the cherts (Mark et al., 2013, 2011) and predates the biostratigraphic dating of the Rhynie Chert relative to the global dating of the base of the Pragian Stage. Therefore, a soft maximum constraint may be defined at 411 Ma for the Rhynie Chert.

## 76. Crown Dermaptera

This clade comprises Protodermaptera and Epidermaptera, their last common ancestor and all of its descendants (Fig. 19). Monophyly is supported based on phylogenetic analysis of transcriptomes (Misof et al., 2014), nuclear genes (Kocarek et al., 2013), and combined molecular and morphological data (Jarvis et al., 2005).

### 76.1. *Fossil specimens*

*Rhadinolabis phoenicica* Engel et al., 2011. Holotype preserving a female in amber, 1013 in the private collection of D. Azar in Fanar (Matn), Lebanon (Fig. 20e). Provisionally housed at MNHN.

### 76.2. *Phylogenetic justification*

*R. phoenicica* is assigned only to Neodermaptera (= Protodermaptera + Epidermaptera), and not to any extant family. Membership within Neodermaptera is based on the presence of morphological characters: trimerous tarsi, absence of a well-developed ovipositor, and tarsal structure (Engel et al., 2011). These characters are sufficient to confirm the fossil belongs within crown Dermaptera.

### 76.3. *Age justification*

This fossil was discovered in amber from the Mdeyrij-Hammana outcrop of the Baabda district, Mount Lebanon (Azar et al., 2010). Recent revision of the stratigraphy of Lebanese amber deposits places the Hammana fossils in the upper interval of the Grès du Liban (Maksoud et al., 2016). This is below the Jezzinian regional stage (Maksoud et al., 2014) and above the Banc de Mrejatt subunit (indicated as Ba3-Ba4 in Fig. 4 of Maksoud et al., 2016). Despite the lack of microfossils to further constrain the oldest boundary of the Jezzinian within the late Barremian, there is evidence that later Lebanese amber deposits bear the same age as Jezzine amber (see 26.3) because the amber itself has been reworked (Maksoud et al., 2016). We adopt the early Barremian minimum age proposed by Maksoud et al. (2016). The upper boundary of the early Barremian is proposed to be the first appearance of the ammonite *Ancyloceras vandenheckii* (Ogg et al., 2012). Cyclostratigraphy dates the *A vandenheckii* Zone beginning at 129.41 Ma (Ogg et al., 2012), providing a minimum age for Jezzine Lebanese amber fossils.

Soft maximum as for 75.3.

## 77. Crown Plecoptera

This clade comprises Antarctoperlaria, Euholognatha and Systellognatha, their last common ancestor and all of its descendants (Fig. 19). This classification and its morphological evidence is reviewed by Zwick (2000). Monophyly of Plecoptera is supported by phylogenetic analysis of transcriptomes (Misof et al., 2014) and combined ribosomal genes, H3, and morphology (Terry and Whiting, 2005).

### 77.1. *Fossil specimens*

*Palaeotaeniopteryx elegans* Sharov, 1961. PIN 1197/333, holotype, preserving a forewing.

### 77.2. *Phylogenetic justification*

*P. elegans* is classified in the fossil family Palaeonemouridae, suborder Nemourina (Sinitshenkova, 1987). At that time, Nemourina was one of only two suborders, the other being Perlina (Rasnitsyn and Quicke, 2002). Zwick (2000) summarized Nemourina as containing the (extant) families Notonemouridae, Nemouridae, Taeniopterygidae, Capniidae, and Leuctridae, therefore equivalent to the new suborder Euholognatha. Grimaldi and Engel (2005; Fig. 7.6) prefer a placement of Palaeonemouridae outside the crown group of Plecoptera, on the basis that diagnostic characters for crown group suborders within Plecoptera are rarely preserved (e.g. muscles, cellular structures). While no formal phylogenetic hypothesis illustrates the relationship of Palaeonemouridae to other Euholognatha, synapomorphies are identified linking *P. elegans* to the crown group of Plecoptera (Béthoux, 2005, pers. comm.).

### 77.3. *Age justification*

The oldest specimen of *P. elegans* was discovered from the Mitino Horizon at the Kaltan locality in the Kuznetsk Basin, Kemerovo, Russia (Sharov, 1961; Shcherbakov, 2000). Correlation of insect faunas suggests the Kaltan locality is late early Kazanian (Lozovsky et al., 2009; Shcherbakov, 2008). The Kazanian is a regional stage of the Russian Permian, which has been correlated to both the Wordian (Menning et al., 2006) and the Roadian global Stages (Lozovsky et al., 2009). Evidence for an age in the Roadian is stronger, demarcated by presence of ammonites such as *Sverdrupites harkeri* and *S. amundseni* (Lozovsky et al., 2009). These ammonites, and associated conodonts *Kamagnathus khalimbadzhai* and *K. volgensis,* correlate globally to the Roadian (Barskov et al., 2014; Leonova, 2007; Leonova and Shilovsky, 2007). The upper boundary of the Roadian is 268.8 Ma ± 0.5 Myr, so a minimum age is 268.3 Ma.

Soft maximum as for 75.3.

### 77.4. *Discussion*

*Gulou carpenteri* Béthoux et al., 2011, from the Pennsylvanian Qilianshan entomofauna of China, is identified as a member of the stem group to Plecoptera (Béthoux et al., 2011), and thus cannot be used to calibrate a minimum age of the crown group. Moreover, other Permian plecopterans had terrestrial nymphs; an aquatic nymph is a symapomorphy uniting crown group Plecoptera (Zwick, 2000), relegating any fossil without it to the stem group.

O. Béthoux (pers. comm.) cautioned that a Jurassic minimum age for crown Plecoptera, as used by Misof et al. (2014), would underestimate their age significantly. The calibrating fossil *Pronemoura angustithorax* Liu et al., 2011 used by Misof et al. (2014), from the Daohugou Beds, is likely a member of total group Euholognatha (Liu et al., 2011). Other crown Plecoptera groups are also known from the same locality (Liu et al., 2009b), suggesting diversification of the crown group was significantly earlier.

## 78. Crown Orthoptera

This clade comprises Ensifera (crickets, katydids) and Caelifera (grasshoppers), their last common ancestor and all of its descendants (Fig. 19). Monophyly is demonstrated by phylogenetic analysis of transcriptomes (Misof et al., 2014), mitochondrial genomes plus four nuclear genes (Song et al., 2015), morphology (Béthoux and Nel, 2002), and combined molecular and morphological data (Terry and Whiting, 2005).

### 78.1. *Fossil specimens*

*Raphogla rubra* Béthoux et al., 2002. Ld LAP 415 B, holotype, preserving the counterpart of a forewing (Fig. 20c).

### 78.2. *Phylogenetic justification*

Although no cladistic analysis exists combining extant and fossil Enserifa, *R. rubra* may be assigned to crown Enserifa based on forewing venation characters (Béthoux et al., 2002). It likely belongs to the stem group of the clade (Gryllidea + Tettigoniidea), based on the following characters: very broad area between anterior margin and Sc; RS moderately long basal of a short fusion with the anterior branch MA1a of MA; MP + CuA1 with only one simple anterior branch (Béthoux et al., 2002). As both Gryllidea and Tettigoniidea are crown clades within crown Enserifa, *R. rubra* is within crown group Orthoptera.

### 78.3. *Age justification*

*R. rubra* was found at the fossiliferous site F21 D, at “Le Moural D”, in the basal Merifrons Member of the Salagou Formation (Saxonian Group), near village Octon, Lodeve Basin, Herault, France (Béthoux et al., 2002; Nel et al., 1999; Schneider et al., 2006). U-Pb dates have been recently measured from zircons lying in tuff beds near the lower two-thirds of the Salagou Formation (Octon Member), in the Artinskian (Michel et al., 2015). The Merifrons Member, overlying the Octon, is therefore inferred to be Artinskian at its base but continuing up through the Kungurian (Michel et al., 2015). As the exact stratum of the F21 site is not indicated within the Merifrons member (Michel et al., 2015; Schneider et al., 2006), we apply a conservative minimum estimate from the upper bound of the Kungurian (272.3 Ma ± 0.5 Myr), at 271.8 Ma.

Soft maximum as for 75.3.

### 78.4. *Discussion*

An undescribed Namurian member of Archaeorthoptera was initially attributed to crown group Orthoptera (Prokop et al., 2005). However, an affinity with Archaeorthoptera is supported in the stem group of Orthoptera based on phylogenetic analysis (Béthoux and Nel, 2002), thus it cannot be used to calibrate the crown. Similarly, *Oedischia williamsoni* Brongniart, 1885, from the Pennsylvanian of France, is recognized as a stem group member of Orthoptera by cladistic analysis (Béthoux and Nel, 2002). Together these fossils suggest a long stem branch for Orthoptera.

The fossil *Eolocustopsis primitiva* Riek, 1976 from Natal, South Africa was listed as the oldest crown group member of Caelifera, and thus another Permian crown group member of Orthoptera (Song et al., 2015). However, the fossil comes from strata of the Normandien Formation, Changhsingian stage, latest Permian (due to the co-occurrence of the vertebrate fossil *Dicynodon;* e.g. Catuneanu et al., 2005), which is younger than the Kungurian.

## 79. Crown Notoptera

This clade comprises Mantophasmatodea (rock crawlers) and Grylloblattodea (ice crawlers), their last common ancestor and all of its descendants (Fig. 19). The monophyly of Notoptera is supported by phylogenetic analysis of transcriptomes (Misof et al., 2014), morphology (Wipfler et al., 2011), and combined molecular and morphological data (Terry and Whiting, 2005).

### 79.1. *Fossil specimens*

*Juramantophasma sinica* D. Huang et al., 2008. NIGP 142171, holotype, preserving a nearly complete adult female (Fig. 20g).

### 79.2. *Phylogenetic justification*

*J. sinica* has several characters in common with crown Mantophasmatodea. These include: a third tarsomere with a sclerotized elongated dorsal process, enlarged and fanlike pretarsal arolia, last tarsomere at right angle to the others, female gonoplacs short and claw shaped, and egg with a circular ridge (D. Huang et al., 2008). As no morphological matrix exists for Mantophasmatodea, relationships to extant lineages are not possible to test (D. Huang et al., 2008). The fossil is excluded from the crown group of Grylloblattodea as it lacks segmented cerci. Thus, conservatively, a position on the stem lineage of Mantophasmatodea is likely (although *J. sinica* could be amended to within the crown of Mantophasmatodea). This would, in any case, mean it is a member of crown Notoptera.

### 79.3. *Age justification*

This fossil was found in deposits of the Daohugou Beds, Ningcheng County, Inner Mongolia, China (D. Huang et al., 2008). There has been controversy about the accuracy and precise age and stratigraphic position of the Daohugou Beds (Gao and Ren, 2006; Wang et al., 2005; Zhang, 2015). The beds consist of 100–150 m thick succession of grey-white or locally reddish, thinly bedded claystones, shales, siltstones and sandy mudstones with intercalated ash-fall tuffs and ignimbrites. Ages have been proposed from Aalenian (Middle Jurassic) to Early Cretaceous (Liu et al., 2014; Wang et al., 2000), with several studies converging on Callovian-Oxfordian (Late Jurassic; Zhang, 2015). Radiometric dating of the ignimbrites with 40Ar/39Ar and SHRIMP U-Pb variously yields dates between 165 Ma ± 2.5 Myr and 158.7 Ma ± 0.6 Myr (Chang et al., 2009b; Gao and Ren, 2006; He et al., 2004; Peng et al., 2012). The fossiliferous shales overlay the volcanic deposits (Gao and Ren, 2006), and are thus younger. The isotopic dates nonetheless provide a reasonable refutation of Cretaceous age estimates. Furthermore, the Daohugou Beds may be correlated to sediments from Oxfordian localities in China and Kazakhstan (Zhang, 2015). The most conservative (i.e. youngest) of the direct radiometric dates is 158.1 Ma (within the Oxfordian), giving a minimum age.

Soft maximum as for 75.3.

### 79.4. *Discussion*

Stem group Grylloblattodea are uncommon, and extant grylloblattids (and mantophasmids) are wingless (Wipfler et al., 2014). A putative grylloblattid insect was described from the Pennsylvanian Montceau-les-Mines Lagerstatte, about 130 Myr older than *J. sinica* (Béthoux and Nel, 2010); however, the fossil preserved only the wings. We take the parsimonious view that there was a single loss of wings in the common ancestor of Grylloblattodea and Mantophasmatodea, i.e. in the crown group of Notoptera (Grimaldi and Engel, 2005). Therefore, any fossil bearing wings, such as the Pennyslvanian and Permian members, would be on the stem branch of Notoptera.

## 80. Crown Eukinolabia

This clade comprises Embioptera (webspinners) and Phasmatodea (stick and leaf insects), their last common ancestor and all of its descendants (Fig. 19). Monophyly of Eukinolabia is supported by phylogenetic analyses of transcriptomes (Letsch and Simon, 2013; Misof et al., 2014) and ribosomal and H3 sequences (Terry and Whiting, 2005).

### 80.1. *Fossil specimens*

*Cretophasmomima melanogramma* Wang et al., 2014. CNU-PHA-NN2012002, holotype, preserving a male (Fig. 20d).

### 80.2. *Phylogenetic justification*

*C. melanogramma* shares the ’;shoulder pad’ structure, and twig-like appearance (elongated meso-and metathorax) diagnostic of total group Phasmatodea (Wang et al., 2014). Crown membership within Phasmatodea is questionable, as important synapomorphies, the vomer and forceps-like extensions of the 10th abdominal tergum, are absent from *C. melanogramma* (Wang et al., 2014). The position of *C. melanogramma* on the stem lineage of Phasmatodea therefore places it within the crown group of Eukinolabia.

### 80.3. *Age justification*

As for 76.3.

### 80.4. *Discussion*

*Renphasma sinica* Nel and Delfosse, 2011 is of equal age, also from the Yixian Formation, and also a member of stem Phasmatodea, and thus crown Eukinolabia.

## 81. Crown Phasmatodea

This clade comprises Timematodea and Euphasmatodea, their last common ancestor and all of its descendants (Fig. 19). Monophyly of Phasmatodea is supported by phylogenetic analyses of transcriptomes (Misof et al., 2014), ribosomal and H3 sequences (Terry and Whiting, 2005), and morphology (Friedemann et al., 2012).

### 81.1 *Fossil specimens*

*Echinosomiscus primoticus* Engel et al., 2016c. NIGP 163536, holotype, preserving a male in amber (Fig. 20f).

### 81.2 *Phylogenetic justification*

Until recently, no Mesozoic fossil qualified for membership in the crown group of Phasmatodea (Bradler and Buckley, 2011). *E. primoticus* bears morphological characters shared with extant members of Euphasmatodea, in particular Heteropterygidae and Aschiphasmatidae, but lack the area apicalis on the tibiae (Engel et al., 2016c). It also shares characters with Lonchodinae, such as absent areole on all tibiae, comparatively long antennae, the absence of wings, and the division of the tenth abdominal tergum into moveable hemitergites (Engel et al., 2016c). Although a new family was erected to accommodate the unique character combination for *E. primoticus*, it is very clearly within Euphasmatodea (Phasmatidae s.l.), and thus a member of crown Phasmatodea (Engel et al., 2016c).

### 81.3 *Age justification*

Minimum as for 12.3. Soft maximum as for 75.3.

## 82. Crown Dictyoptera

This clade comprises Mantodea (praying mantids) and Blattodea (cockroaches, termites), their last common ancestor and all of its descendants (Fig. 19). Phylogenetic analysis of transcriptomes (Misof et al., 2014), mitochondrial genes (Legendre et al., 2015), and combined molecular and morphological data (Djernæs et al., 2015; Ware et al., 2008), support monophyly of Dictyoptera.

### 82.1. *Fossil specimens*

*Valditermes brenanae* Jarzembowski, 1981. BMNH In. 64588, holotype, preserving a forewing.

### 82.2. *Phylogenetic justification*

The position of *V. brenanae* was confirmed by morphological phylogenetic analysis (Engel et al., 2009). *V. brenanae* was resolved on the stem lineage of Mastotermitidae (Engel et al., 2009). This position was separately found for a congeneric species in the combined morphological and molecular phylogeny of Ware et al. (2010). Mastotermitidae is a monotypic family, sister to all other termites (Djernas et al., 2015; Ware et al., 2010), so this phylogeny would also place *V. brenanae* in the crowns of Isoptera and Blattodea.

### 82.3. *Age justification*

The fossil is from the Clockhouse Brickworks pit site, Surrey, England (Jarzembowski, 1981). The Clockhouse Brickworks belongs to the Lower Weald Clay, as indicated by the presence of the ostracods *Cytheridea clavata, C. tuberculata,* and *C. valdensis* (Anderson, 1985). Of these, *C. tuberculata* has the narrowest range, and is restricted to the middle portion of the Lower Weald Clay, at BGS Bed 3a (Anderson, 1985; Ross and Cook, 1995). This faunal Zone is now assigned to the lower portion of the *C. pumila* Subzone of the *Theriosynoecum fittoni* Zone (Horne, 1995). Based on palynomorph stratigraphy, the boundary between Lower and Upper Weald Clay at the base of BGS Bed 3a corresponds accurately to the boundary between the Hauterivian and Barremian Stages (Ross and Cook, 1995). The younger boundary of the Hauterivian is 130.8 Ma ± 0.5 Myr (Ogg et al., 2012a), therefore a minimum age of the Lower Weald Clay is 130.3 Ma.

Soft maximum as for 75.3.

### 82.4. *Discussion*

There is ongoing debate over whether various Palaeozoic “roachoid” fossils, such as *Qilianiblatta namurensis* and *Homocladus grandis* Carpenter, 1966 (known only from wing venation characters), truly represent crown group members of Dictyoptera (e.g. Guan et al., 2016; Kjer et al., 2015; Legendre et al., 2015; Tong et al., 2015). If roachoids are indeed within crown group Dictyoptera, their antiquity may push back the origins of crown Dictyoptera and crown Polyneoptera by 80–88 Myr (Legendre et al., 2015; Tong et al., 2015). However, wing venation alone may be insufficient to place most fossils within the crown group of Blattodea or even Dictyoptera. Particularly in cockroaches, asymmetry at the individual level and convergence at higher taxonomic levels are impediments to wing venation taxonomy, as well as the paucity of comparative venation data for extant species (e.g. Béthoux et al., 2010; Béthoux and Wieland, 2009; Garwood and Sutton, 2010; Gorochov, 2013; Guan et al., 2016).

In the absence of detailed cladistic analysis of fossil roachoid wings together with extant Dictyoptera and other outgroups, we take the conservative view (Guan et al., 2016; Kjer et al., 2015; Prokop et al., 2014) that roachoids (including the family Anthracoptilidae/Strephocladidae, containing *H. grandis*) likely represent a diverse and speciose fauna on the stem group of Dictyoptera, perhaps with some members on the stem group of Neoptera or Polyneoptera (Grimaldi and Engel, 2005). At least some such roachoids may be used to calibrate crown group Polyneoptera (as done herein by assigning *Q. namurensis*), but they cannot calibrate crown Dictyoptera. As a consequence, many recent analyses have begun to rewrite the traditional assumption of extremely old crown ages for Dictyoptera, with crown origins likely in the Jurassic (Misof et al., 2014).

Morphological phylogenetic analyses have excluded all Cretaceous fossils from the crown group of Mantodea, although they remain as likely crown Dictyoptera (Delclòs et al., 2016; Grimaldi, 2003). Given that the analysed morphological matrix has fewer characters than taxa and has substantial missing data, it is not definitive (Delclòs et al., 2016). A recently discovered fossil from Crato (Aptian of Brazil), *Cretophotina santanensis* Lee, 2014, may be a stem group Chaeteessidae, which may be the most basal crown family of Mantodea (Svenson and Whiting, 2009). The position of Chaeteessidae may be within polyphyletic Mantidae (Legendre et al., 2015), but we exercise caution and do not place *C. santanensis* in the crown group, as recommended by Lee (2014).

## 83. Crown Blattodea

This clade comprises Lamproblattidae, Blaberoidea, Blattidae, Nocticolidae, Polyphagidae, Cryptocercidae and Isoptera, their last common ancestor and all of its descendants (Fig. 19). The position of Isoptera (termites) within a paraphyletic Blattodea was first identified by Inward et al. (2007) with six molecular loci. Further support for the monophyly of the new concept of Blattodea comes from transcriptomes (Misof et al., 2014), mitochondrial genes (Legendre et al., 2015), housekeeping genes (Djernæs et al., 2012), and combined molecular and morphological data (Djernæs et al., 2015; Ware et al., 2008).

### 83.1. *Fossil specimens*

As for 82.1.

### 83.2. *Phylogenetic justification*

As for 82.2.

### 83.3. *Age justification*

As for 82.3.

### 83.4. *Discussion*

*Baissatermes lapideus* Engel et al., 2007 is a crown group member of Isoptera, and *Mastotermes nepropadyom* Vrsansky and Aristov, 2014 is crown Blattodea (Engel et al., 2016a); both have been used to calibrate Isoptera and Blattodea (Djernæs et al., 2015; Ware et al., 2010). Note that fossil *Mastotermes* are likely polyphyletic (Engel et al., 2016a). Although these are appropriate calibration fossils, they are not the oldest Blattodea. *B. lapideus* is from the Zaza Formation (Transbaikalian Russia), and *M. nepropadyom* is from Chernovskie Kopi, also in Transbaikalia and similar in age to the Turga Formation (Sukatsheva and Vassilenko, 2011; Vrsansky and Aristov, 2014). The Turga Formation has been correlated to the Baissa locality, Zaza Formation; this is based on shared presence of *Asteropollis asteroides* spores (Vakhrameev and Kotova, 1977). As discussed in section 86.3, the Zaza Formation may be much younger than initially described, with a minimum age of Campanian (71.9 Ma). Therefore both Russian fossils are younger than *V brenanae.*

## 84. Crown Eumetabola

This clade comprises Condylognatha, Psocodea and Holometabola, their last common ancestor and all of its descendants (Fig. 21). Monophyly of the clade is supported by transcriptomes (Letsch and Simon, 2013; Misof et al., 2014) and protein-coding genes (Ishiwata et al., 2011). Some morphological analyses do not support monophyly of Eumetabola (Kristensen, 1981; Wheeler et al., 2001), although it is recovered (Kristensen, 1991; Kukalová-Peck, 1991) and assumed (Yoshizawa and Saigusa, 2001) in other analyses. Note also that support for Psocodea as the sister clade of Holometabola was statistically weak in analyses of transcriptomes (Misof et al., 2014), although monophyly of Eumetabola was well supported.

### 84.1. *Fossil specimens*

As for 74.1.

### 84.2. *Phylogenetic justification*

As for 74.2.

### 84.3. *Age justification*

Minimum as for 74.3. Soft maximum as for 75.3.

## 85. Crown Condylognatha

This clade comprises Hemiptera (true bugs) and Thysanoptera (thrips), their last common ancestor and all of its descendants (Fig. 21). Monophyly is determined by phylogenetic analysis of transcriptomes (Misof et al., 2014), nuclear protein-coding genes (Ishiwata et al., 2011), and wing morphology (Yoshizawa and Saigusa, 2001).

### 85.1. *Fossil specimens*

*Aviorrhyncha magnifica* Nel et al., 2013, holotype Avion No. 2 (provisionally stored in the collection of Entomological Laboratory, MNHN; to be deposited in the Musée Géologique Pierre Vetter, Decazeville, France), preserving a single forewing (Fig. 22a).

**Fig. 22.**
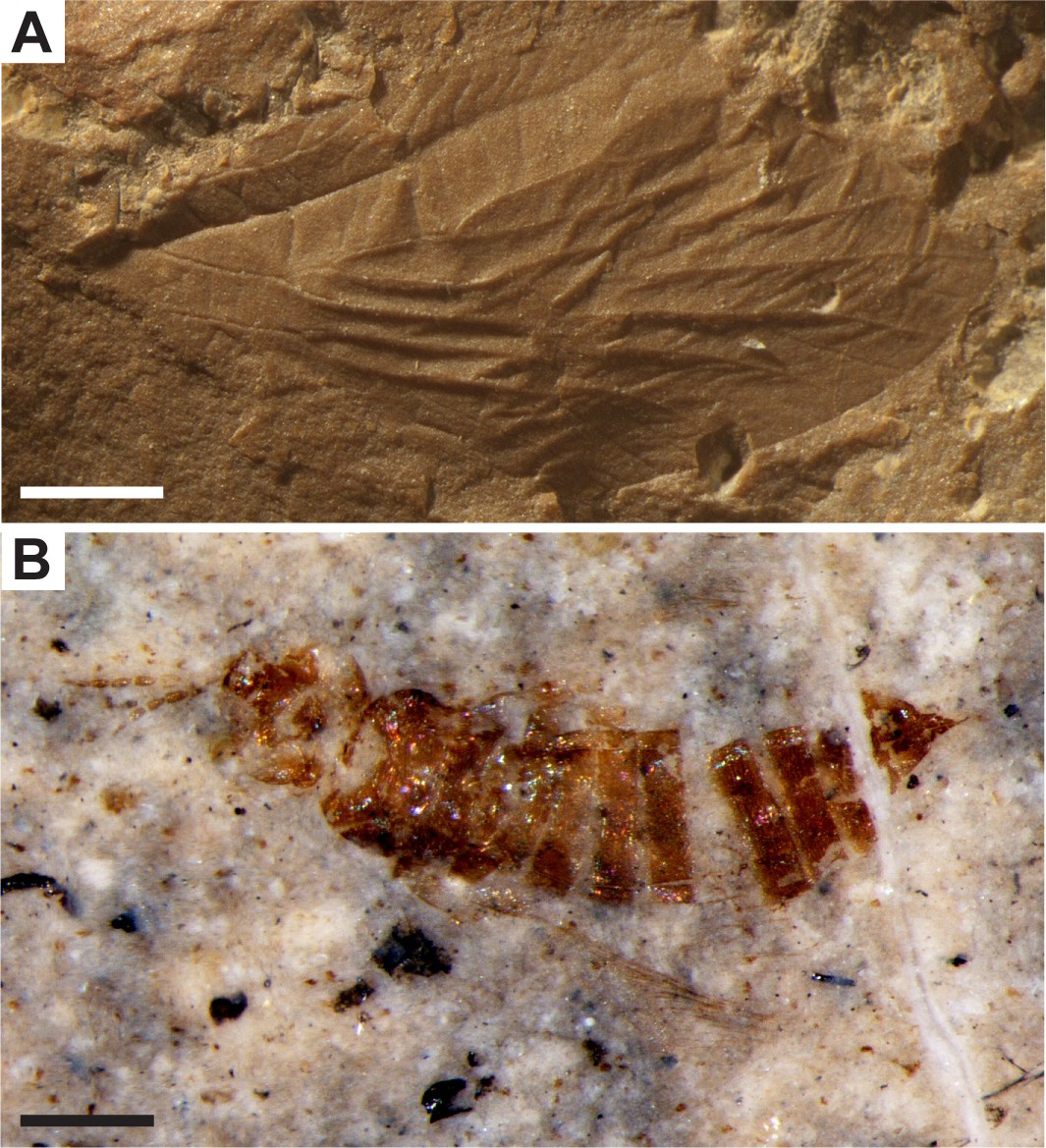
Condylognatha fossil calibrations for (A) nodes 85, 87: *Aviorrhyncha magnifica,* holotype MNHN Avion No. 2, scale bar 1mm, image credit A. Nel; (B) node 86: *Fusithrips crassipes,* PIN 3064/8547, scale bar 200pm, image credit A. Rasnitsyn.

### 85.2. *Phylogenetic justification*

Nel et al. (2013) diagnose *A magnifica* as stem Euhemiptera (the clade sister to Sternorrhyncha containing all other living Hemiptera; Cryan and Urban, 2012; Cui et al., 2013). At least two forewing characters in *A. magnifica* are synapomorphies of Euhemiptera, including presence of an ambient vein and a well-developed concave CP (its presence is a synapomorphy) (Nel et al., 2013). Unlike many extant Euhemiptera, the cua-cup vein is not zigzagged. Given that much of fossil insect taxonomy is conducted with only preserved wings, an assignment to the crown group of Hemiptera and thus Condylognatha is reasonable.

### 85.4. *Age justification*

*A magnifica* was found in “Terril No. 7”, a layer containing rocks from the slag heap of coal mines 3 and 4 of Liévin, in the Avion outcrop of Pas-de-Calais, France (Nel et al., 2013). The coal mines are dated to the Westphalian C/D, or Bolsovian/Asturian, which has a youngest boundary equivalent to the uppermost Moscovian stage of the Pennsylvanian (Nel et al., 2013; Richards, 2013). As the upper boundary of the Moscovian is 307.0 Ma ± 0.1 Myr, this provides a minimum age of 306.9 Ma.

Soft maximum as for 75.3.

## 86. Crown Thysanoptera

This clade comprises Tubulifera and Terebrantia, their last common ancestor and all of its descendants (Fig. 21). Monophyly is established based on phylogenetic analysis of transcriptomes (Misof et al., 2014) and five nuclear protein-coding genes (Buckman et al., 2013). There is no adequate phylogenetic test of thysanopteran monophyly using morphology (reviewed by Mound and Morris, 2007).

### 86.1. *Fossil specimens*

*Fusithrips crassipes* Shmakov, 2009. PIN 3064/8547, holotype, preserving a female body (Fig. 22b).

### 86.2. *Phylogenetic justification*

*F. crassipes* has not been included in a phylogenetic analysis, however, Shmakov (2009) describes characters placing it in the crown of Thysanoptera. In particular, Rs1 and M2 as crossveins rather than oblique veins suggest membership in the family Aeolothripidae (Shmakov, 2009). Whether crown or stem Aeolothripidae, the family is a part of Terebrantia, indicating crown group membership in Thysanoptera.

### 86.3. *Age justification*

The fossil *F. crassipes* was found in Layer 31 on the left bank of the Vitim River, 3 km downstream from the mouth of the Baissa River, Buryatia, Transbaikalian Russia (Shmakov, 2009). The fossiliferous lacustrine deposits are part of the Zaza Formation, Baissa locality. The Zaza Formation was once assigned to the Valanginian, based on correlation of fossil insect species, however palynological data from the appearance of *Asteropollis* spores suggests a younger date (O’Reilly et al., 2015; Zherikhin et al., 1998). *A asteroides,* one of the earliest angiosperms, has a worldwide distribution during the Early and mid Cretaceous (Friis et al., 2005), and has been biostratigraphically assigned to the Barremian-Aptian (Friis et al., 1999; Zherikhin et al., 1998). This range has since been completely revised to Albian-Campanian, on the basis of pollen morphology (Doyle and Endress, 2014) and younger appearances of *Asteropollis* (Dettmann and Thomson, 1987; Eklund et al., 2004; Martinez et al., 2013). A minimum age can thus be estimated by the upper bound of the Campanian, 72.1 Ma ± 0.2 Myr, or 71.9 Ma.

Soft maximum as for 75.3.

### 86.4. *Discussion*

*Triassothrips virginicus* Grimaldi et al., 2004 was described from the Late Triassic of Virginia. Although it was used as a calibration for Thysanoptera by Misof et al. (2014), *T. virginicus* is found in a basal polytomy outside of any crown group members in a morphological phylogeny (P. Nel et al., 2012). *Liassothrips crassipes* Martynov, 1927 is known from the Late Jurassic of Kazakhstan, but is classified in an extinct family, Liassothripidae (Shmakov, 2008). While Shmakov (2008) suggests Liassothripidae is the oldest family in the Tubulifera, making it crown Thysanoptera, characters are also listed linking it with Terebrantia; without a phylogenetic analysis it is difficult to assess their polarity and thus crown affinity.

## 87. Crown Hemiptera

This clade comprises Sternorrhyncha (aphids, scale insects), Fulgoromorpha (planthoppers), Cicadomorpha (cicadas, leafhoppers, treehoppers), Coleorrhyncha (moss bugs) and Heteroptera (typical bugs), their last common ancestor and all of its descendants (Fig. 21). Monophyly of Hemiptera is established by phylogenetic analysis of housekeeping genes (Cryan and Urban, 2012), whole mitochondrial genomes (Cui et al., 2013), transcriptomes (Misof et al., 2014; Simon et al., 2012, and wing morphology (Yoshizawa and Saigusa, 2001).

### 87.1. *Fossil specimens*

As for 85.1.

### 87.2. *Phylogenetic justification*

As for 85.2.

### 87.3. *Age justification*

As for 85.3.

## 88. Crown Psocodea

This clade comprises Psocoptera (barklice) and Phthiraptera (true lice), their last common ancestor and all of its descendants (Fig. 21). Monophyly of this clade is supported by phylogenies of transcriptomes (Misof et al., 2014), nuclear protein-coding genes (Ishiwata et al., 2011), mitochondrial genes (Yoshizawa and Johnson, 2003), and morphology (Lyal, 1985).

### 88.1. *Fossil specimens*

*Cretoscelis burmitica* Grimaldi and Engel, 2006. AMNH Bu912, holotype, female in amber.

### 88.2. *Phylogenetic justification*

In a morphological phylogenetic analysis, *C. burmitica* was a stem group member of Liposcelididae (Grimaldi and Engel, 2006). Liposcelididae is an extant family of Psocoptera, thus within the crown group of Psocodea.

### 88.3. *Age justification*

As for 81.3.

## 89. Crown Holometabola

This clade comprises Hymenoptera (sawflies, ants, bees, wasps) and Aparaglossata, their last common ancestor and all of its descendants (Figs. 23 and 24). Support for monophyly comes from phylogenetic analysis of transcriptomes (Misof et al., 2014; Peters et al., 2014), morphology (Beutel et al., 2011), and morphology plus molecules (Oakley et al., 2013). This clade exists in GenBank, but as Endopterygota.

**Fig. 23.**
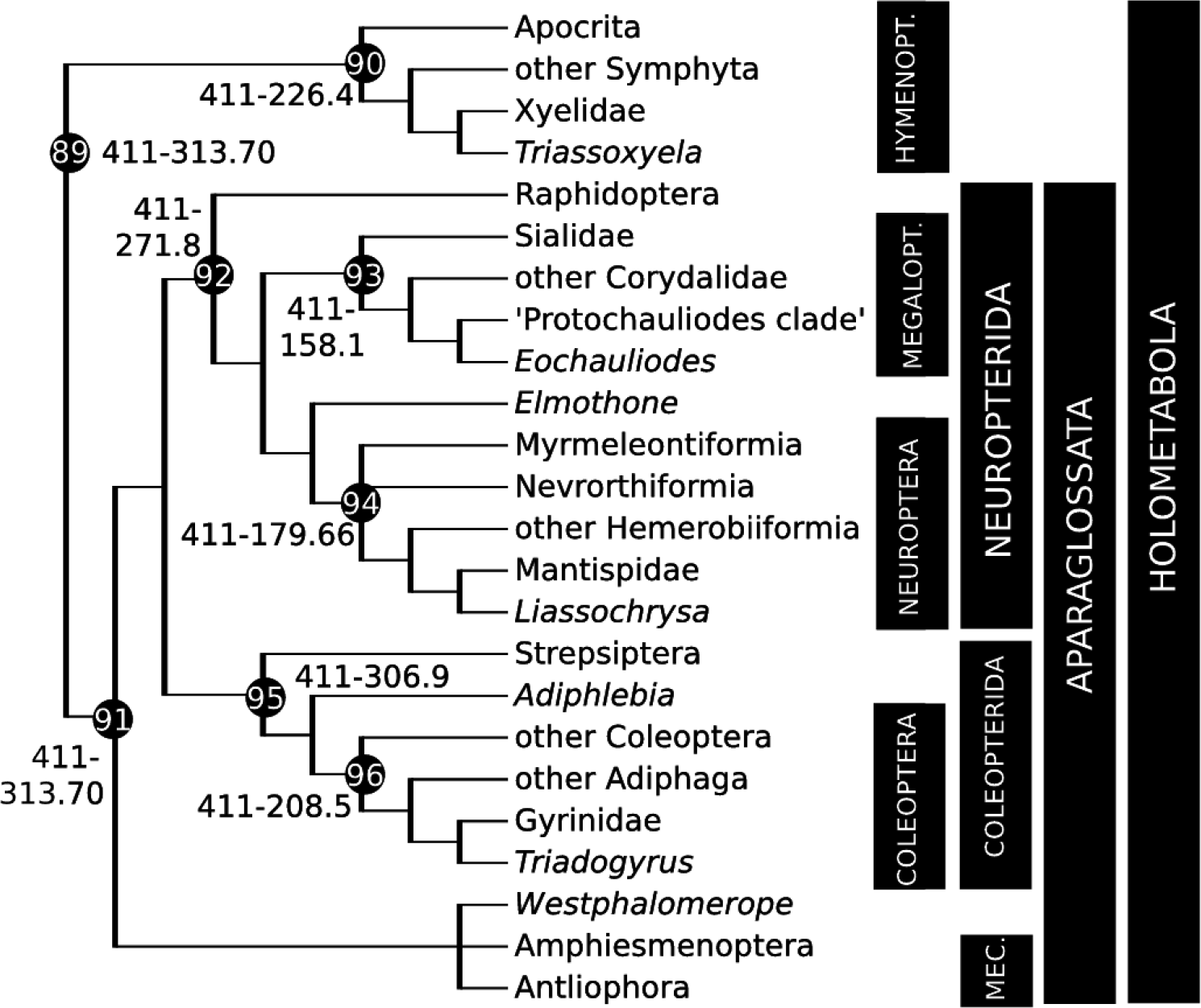
Calibration diagram for Hymenoptera, Neuropterida, and Coleopterida (nodes 89–96). Hymenopt. = Hymenoptera, Mec. = Mecopterida, Megalopt. = Megaloptera.

**Fig. 24.**
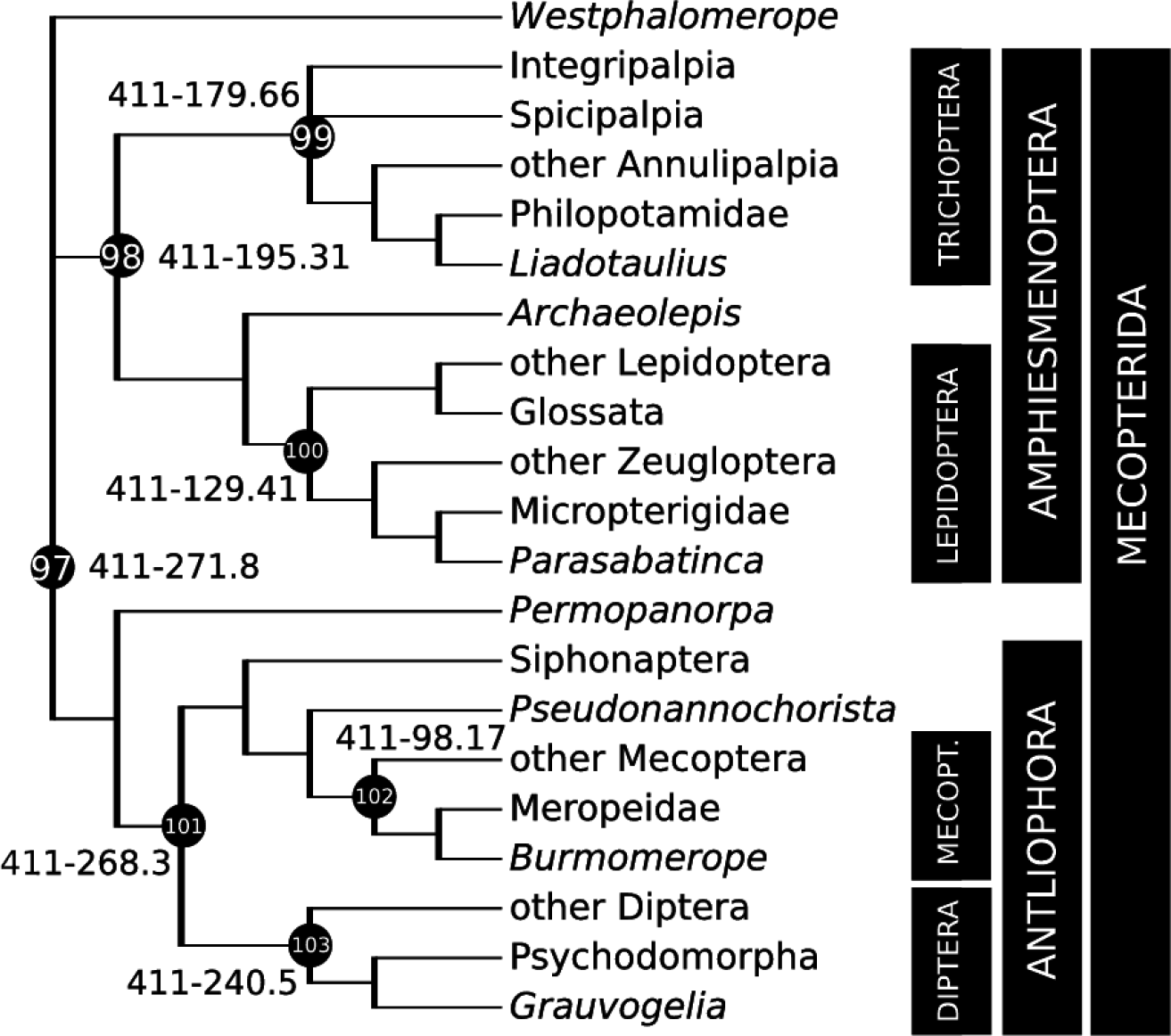
Calibration diagram for Mecopterida (nodes 97–103). Mecopt. = Mecoptera.

### 89.1. *Fossil specimens*

*Westphalomerope maryvonneae* Nel et al., 2007. MNHN-LP-R.55181, holotype, preserving a hindwing. Pictured in Nel et al. (2007; Fig. 1a).

### 89.2. *Phylogenetic justification*

*W. maryvonneae* has not been included in a phylogenetic analysis, nor have any other members of its family, Protomeropidae. Mostly Permian members of Protomeropidae have been proposed to have affinities with a variety of holometabolan clades, including the total groups of Trichoptera, Mecoptera, and more generally Amphiesmenoptera or Antliophora (Grimaldi and Engel, 2005; Kukalova-Peck and Willmann, 1990; Morse, 1997; Nel et al., 2013, 2007; Sukatsheva et al., 2007). Crown amphiesmenopteran (and thus trichopteran) affinity may be unlikely, as Protomeropidae lack a key synapomorphy, a true ’;double-Y loop’ arrangement of the anal veins on the forewing (Labandeira, 2011; Minet et al., 2010). However, Permian Protomeropidae possess Carpenter’;s organs in the male, a probable apomorphy for total group Mecoptera (Minet et al., 2010). Protomeropidae (with a younger date) was subsequently used to calibrate the basal split of Mecopterida for divergence time estimation (Rehm et al., 2011). Pending phylogenetic analysis of wing morphology, it is difficult to assign a specific placement for Protomeropidae, however, even with a conservative view all these possibilities are within crown Aparaglossata, and therefore crown Holometabola.

### 89.3. *Age justification*

*W. maryvonneae* was collected from black shales in the “Terril no. 5” horizon at the “Faisceau de Modeste”, “Veine Maroc” locality in Bruay-la-Bussiere, Pas-de-Calais, France (Nel et al., 2007). The locality is dated as early Langsettian (Nel et al., 2007), equivalent to the Westphalian A stage (Pointon et al., 2012). There is a SHRIMP U-Pb date within the middle Langsettian estimated at 317.63 Ma ± 0.12 Myr, however, the stratigraphy of Bruay-la-Bussiere is not precise enough to determine when in the Westphalian A the fossil occurred (Nel et al., 2007; Pointon et al., 2012). Therefore, we use a date inclusive of the upper boundary of the Westphalian A, which is the upper boundary of Westphalian B. U-Pb dating of zircons constrains the upper boundary of the Westphalian B to 313.78 Ma ± 0.08 Myr (Pointon et al., 2012), so a minimum age for *W. maryvonneae* is 313.70 Ma.

Soft maximum as for 75.3.

### 89.4. *Discussion*

*Srokalarva berthei* Haug et al., 2015 is a putative holometabolan larva, from the Mazon Creek and thus younger than *W. maryvonneae. S. berthei* has been interpreted as both an antliophoran (Labandeira, 2011) and a neuropterid (Haug et al., 2015), both positions within the crown group of Holometabola. *S. berthei,* however, remains informative about the evolutionary timing of insect metamorphosis.

## 90. Crown Hymenoptera

This clade comprises Symphyta (sawflies) and Apocrita (itself comprising Apoidea, Formicidae, and a paraphyletic group of wasps), their last common ancestor and all of its descendants (Fig. 23). Monophyly is supported by phylogenetic analysis of transcriptomes (Misof et al., 2014), morphology (Beutel et al., 2011), and morphology analysed together with molecular data (Ronquist et al., 2012).

### 90.1. *Fossil specimens*

*Triassoxyela foveolata* Rasnitsyn, 1964. PIN 2070/1, holotype (Fig. 25a).

**Fig. 25.**
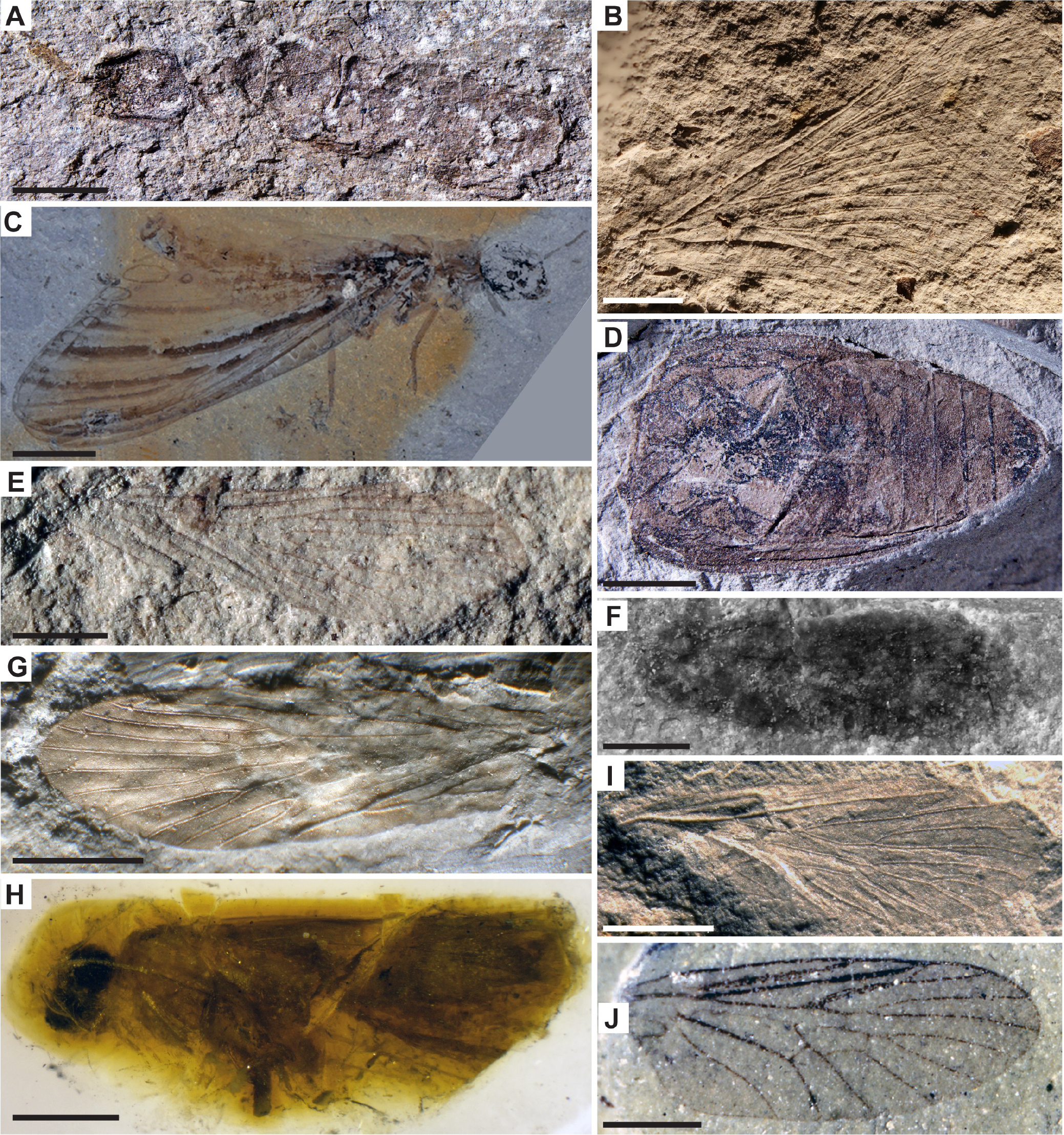
Holometabola fossil calibrations for (A) node 90: *Triassoxyela foveolata,* PIN 2070/1, scale bar 1mm, image credit A. Rasnitsyn; (B) node 92: *Elmothone martynovae,* MCZ 5585, scale bar 2mm, image credit Museum of Comparative Zoology, Harvard University (©President and Fellows of Harvard College); (C) node 93: *Eochauliodes striolatus,* CNU-MEG-NN2011004 P/C, scale bar 5mm, image credit D. Ren; (D) node 96: *Triadogyrus sternalis,* PIN 3320/13, scale bar 2mm, image credit A. Rasnitsyn; (E) node 97: *Permopanorpa inaequalis,* YPM IP 005058, scale bar 1mm, image credit J. Utrup; (F) node 98: *Archaeolepis mane,* BMNH In. 59397, scale bar 2mm, image credit NHMUK; (G) node 99: *Liadotaulius maior,* LGA 1995, scale bar 2mm, image credit J. Ansorge; (H) node 100: *Parasabatinca aftimacrai,* holotype, scale bar 2mm, image credit D. Azar; (I) node 101: *Pseudonannochorista willmanni,* PIN 966/21, scale bar 1mm, image credit A. Bashkuev; (J) node 103: *Grauvogelia arzvilleriana,* Louis Grauvogel collection No. 5514, scale bar 1mm, image credit L. Grauvogel-Stamm.

### 90.2. *Phylogenetic justification*

In the total-evidence phylogenetic analysis of Ronquist et al. (2012), *T. foveolata* was found as a stem group member of the family Xyelidae. As Xyelidae is a crown group family of sawflies, it is thus crown Hymenoptera.

### 90.3. *Age justification*

*T. foveolata* was found in the Madygen Formation, south of the Fergana Valley, Kyrgyzstan. Key plant fossils *Scytophyllum* and *Neocalamites* correlate the Madygen to the *Scytophyllum* flora of the upper Keuper lithographic unit, of Ladinian-Carnian age (Dobruskina, 1995). The *Scytophyllum* flora is correlated with the Cortaderitian Stage of Gondwana due to the abundance of *Scytophyllum* (Morel et al., 2003). The Cortaderitian Stage is divided into 3 Biozones; a 40Ar/39Ar radiometric date for the middle biozone of the Cortaderitian Stage of 228.5 Ma ± 0.3 Myr was measured by Rogers et al. (1993), falling within the Carnian (O’Reilly et al., 2015). The Gondwanan Puesto Viejo Formation, part of the Barrealian Stage underlying the Cortaderitian Stage (and therefore the *Scytophyllum* flora) has been radiometrically dated to 232 Ma ± 4 Myr, also within the Carnian (Valencio et al., 1975). Recently calculated SHRIMP U-Pb dates dispute this age, instead constraining the Puesto Viejo Formation to 235.8 Ma ± 2.0 Myr (Ottone et al., 2014). This suggests the Cortaderitian Stage is no older than 237.8 Ma, and therefore the *Scytophyllum* flora and Madygen Formation can be no older than this age, still within the Carnian. Note that a GSSP for the Carnian-Norian boundary has not yet been identified; radioisotopic ages may suggest a younger boundary at |220 Ma (Lucas et al., 2012). Nevertheless, a commonly accepted date for the Carnian upper boundary is estimated at 228.4 Ma ± 2 Myr based on cyclostratigraphy and a candidate GSSP (e.g. Ogg, 2012; Ogg et al., 2014), so this provides a minimum age at 226.4 Ma.

Soft maximum as for 75.3.

### 90.1 *Discussion*

Previous work has suggested *Archexyela* ipswichensis Engel, 2005 from the Mt. Crosby Formation, Ipswich Coal Measures of Queensland, Australia may be the oldest hymenopteran (e.g. Nicholson et al., 2015). However, the stratigraphy of the Ipswich Basin also provides a minimum age of uppermost Carnian (Purdy and Cranfield, 2013), thus we calibrate crown Hymenoptera with the equally old *T. foveolata,* which has been included in landmark total-evidence phylogenies (O’Reilly et al., 2015; Ronquist et al., 2012).

## 91. Crown Aparaglossata

This clade comprises Neuropterida, Coleopterida and Mecopterida, their last common ancestor and all of its descendants (Fig. 23). The clade was first named by Peters et al. (2014) based on support from phylogenetic analysis of transcriptomes.

### 91.1. *Fossil specimens*

As for 89.1.

### 91.2. *Phylogenetic justification*

As for 89.2.

### 91.3. *Age justification*

As for 89.3.

## 92. Crown Neuropterida

This clade comprises Raphidioptera (snakeflies), Megaloptera (fishflies), Neuroptera (net-winged insects), their last common ancestor and all of its descendants (Fig. 23). Monophyly is established by phylogenetic analysis of transcriptomes (Misof et al., 2014; Peters et al., 2014), protein-coding genes (Wiegmann et al., 2009), morphology (Beutel et al., 2011), and combined molecular and morphological data (Winterton et al., 2010).

### 92.1. *Fossil specimens*

*Elmothone martynovae* Carpenter, 1976. MCZ 5585, holotype, a forewing (Fig. 25b). Figured by Prokop et al. (2015; Fig. 7A).

### 92.2. *Phylogenetic justification*

A morphological phylogenetic analysis placed the Permithonidae *sensu lato* as a stem group to the extant Neuroptera (Ren et al., 2009; shown in supplementary information therein). A position on the stem lineage of Neuroptera is thus part of the crown group of Neuropterida. As the coding was done at a family level, we note with caution that monophyly of the Permithonidae and the exact relationships of its member species with Neuroptera has not been examined in a phylogenetic context and remain obscure (Prokop et al., 2015). Nonetheless, most members of Permithonidae possess the character states coded by Ren et al. (2009), so we take the oldest well-described member, *E. martynovae,* as a calibration fossil.

### 92.3. *Age justification*

This fossil occurs in the Carlton Limestone Member of the Wellington Formation in the Sumner Group of Elmo, Dickinson County, central Kansas (Prokop et al., 2015; Zambito et al., 2012). The insect-bearing locality is correlated with the Leonardian regional Stage (Sawin et al., 2008; Zambito et al., 2012) on the basis of conchostracan biostratigraphy (Tasch, 1962). The Leonardian spans the Artinskian and the younger Kungurian Stage (Henderson et al., 2012). The upper bound of the Kungurian is 272.3 Ma ± 0.5 Myr, thus providing a conservative minimum age estimate of 271.8 Ma. Soft maximum as for 75.3.

### 92.4. *Discussion*

A younger member of the Permithonidae, *Permithone belmontensis* Tillyard, 1922 (Changhsingian or latest Permian of Belmont, Australia), was previously used as a calibration fossil for Neuropterida (Misof et al., 2014).

## 93. Crown Megaloptera

This clade comprises Sialidae and Corydalidae, their last common ancestor and all of its descendants (Fig. 23). Monophyly is established by phylogenetic analysis of full mitochondrial genomes (Wang et al., 2012), transcriptomes (Misof et al., 2014), and morphology of the wing base (Zhao et al., 2014).

### 93.1. *Fossil specimens*

*Eochauliodes striolatus* Liu et al., 2012. CNU-MEG-NN2011004 P/C, holotype part/counterpart, laterally preserving an adult including forewing and hindwing (Fig. 25c).

### 93.2. *Phylogenetic justification*

*E. striolatus* was included in a phylogenetic analysis of morphology, encompassing characters from wing venation, adult genitalia, and larval morphology where possible (Liu et al., 2012). *E. striolatus* was found within the crown Corydalidae, at the base of the ‘Protochauliodes clade’ (comprising extant species). This position is therefore within crown Megaloptera. The bifurcated anterior branch of the Rs vein is a relevant synapomorphy (Liu et al., 2012).

### 93.3. *Age justification*

As for 79.3.

### 93.4. *Discussion*

Another fossil from the Daohugou fauna, *Jurochauliodes ponomarenkoi* Wang and Zhang, 2010 was found in a slightly more basal position within the Corydalidae (and thus Megaloptera) crown group (Liu et al., 2012). As it is of equal age to *E. striolatus,* it is also an acceptable calibration fossil for this clade.

## 94. Crown Neuroptera

This clade comprises Hemerobiiformia, Nevrorthiformia and Myrmeleontiformia, their last common ancestor and all of its descendants (Fig. 23). Monophyly is supported by phylogenetic analysis of ribosomal and mitochondrial genes (Haring and Aspöck, 2004), housekeeping genes (Wiegmann et al., 2009), and morphology (Beutel et al., 2011; Zhao et al., 2014).

### 94.1. *Fossil specimens*

*Liassochrysa stigmatica* Ansorge and Schluter, 1990. MBA.I 5046 (formerly from the Ansorge collection, Dobbertin, Germany: No. LDA301), holotype, preserving a forewing. Imaged in (Wedmann and Makarkin, 2007: Fig. 5a).

### 94.2. *Phylogenetic justification*

*L. stigmatica* was coded in the morphological matrix for the total evidence phylogeny of Liu et al. (2015). In that tree, it was a member of crown group Mantispidae, a family within Hemerobiiformia, and therefore crown Neuroptera (Liu et al., 2015). This fossil was also the oldest calibration used for Neuroptera in the divergence time analysis of Winterton et al. (2010).

### 94.3. *Age justification*

The fossil was recovered from the former clay pit of Schwinz, near Dobbertin, Mecklenburg, northeast Germany (Ansorge and Schluter, 1990). Multiple associated ammonites *(Eleganticeras elegantulum, Lobolytoceras siemensi, Harpoceras capellatum)* indicate that the locality is correlated to the lower part of the *H. falciferum* ammonite Zone, early Toarcian (Ansorge and Schluter, 1990; O’Reilly et al., 2015; Palfy et al., 2002, 2000). The Boreal *falciferum* Zone is equivalent to the Tethyan H. *serpentinum* ammonoid Zone (Macchioni, 2002), and succeeded by the *Hildoceras bifrons* ammonoid Zone. The base of the *bifrons* Zone has been dated to 180.36 Ma ± 0.7 Myr (Ogg et al., 2012b). From this, an upper bound of the *falciferum* Zone can be derived, and thus a minimum age for Dobbertin, at 179.66 Ma.

Soft maximum as for 75.3.

## 95. Crown Coleopterida

This clade comprises Strepsiptera (twisted wing parasites) and Coleoptera (beetles), their last common ancestor and all of its descendants (Fig. 23). Monophyly of this clade is an extremely well-examined question in systematics, often used to illustrate the principle of long branch attraction (e.g. Boussau et al., 2014; Carmean and Crespi, 1995; Huelsenbeck, 1998, 1997; Niehuis et al., 2012; Siddall and Whiting, 1999; Whiting et al., 1997; Wiegmann et al., 2009). However, recent analyses of genomes and transcriptomes have consistently converged on a sister group relationship between Strepsiptera and Coleoptera (Boussau et al., 2014; Misof et al., 2014; Niehuis et al., 2012).

### 95.1. *Fossil specimens*

*Adiphlebia lacoana* Scudder, 1885. USNM 38140, holotype, preserving a forewing. Imaged in Béthoux (2009: Fig. 1(3–5)).

### 95.2. *Phylogenetic justification*

A. *lacoana* has not been included in formal cladistic analysis to test its phylogenetic position. Variation in forewing venation within species provides a challenge to homology statements (Béthoux, 2009; Garwood and Sutton, 2010). However, *A lacoana* shares a forewing character with the crown beetle suborder Archostemata, specifically: “the areas between ScP, RA, branches of RP, M, branches of CuA, and AA filled with a regular network of quadrangular to pentagonal cells forming intercalary longitudinal pseudo-veins” (Béthoux, 2009). Intercalary pseudo-veins are also observed in fossils throughout Pterygota, suggesting this is either a symplesiomorphic or homoplastic state. However, some of the wing regions exhibiting intercalary veins (ScP and RA) are restricted to fossil Coleoptera (Béthoux, 2009). *A. lacoana* has thus been designated as a sister group to all crown Coleoptera, i.e. a member of its stem lineage (Béthoux, 2009), and thus a crown group member of Coleopterida. This assignment is also supported by putative larval material (Labandeira, 2011).

There has been debate, however, about the taphonomy of the network of wing veins, suggesting they are clumps of clay instead of morphological characters (Kukalova-Peck and Beutel, 2012). Alternative placements were on the stem lineage of Neuroptera or Neuropterida (Kukalova-Peck and Beutel, 2012), or outside Holometabola altogether (Guan et al., 2016; Nel et al., 2013).

### 95.3. *Age justification*

Minimum as for 31.3. Soft maximum as for 75.3.

## 96. Crown Coleoptera

This clade comprises Archostemata, Myxophaga, Adephaga (ground beetles, tiger beetles, diving beetles, whirligig beetles) and Polyphaga (ladybugs, longhorn beetles, weevils, fireflies, scarabs, stag beetles, rove beetles), their last common ancestor and all of its descendants (Fig. 23). Monophyly is supported by phylogenetic analysis of nuclear protein-coding genes (McKenna et al., 2015) and morphology (Beutel et al., 2011).

### 96.1. *Fossil specimens*

*Triadogyrus sternalis* Ponomarenko, 1977. PIN 3320/13, holotype (Fig. 25d), an exoskeleton without head, prothorax, or legs (Arnol’di et al., 1992).

### 96.2. *Phylogenetic justification*

In the morphological phylogenetic analysis of Beutel et al. (2013), *T. sternalis* is a stem group member of the family Gyrinidae. As Gyrinidae is part of Adephaga, crown membership within both Adephaga and Coleoptera is justified.

### 96.3. *Age justification*

*T. sternalis* was recovered from the mouth of Bereni River near Garazhokva settlement, Khar’kov oblast, Izyum region, Ukraine (Arnol’di et al., 1992). Fossiliferous strata are lacustrine claystone, part of the Protopivka Formation, estimated as Late Carnian-Early Norian age within the late Triassic (Arnol’di et al., 1992; Rasnitsyn and Quicke, 2002; Shcherbakov, 2008). The upper Norian boundary is estimated at ~208.5 Ma, so this provides a conservative minimum age.

Soft maximum as for 75.3.

### 96.4. *Discussion*

The Pennsylvanian (Mazon Creek) fossil *A. lacoana* is a member of the stem lineage of Coleoptera (Béthoux, 2009). Recent divergence time analyses (McKenna et al., 2015; Misof et al., 2014) have therefore elected not to calibrate the crown group of Coleoptera with *A lacoana.* Resulting age estimates for crown Coleoptera ranged from 250–280 Ma, substantially younger than *A lacoana* itself.

The Permian *Coleopsis archaica* Kirejtshuk et al., 2014 was noted as a stem group fossil, potentially Protocoleoptera (stem Coleoptera) (McKenna et al., 2015). Thus it was not used as a calibration. However, using younger internal calibrations (the oldest from Madygen, 225 Ma in their dataset), the crown origin of Coleoptera was estimated around 250 Ma (McKenna et al., 2015). Accounting for error bars, the crown group may have originated shortly before (or shortly after) the end-Permian extinction, timing that is compatible with *C. archaica* as a very early representative.

## 97. Crown Mecopterida

This clade comprises Amphiesmenoptera and Antliophora, their last common ancestor and all of its descendants (Fig. 24). Phylogenetic support for monophyly of Mecopterida comes from genomes (Niehuis et al., 2012), transcriptomes (Misof et al., 2014) and morphology (Beutel et al., 2011). Note that the clade we refer to as Mecopterida was called Panorpida by Grimaldi and Engel (2005), because they used Mecopterida to refer to paraphyletic Mecoptera containing Siphonaptera (and excluding Diptera). Our clade Mecopterida was also referred to as Panorpoidea by several sources, including Ren et al. (2009). Analyses relying on molecular data alone do not support the paraphyly of Mecoptera (Misof et al., 2014; Peters et al., 2014; Wiegmann et al., 2009), and these tend to utilize our conception of Mecopterida.

### 97.1. *Fossil specimens*

*Permopanorpa inaequalis* Tillyard, 1926. YPM IP 005058, holotype, a wing (Fig. 25e).

### 97.2. *Phylogenetic justification*

*P. inaequalis* is the oldest member of the family Permopanorpidae, which was coded (at the family level) in the morphological matrix of Ren et al. (2009). In that tree, it was part of the total group of Antliophora (although the figure label indicated it was inside crown Antliophora, the topology clearly indicates it was on the stem lineage; Ren et al., 2009). As a stem group member of Antliophora, *P. inaequalis* is thus crown group Mecopterida.

### 97.3. *Age justification*

As for 92.3.

### 97.4. *Discussion*

*W. maryvonneae* (family Protomeropidae), from the Bashkirian of France, has been proposed as an “amphiesmenopteran or antliophoran”, which may mean it belongs on the stem lineage of one of those clades and thus in crown Mecopterida. It has also been drawn onto a phylogram as a stem member of Mecoptera (Nel et al., 2013). However, the evidence for any particular placement within Mecopterida is ambiguous, and thus it is possible the fossil is only stem Mecopterida. See 89.2 for greater detail.

Conversely, the early to middle Permian family Kaltanidae has been included in a phylogenetic analysis, and shown to fall on the stem lineage of Amphiesmenoptera (Ren et al., 2009). Although Kaltanidae have been previously discussed as calibration fossils for Mecopterida (e.g. Benton and Donoghue, 2007), their oldest members are from the Kungurian Stage of Russia, the same age as Elmo (Bashkuev, 2008). Additionally, the family Belmontiidae is within crown Mecopterida in a phylogenetic analysis (Ren et al., 2009), but its oldest fossil is from the late Permian (Changhsingian) of Australia (Grimaldi and Engel, 2005).

## 98. Crown Amphiesmenoptera

This clade comprises Trichoptera (caddisflies) and Lepidoptera (butterflies, moths), their last common ancestor and all of its descendants (Fig. 24). Monophyly is supported by phylogenetic analysis of transcriptomes (Misof et al., 2014), nuclear protein-coding genes (Wiegmann et al., 2009), and morphology (Beutel et al., 2011; Kristensen, 1991).

### 98.1. *Fossil specimens*

*Archaeolepis mane* Whalley, 1985. BMNH In. 59397, holotype, preserving a hindwing (Fig. 25f).

### 98.2. *Phylogenetic justification*

*A. mane* has been used to calibrate the lepidopteran root in previous molecular dating analyses (e.g. Wahlberg et al., 2013), where this relationship is based on two lines of evidence: wing scales and wing venation. The preservation of extremely rare scales completely covering the hindwing excludes *A mane* from crown Trichoptera (Whalley, 1986, 1985) because Trichoptera only bear scales on the forewing. The presence of scales across the entirety of a wing with panorpoid venation can only signal amphiesmenopteran affinity.

The Sc vein (with one visible branch) of *A. mane* is unlikely to represent the ancestral state for Lepidoptera, as the number of Sc vein branches varies in early-diverging moths, but is usually two branches, and this vein is multi-branched ancestrally in Amphiesmenoptera (Kukalova-Peck and Willmann, 1990; Minet et al., 2010; Sukatsheva and Vassilenko, 2011; Schachat and Brown, 2016). Although Trichoptera tend to have more wing veins than Lepidoptera, it is likely that fewer veins were lost by ancestral moths than is currently accepted in the literature, making distinctions between amphiesmenopteran branches difficult for Mesozoic fossils (Schachat and Brown, 2015, 2016). As reconstructed by Whalley (1986), the wing venation of *A. mane* differs markedly from the venation of crown Lepidoptera: there is a single, branched Cubitus vein, with CuP apparently absent, and all three branches of the anal vein reach the dorsum. In contrast, basal crown Lepidoptera nearly always have a CuP vein, and the anal vein becomes fused before reaching the dorsum (Common, 1973; Schachat and Brown, 2016). A position for *A mane* within the crown group of Lepidoptera thus cannot be confirmed, but an identity as stem Lepidoptera is highly likely based on the wing scales in particular, in agreement with Whalley (1986). This fossil would therefore be within the crown group of Amphiesmenoptera.

### 98.3. *Age justification*

Whalley (1985) noted the locality as Black Ven, Charmouth, Dorset, on the Jurassic Coast of England. This was further specified as calcareous flatstone, probably from Bed 75a, of the *Caenisitesturneri* ammonoid Zone (Sohn et al., 2012), or *Microderoceras birchi* Nodular of the “Shales with Beef” (Lang et al., 1923). Chemostratigraphy places the *C. turneri* (and *M. birchi)* Zones within the middle Sinemurian (Jenkyns et al., 2002). The younger bound of the *C. turneri* Zone is thus dated to 195.31 Ma (Ogg et al., 2012b), providing a minimum age.

Soft maximum as for 75.3.

## 99. Crown Trichoptera

This clade comprises ‘Spicipalpia’, Annulipalpia and Integripalpia their last common ancestor and all of its descendants (Fig. 24). Monophyly is supported by phylogenetic analysis of transcriptomes (Misof et al., 2014), housekeeping genes (Kjer et al., 2002, 2001; Malm et al., 2013), and morphology (Beutel et al., 2011).

### 99.1. *Fossil specimens*

*Liadotaulius maior* Handlirsch, 1906. Handlirsch (1906) did not designate a holotype, thus we refer to three specimens figured by Ansorge (2002): (Ansorge collection, to be deposited at MBA: LGA 1995; Fig. 25g), female forewing, (LGA 672), male forewing, and (LGA 1710), hindwing.

### 99.2. *Phylogenetic justification*

Taxonomic placement of *L. maior* requires apomorphies from the male wings, as female wings have many plesiomorphic venation characters (Ansorge, 2002). Further studies of a younger congeneric, *L. daohugouensis* Wu and Huang, 2012, reveal new characters shared with crown group Trichoptera. These include the apical part of Cu2 bending towards the wing margin, its desclerotisation, and complete anal veins (Ansorge, 2002; Wu and Huang, 2012). These apomorphies place the genus *Liadotaulius* in Philopotamidae, a family within crown group Annulipalpa, and thus Trichoptera (Wu and Huang, 2012).

### 99.3. *Age justification*

The fossils were recovered from Grimmen, northeast Germany (Ansorge, 2002). Further age information as for 94.3.

### 99.4. *Discussion*

Possible caddisfly larval cases were recently described from the early Permian of Brazil (Mouro et al., 2016). The cases are tubular in form, and particulate matter is stuck to some of the cases (Mouro et al., 2016). Of the two most basal extant trichopteran families that construct larval cases, the Hydroptilidae construct tubular cases and the Glossosomatidae construct their cases from particulate matter (Malm et al., 2013). However, various types of larval cases, including tubular cases, are constructed by caddisflies belonging to distantly-related lineages (Malm et al., 2013); at present, it cannot be assumed that tubular larval cases have originated only once within the Trichoptera. Assuming the Permian fossils are indeed insect larval cases, therefore, does not confirm the phylogenetic position of their inhabitants within crown versus stem Trichoptera, as larval cases might have evolved in the stem group.

## 100. Crown Lepidoptera

This clade comprises Zeugloptera, Aglossata, Heterobathmiina and Glossata (itself comprising six infraorders, over 100 families, and over 160,000 species), their last common ancestor and all of its descendants (Fig. 24). The list of lepidopteran subclades comes from a recently published consensus (van Nieukerken et al., 2011), although some molecular analyses have not recovered these clades (Regier et al., 2013). Monophyly is supported by nuclear protein-coding genes (Regier et al., 2013), transcriptomes (Misof et al., 2014), and morphology (Beutel et al., 2011).

### 100.1. *Fossil specimens*

*Parasabatinca aftimacrai* Whalley, 1978. Although the holotype (the specimen to which we refer; Fig. 25h) and two paratypes were attributed to the NHMUK (Sohn et al., 2012), they are now housed as part of the Acra collection, curated by D. Azar, pending construction of a fossil museum at the Lebanese University in Fanar (Matn), Lebanon (D. Azar, pers. comm.).

### 100.2. *Phylogenetic justification*

Although not included in formal phylogenetic analysis, morphological characters (morphology of the antennae with ascoids, number and shape of tibial spurs, mouthpart and maxillary palp morphology) support the position of *P. aftimacrai* in what was formerly known as the ‘*Sabatinca’* group of genera within Micropterigidae (Kristensen and Skalski, 1998; D. Azar, pers. comm.). As *P. aftimacrai* has an unbranched R vein and because Rs4 terminates below the apex, the wing venation of *P. aftimacrai* most closely resembles that of the extant species *Austromartyria porphyrodes,* which belongs to what is now known as the ‘Southern sabatincoid’ lineage of Micropterigidae (Gibbs, 2010). Based on the above morphological characters, *P. aftimacrai* is supported as the oldest crown group member in multiple summaries of the fossil record of Lepidoptera (Grimaldi and Engel, 2005, Fig. 13.13; Sohn et al., 2015, Fig. 2). Membership either within the crown or stem of Micropterigidae places the fossil within crown Zeugloptera and crown Lepidoptera.

### 100.3. *Age justification*

As for 76.3.

### 100.4. *Discussion*

A number of older fossils have been assigned to Lepidoptera, but their position within the crown is ambiguous. *A. mane*, from the Early Jurassic of Dorset, England, has been used to calibrate the lepidopteran root in previous molecular dating analyses (e.g. Wahlberg et al., 2013). See 98.2 for exclusion of *A. mane* from the lepidopteran crown.

Sohn et al. (2015) suggested that two undescribed fossil species are members of the Micropterigidae stem lineage (and thus crown Zeugloptera and crown Lepidoptera). These are from the Toarcian (Early Jurassic) Grüne Series of Grimmen, Germany (Ansorge, 2002), but they are too poorly characterized to be confidently used as calibration fossils.

## 101. Crown Antliophora

This clade comprises Siphonaptera (fleas), Mecoptera (scorpionflies) and Diptera (flies), their last common ancestor and all of its descendants (Fig. 24). Support for monophyly comes from phylogenetic analysis of transcriptomes (Misof et al., 2014; Peters et al., 2014) and morphology (Beutel et al., 2011). A recent analysis of mitochondrial genomes finds Amphiesmenoptera within Antliophora under some analytical conditions, potentially challenging antliophoran monophyly (Song et al., 2016).

### 101.1. *Fossil specimens*

*Pseudonannochorista willmanni* Novokshonov, 1994. PIN 966/21, holotype, preserving a forewing (Fig. 25i).

### 101.2. *Phylogenetic justification*

Pseudonannochoristinae is a subfamily of the Permochoristidae (Bashkuev, 2011; Novokshonov, 1994), however, a morphological phylogenetic analysis indicates polyphyly of this family (Ren et al., 2009). While the Permochoristinae (the other subfamily of Permochoristidae) fall on the stem lineage of Antliophora, Pseudonannochoristinae are part of the stem lineage of Mecoptera (Ren et al., 2009). Therefore, *P. willmanni,* as a member of Pseudonannochoristinae, is part of the crown group of Antliophora.

### 101.3. *Age justification*

Minimum as for 77.3. Soft maximum as for 75.3.

### 101.4. *Discussion*

Other fossils from the families Permotanyderidae, Permotipulidae, and Nannochoristinae are all demonstrably within crown group Antliophora (Ren et al., 2009). However, the localities from which they are known are younger, of Changhsingian age (Belmont, Australia and Mooi River, South Africa).

Fossils of Siphonaptera from the Jurassic (Gao et al., 2012) and Cretaceous (Gao et al., 2014; Huang, 2015), suggested to parasitise dinosaurs, have been excluded from the crown group in a recent molecular phylogenetic analysis (Zhu et al., 2015). The molecular clock analysis (which was calibrated with better established fossils from Dominican and Baltic amber, of Miocene and Eocene age, respectively) estimated the origins of crown Siphonaptera in the Valanginian to Aptian stages of the Early Cretaceous (Zhu et al., 2015). Furthermore, none of the Mesozoic fossils possess a diagnostic character combination for the crown group (Dittmar et al., 2016). As siphonapteran parasites of dinosaurs would require a strong assumption that piercing mouthparts must be used for blood-feeding (Dittmar et al., 2016), we do not include any crown fossil calibrations.

## 102. Crown Mecoptera

This clade comprises Apteropanorpidae, Bittacidae, Boreidae, Choristidae, Eomeropidae, Meropeidae, Nannochoristidae, Panorpidae, and Panorpodidae, their last common ancestor and all of its descendants (Fig. 24). Paraphyly of the traditional concept of Mecoptera (i.e. inclusion of Siphonaptera) was proposed by Whiting (2002) on the basis of four nuclear genes. This was, however, contradicted by analysis of six nuclear genes (Wiegmann et al., 2009), transcriptomes (Misof et al., 2014; Peters et al., 2014), and morphology (Beutel et al., 2011). Each of the latter studies recovered a monophyletic Mecoptera as the sister group of Siphonaptera.

### 102.1. *Fossil specimens*

*Burmomerope eureka* Grimaldi and Engel, 2013 (collection of James Zigras, available for study at AMNH: JZC Bu-84), holotype in amber.

### 102.2. *Phylogenetic justification*

*B. eureka* was assigned to the stem group of the extant family Meropeidae, based on several wing venation characters, including loss of slanted portion of CuA in forewing and R-Rs fork basal (Grimaldi and Engel, 2013). A position on the stem lineage of Meropeidae is therefore within the crown group of Mecoptera.

### 102.3. *Age justification*

As for 87.3.

## 103. Crown Diptera

This clade comprises Tipulomorpha (crane flies), Psychodomorpha (moth flies, sand flies), Culicomorpha (mosquitoes, black flies), Bibionomorpha (march flies, fungus gnats) and Brachycera (horse flies, robber flies, fruit flies, house flies, blow flies, etc.), their last common ancestor and all of its descendants (Fig. 24). Monophyly of Diptera has been supported in many phylogenetic analyses, including those based on transcriptomes (Misof et al., 2014), mitochondrial genomes and microRNA presence (Wiegmann et al., 2011), and morphology (Beutel et al., 2011; Wiegmann et al., 2011).

### 103.1. *Fossil specimens*

*Grauvogelia arzvilleriana* Krzeminski et al., 1994 (part of the private Louis Grauvogel collection, Ringendorf, Bas-Rhin, France, No. 5514), holotype (Fig. 25j).

### 103.2. *Phylogenetic justification*

*G. arzvilleriana* was included in the morphological cladistic analysis of Blagoderov et al. (2007). In that analysis, it was supported on the stem group of Psychodomorpha (Blagoderov et al., 2007). Although the original description assigned *G. arzvilleriana* to its own family, several wing venation characters were noted as similarities with families of Psychodomorpha (Krzeminski et al., 1994). The stem group of Psychodomorpha is within crown Diptera.

### 103.3. *Age justification*

Minimum as for 72.3. Soft maximum as for 75.3.

### 103.4. *Discussion*

A number of other dipteran fossils have been described from Grès à Voltzia; of special interest is *Voltziapupa tentata* Lukashevich et al., 2010, known only from well-preserved pupae, while *G. arzvilleriana* is known only from adult wings. The venation of the wing sheath in *V. tentata* is not well preserved, preventing a clear link of characters with *G. arzvilleriana* (Lukashevich et al., 2010).

## 104. Conclusions

Based on this compilation, qualitative trends in the fossil record of arthropod crown group members can be described. Patchiness in our taxonomic coverage results from differential preservation potential, with a notable scarcity of taxa living in habitats with poorly preserved facies (e.g. intertidal barnacles or pelagic copepods). More completely preserved fossils representing ancient divergences are well represented in our list of calibrations, perhaps owing to the attention devoted to phylogenetic analysis of deep fossil relationships within arthropods (e.g. Garwood and Dunlop, 2014; Lee et al., 2013; Legg et al., 2013; Oakley et al., 2013; Rota-Stabelli et al., 2011). However because our review is focused on crown groups, about half of our calibrations are from mid-Palaeozoic localities, making them much more numerous than those from Cambrian Burgess Shale-type biotas. Throughout the dataset, there is less of a relationship between ‘hard part’ preservation (calcified body parts, such as ostracod carapaces) and phylogenetic accuracy than might be expected.

Particular Konservat-Lagerstätten, such as Herefordshire (Silurian), Rhynie Chert (Devonian), Mazon Creek (Carboniferous), Crato (Cretaceous) and Burmese amber (Cretaceous), provide several calibrations representing different arthropod clades, presumably owing to their preservation of nonbiomineralised tissues required to identify arthropods to the ordinal level. These sites are critically important for the arthropod fossil record because of their relatively low preservation potential of many groups, as is particularly the case for terrestrial arthropods, being less well mineralised than many of the marine groups. This is reflected in the low level of congruence between the order of appearance of lineages in the fossil record (stratigraphic appearance) and the order of phylogenetic branching (Wills, 2001; O’Connor and Wills, 2016) in arthropods, as compared to more congruent datasets such as tetrapods (Benton et al., 1999, 2000; Norell and Novacek, 1992). Clustering of calibrations at Konservat-Lagerstätten localities may lead to highly variable lengths of ghost lineages for the different taxa that are preserved together at these sites, and indeed many of the clades in our database have soft maxima that are substantially older than their hard minimum date. Konservat-Lagerstätten localities are, however, fairly regularly spaced throughout the Middle to Upper Palaeozoic and the Mesozoic, and interim periods of time are punctuated by numerous other fossil localities yielding fewer calibration points. This results in our database having good coverage throughout the Phanaerozoic with fossil localities occurring on average every 4–10 million years. A notable exception is the 43.2 million year gap in the Ordovician, during which no definite earliest appearances of any crown-group orders have been identified in our study, despite this period being known as the Ordovician biodiversification event (Servais et al., 2008, Servais et al., 2010). Numerous arthropod stem lineage taxa were abundant during the Ordovician (e.g. trilobites), while the possible crown group taxa that have been described, e.g. ostracods (Siveter et al., 2014; Williams et al., 2008), barnacles (Van Roy et al., 2010), pycnogonids (Rudkin et al., 2013), xiphosurans (Lamsdell, 2013; Rudkin et al., 2008; Van Roy et al., 2010), and acariform mites (Bernini et al., 2002) do not meet the rigorous standards employed herein for determining calibration points. From the Late Devonian through the Mississippian (382.7 to 323.2 Ma) our dataset has another large gap during which we have only one calibration point, which interestingly corresponds with one of the largest mass extinctions events known in the fossil record (McGhee, 2013).

The field of divergence time estimation itself is rapidly advancing. New methodologies to incorporate fossil morphology and stratigraphy into the model of diversification (Heath et al., 2014; Wilkinson et al., 2011) and the phylogenetic topology itself (’;tip-dating’; Ronquist et al., 2012) are growing in popularity. Precision and accuracy of date estimates are improved with the inclusion of as many *a priori* justified fossils as possible in tip-(Lee et al., 2013; Ronquist et al., 2012; Zhang et al., or node-dating studies (Heath et al., 2014; Ho and Phillips, 2009; Warnock et al., 2012, 2015; Yang and Rannala, 2006). When examined in the context of geological and evolutionary history, the distribution of fossil calibrations in our dataset are comprehensive, and have been rigorously vetted to ensure they meet *a priori* requirements (Parham et al., 2012; Warnock et al., 2015). Following their use in molecular clock analyses, *a posteriori* methods such as cross-validation could be employed to explore the impact of calibrations on the resulting divergence time estimates (e.g. Battistuzzi et al., 2015; Dornburg et al., 2011;Marshall, 2008; Near et al., 2005), although these methods cannot justify removal of individual constraints (Warnock et al., 2015). Fossils mentioned in Discussion sections occupy key positions along clade stems, and should also be considered in divergence time applications. To this end, we have compiled a robust list of over 100 fossil calibrations covering much of the arthropod Tree of Life. We hope this summary will inspire further work clarifying the phylogenetic relationships of fossil arthropods, and morphological studies of characters linking them to their crown clades.

## ACKNOWLEDGEMENTS

We thank colleagues who assisted with tracking down and photographing fossils: S. Ahyong, L. Anderson, J. Ansorge, D. Azar, A. Bashkuev, G. Bechly, G. Boxshall, C. Brauckmann, D. Briggs, S. Butts, J. Collette, M. Coyne, J. Dunlop, M. Engel, R. Feldmann, S. Fouche, D. Garria-Bellido, L. Grauvogel-Stamm, T. Hegna, H. Henderickx, D. Huang, B. Hussaini, W. Jones, A. Kotov, J. Lamsdell, G. Li, C. Mellish, A. Nel, E. Olempska, R. Perez de la Fuente, G. Poinar, A. Rasnitsyn, D. Ren, A. Ross, D. Rudkin, A. Sanchez-Garcia, P. Selden, W. Shear, Y. Shen, N. Sinitshenkova, D. Siveter, D. Siveter, M. Sutton, J. Utrup, J. Waddington, D. Waloszek, J. Wittry, X. Zhang, and Y. Zhen. We acknowledge additional discussions and comments on fossil calibrations from J. Antcliffe, T. Astrop, O. Béthoux, A. Gale, R. Garwood, C. Haug, J. Luque, S. Maksoud, V. McCoy, D. Nicholson, T. Oakley, J. Ortega-Hernandez, J. Pineda, D. Pisani, O. Rota-Stabelli, S. Schachat, B. Wang, and two anonymous reviewers. We thank APSOMA for access to literature. JMW thanks G. Fournier and the Simons Foundation Collaboration on the Origin of Life #339603 for support. ACD and DAL are funded by the Oxford University Museum of Natural History. ACD also acknowledges funding from ZOONET Marie Curie Research Training network.

